# Blind exploration of the unreferenced transcriptome reveals novel RNAs for prostate cancer diagnosis

**DOI:** 10.1101/644104

**Authors:** M. Pinskaya, Z. Saci, M. Gallopin, N. H. Nguyen, M. Gabriel, V. Firlej, M. Descrimes, A. de la Taille, A. Londoño-Vallejo, Y. Allory, D. Gautheret, A. Morillon

**Author notes:** equal contribution (M. Pinskaya, Z. Saci). Institute for Research in Immunology and Cancer, Université de Montréal, H3T 1J4, Canada (Z. Saci). Correspondence to: Antonin Morillon,; Daniel Gautheret,.

## Abstract

The broad use of RNA-sequencing technologies held a promise of improved diagnostic tools based on comprehensive transcript sets. However, mining human transcriptome data for disease biomarkers in clinical specimens is restricted by the limited power of conventional reference-based protocols relying on uniquely mapped reads and transcript annotations. Here, we implemented a blind reference-free computational protocol, DE-kupl, to directly infer RNA variations of any origin, including yet unreferenced RNAs, from high coverage total stranded RNA-sequencing datasets of tissue origin. As a bench test, this protocol was powered for detection of RNA subsequences embedded into unannotated putative long noncoding (lnc)RNAs expressed in prostate cancer tissues. Through filtering and visual inspection of 1,179 candidates, we defined 21 lncRNA probes that were further validated for robust tumor-specific expression by NanoString single molecule-based RNA measurements in 144 tissue specimens. Predictive modeling yielded a restricted probe panel enabling over 90% of true positive detection of cancer in an independent dataset from The Cancer Genome Atlas. Remarkably, this clinical signature made of only 9 unannotated lncRNAs largely outperformed PCA3, the only RNA biomarker approved by the Food and Drug Administration agency, specifically, in detection of high-risk prostate tumors. The proposed reference-free computational workflow is modular, highly sensitive and robust and can be applied to any pathology and any clinical application.

## INTRODUCTION

RNA sequencing (RNA-seq) has revolutionized our knowledge of human transcriptome and has been implemented as a pivot technique in clinical applications for the discovery of RNA-based biomarkers allowing disease diagnosis, prognosis and therapy follow-up. However, most biomarker discovery pipelines are blind to uncharacterized RNA molecules since they rely on the alignment of uniquely mapped reads to annotated references of the human transcriptome which are far from complete (Uszczynska-Ratajczak et al. 2018), (Deveson et al. 2018), (Morillon and Gautheret 2019). Indeed, state-specific unspliced variants, rare mRNA isoforms, RNA hybrids originating from trans-splicing or genome rearrangements, unannotated intergenic or antisense noncoding RNAs, mobile elements or viral genome insertions would be systematically missed. A recent approach to RNA-seq data analysis, DE-kupl, combines k-mer (subsequences of fixed size) decomposition and differential expression analysis to discover transcript variations yet unreferenced in the human transcriptome (Audoux et al. 2017). Applied to poly(A)+ RNA-seq datasets of *in vitro* cell system, DE-kupl unveiled a large number of RNA subsequences embedded into novel long noncoding RNAs. These transcripts of more than 200 nucleotides in length transcribed by RNA polymerase II from intergenic, intronic or antisense noncoding genomic locations constitute a prevalent class of human genes. Some lncRNAs are now recognized as precisely regulated stand-alone molecules participating in the control of fundamental cellular processes (Jarroux et al. 2017), (Quinn and Chang 2015). They show aberrant and specific expression in various cancers and other diseases promoting them as biomarkers, therapeutic molecules and drug targets (Leucci 2018), (Van Grembergen et al. 2016). Importantly, some lncRNAs can be robustly detected in biological fluids (blood and urine) as circulating molecules or encapsulated into extracellular vesicles, hence, raising an attractive possibility of lncRNA biomarkers usage in non-invasive clinical tests (Silva et al. 2015), (Deng et al. 2017), (Wang et al. 2014), (Zhao et al. 2018a), (Wang et al. 2018). The only example of an RNA-based biomarker so far introduced in clinical practice is the PCA3 lncRNA in prostate cancer (PCa) (de Kok et al. 2002). PCA3 is transcribed antisense to the tumor suppressor *PRUNE2* gene and promotes its pre-messenger RNA editing and degradation (Salameh et al. 2015). Being overexpressed in 95% of PCa cases, PCA3 is detected in urines and helps diagnosis providing, in addition to other clinical tests, more accurate metrics regarding repeated biopsies (Groskopf et al. 2006), (Galasso et al. 2010). However, it remains inaccurate in discrimination between low- and high-risk tumors since its expression may dramatically decrease in aggressive PCa cases tempering its systematic usage (Loeb and Partin 2011), (Alshalalfa et al. 2017).

Since PCA3 discovery and the development of RNA-seq technologies, the PCa transcriptome has been extensively explored by The Cancer Genome Atlas (TCGA) consortium and others to identify numerous PCa-associated lncRNAs (PCAT family) such as PCAT1, PCAT7 or PCAT114/SChLAP1 (Iyer et al. 2015), (Prensner et al. 2014). However, none of them has been yet introduced into clinical practice because of the variable expression incidence, as for SChLAP1 detected in 25% of PCa cases presenting metastatic traits (Prensner et al. 2013), or low specificity, as PCAT1 or PCAT7, thus infringing their clinical value. Additional efforts are required for more accurate and exhaustive RNA identification, as well as more rigorous validations of clinical potency through independent RNA measurement technologies and clinical cohorts. Regardless a large number of transcriptomic studies and variety of clinical samples analyzed, discovery of RNA-based molecular biomarkers from publicly available RNA-seq datasets is still limited at two levels: (i) most experimental setups are based on poly(A) selected, unstranded cDNA sequencing, and (ii) computational analyses are generally focused on annotated genes and full-length RNA assemblies. This impedes the detection of low and poorly polyadenylated RNAs but also partially degraded RNAs from formalin-fixed paraffin-embedded (FFPE) tissues or other clinical samples (Zhao et al. 2018b), (Zhao et al. 2014). In addition, RNA-seq reads counting is less accurate at 5’ RNA ends or even impossible for co-expressed paired sense/antisense transcripts and for yet unannotated RNAs among noncoding, fusion, repeat-derived transcripts (Audoux et al. 2017), (Davila et al. 2016).

Here, we propose a conceptually novel exploratory framework combining the total stranded RNA-seq of clinical samples and the reference-free DE-kupl algorithm for discovery of novel tumor-specific transcript variations. As a proof-of-concept, we focused on the least explored, noncoding portion of the genome devoid of annotated protein-coding sequences to build an exhaustive catalog of PCa associated subsequences (contigs) embedded into lncRNA genes. The catalog was further refined through minimal filtering to isolate the most potent subset of contigs and validate 21 of them by an alternative NanoString assay in the extended cohort of 144 prostate specimens. From this, a predictive modeling derived a panel of 9 yet unannotated lncRNAs validated for robust expression in an independent TCGA cohort. Importantly, its clinical performance surpassed the PCA3 lncRNA specifically in discrimination of high-risk tumors. The proposed probe-set can be further used for development of a PCa diagnostic test. Moving beyond this point, the proposed computational and experimental platform may serve as a tool for biomarkers discovery for any disease and any clinical task aiming at improved medical care and development of precision medicine approaches.

## RESULTS

### Identification of PCa-specific RNA variants in the Discovery Set by DE-kupl

The biomarker discovery workflow included three major phases: discovery, selection and validation (Fig. 1). First, for discovery, we performed a deep total stranded RNA-seq of ribosomal RNA-depleted RNA samples isolated from prostate tissues after radical prostatectomy (*Discovery Set*, PAIR cohort, Supplemental Table S1). This *Discovery set* was processed by DE-kupl to identify tumor-specific transcripts. DE-kupl directly queries FASTQ files for subsequences (k-mers) with differential counts/expression (DE) between two conditions (Fig. 2A) (Audoux et al. 2017). Overlapping k-mers are then assembled into contigs and, in a final step, mapped to the human genome for annotation. In the aim to focus exclusively on novel, yet unannotated RNA elements, k-mers exactly matching GENCODE annotated transcripts were masked. We eventually retained contigs longer than 200 nucleotides and showing adjusted p-values below 0.01 to capture the most significant expression changes linked either to new transcriptional or processing events within known or putative lncRNA loci.

**Figure 1.**
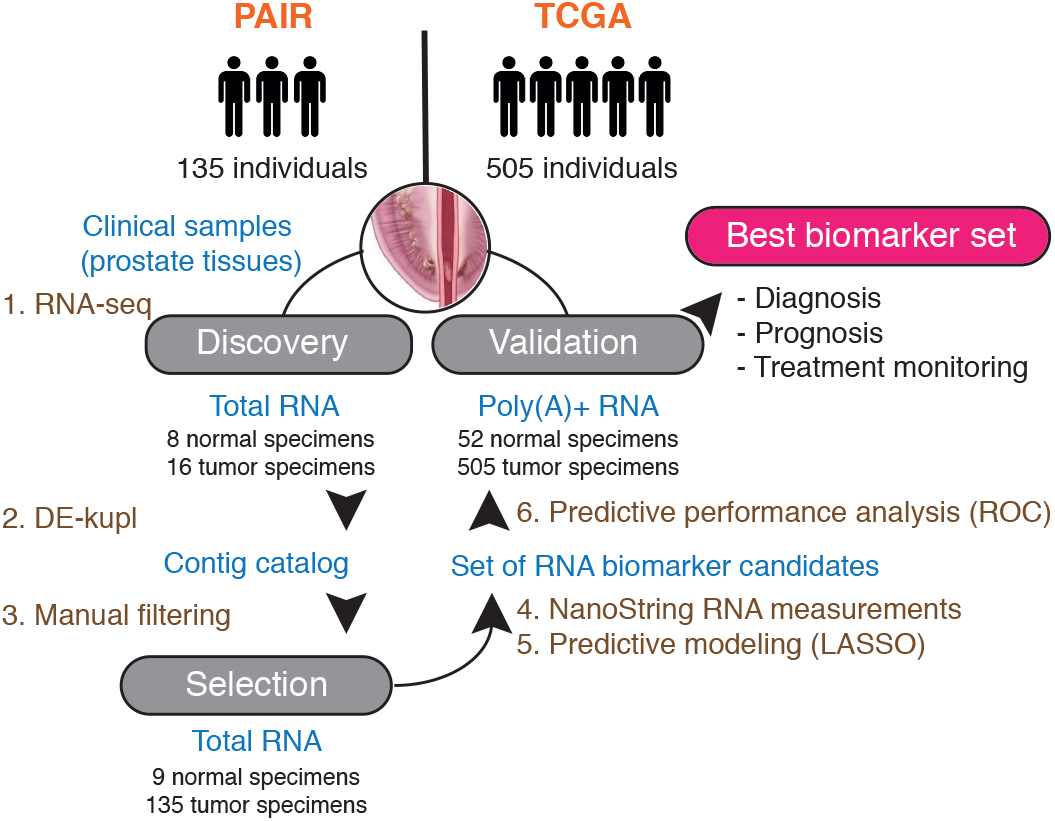
Experimental and computational workflow for discovery and validation of RNA-based clinical biomarkers. Raw total stranded RNA-seq data of a small clinical cohort is processed by DE-kupl to allow comparison of 8 normal against 16 tumor specimens (in this case formaldehyde fixed paraffin embedded tissues from radical prostatectomy) and cataloguing of all differentially expressed RNA variations (contigs). The whole set is filtered according to desired criteria and the top ranked contigs are selected for an independent experimental validation by NanoString in the extended clinical cohort. Finally, predictive modeling infers the best panel of candidate RNAs for validation of its clinical potency in an independent cohort (in this case TCGA).

**Figure 2.**
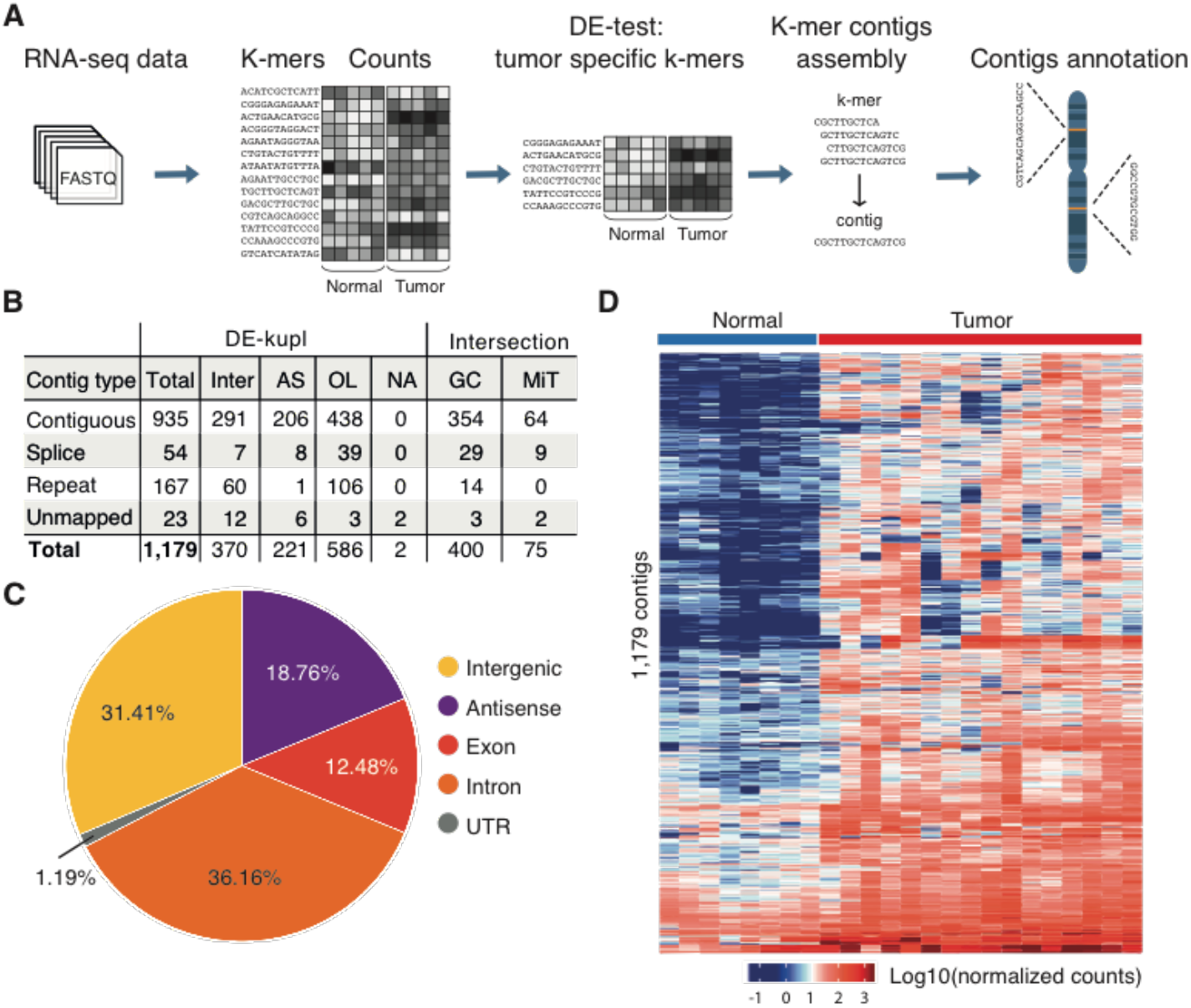
K-mer decomposition protocol for detection of differentially expressed RNA variants in PCa. (**A)** DE-kupl workflow with principle steps of contigs counting, DE-test and filtering, assembly and annotation. **(B)** Catalog of DE-kupl contigs of different subgroups: contiguous - contigs mapped as unique fragments; spliced - contigs mapped as spliced fragments; repeat - multiply mapped contigs; Inter - contigs mapping into intergenic regions, OL - overlapping GENCODE lncRNA annotations, AS - antisense to a protein-coding or a noncoding gene. Contigs of each subgroup showing 50% sequence overlap with GENCODE (GC) and MiTranscriptome (MiT) annotated genes are counted. **(C)** Pie chart of contigs distribution across GENCODE annotated features. **(D)** Unsupervised hierarchical cluster heatmap of Log10(normalized counts) of 1,179 contigs assessed in 8 normal and 16 tumor specimens by total stranded RNA-seq of the *Discovery Set*.

With these criteria, we identified 1,179 tumor up-regulated contigs assigned to four main categories according to their mapping features: contiguous (uniquely mapped) contigs (N=935), splice variants (N=54), repeats (N=167) and unmapped contigs (N=23) (Fig. 2B, Fig. S1). Among them, 33.93% and 6.36% were embedded into already referenced GENCODE or MiTranscriptome lncRNA genes, respectively, but represented new sequence variations or RNA processing events, as PCAT7 (ctg_111348, P16) or CTBP1-AS (ctg_25348, P10). The rest mapped to intergenic noncoding locations or antisense to referenced protein-coding or noncoding genes (Fig. 2C). An unsupervised clustering of prostate specimens based on contigs expression counts allowed proper discrimination of tumor from normal tissues of the *Discovery Set* (Fig. 2D).

In conclusion, DE-kupl identified thousands of PCa-associated RNA variants for the majority embedded into yet unreferenced transcripts which may represent putative novel lncRNAs. This depository was further explored for clinical relevance.

### Naïve assembly of Transcription Units identifies novel prostate cancer associated lncRNAs

To complement the reference-free protocol, we applied a reference-based protocol to build a catalog of lncRNAs from the same *Discovery Set*. Total RNA-seq produces much more intronic and exon-exon junction reads than poly(A)-selected RNA-seq, which is deleterious for splice graph-based assemblers such as Cufflinks (Kukurba and Montgomery 2015), (Hayer et al. 2015a). To bypass this difficulty, we developed a more straightforward lncRNA annotation pipeline, HoLdUp, which identifies transcription units (TU) based on coverage analysis (Fig. 3A). In this workflow, uniquely mapped reads were assembled into TUs and mapped to the GENCODE annotation to extract intergenic and antisense lncRNAs (see Methods for details). They were further ranked according to their expression level, presence of splice junctions and existence of matched expressed sequence tags (EST). In total, we retained 168,163 TUs with above-threshold expression of 0.2 quartile of mRNA expression (Class 2) and, within this group, the most robust 2,972 TUs with at least one splice junction and one EST (Class 1) (Fig. 3B). Globally, newly detected transcripts were as much expressed as GENCODE annotated lncRNAs but lower than mRNAs (Fig. S2A). Only 0.33% of Class 1 lncRNAs were present with at least 50% nucleotide sequence overlap in the recent GENCODE v26 catalog and 43.37% of TUs in the MiTranscriptome lncRNA repertoire; the rest represented putative novel lncRNA genes (Fig. 3B, Fig. S2B). Of 2,972 TUs, DE analysis retrieved 127 of Class 1 TUs significantly up-regulated in tumor specimens (adjusted p-value below 0.01, DESeq), including multiple intergenic transcripts and transcripts antisense to protein-coding genes, such as *HDAC9, TPO, FBXL7*.

**Figure 3.**
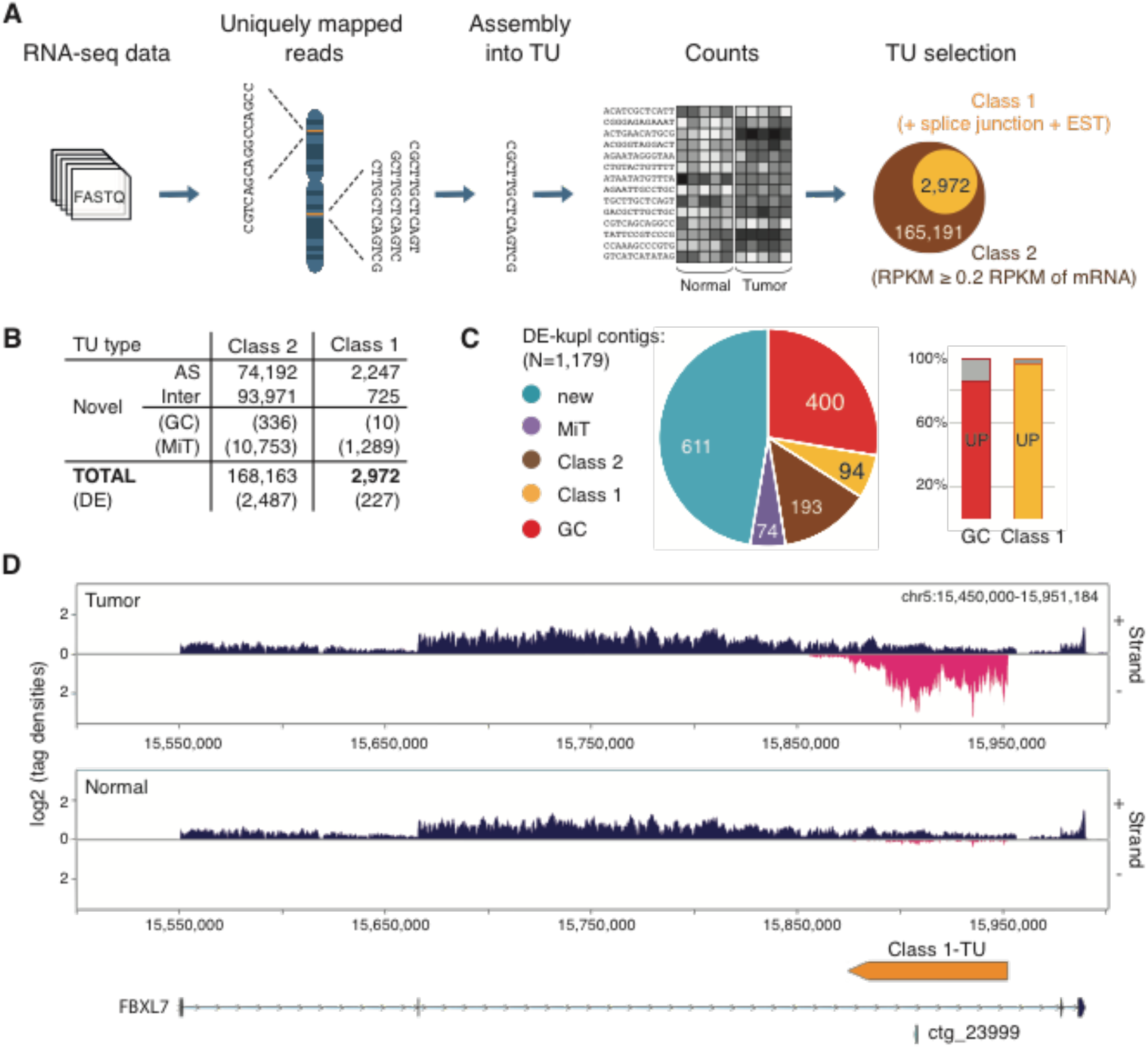
Reference-based lncRNA discovery from total stranded RNA-seq. **(A)** The HoLdUp protocol for the *ab initio* assembly of TUs constituting putative lncRNA genes and their classification into Class 2 and Class 1 TUs according to robustness of detection. **(B)** HoLdUp catalog and TUs overlap with GENCODE v26 (GC) and MiTranscriptome (MiT) annotated lncRNAs. DE stands for differentially expressed transcripts (DESeq adj. p-value < 0.01). **(C)** Pie chart representation of non-exclusive distribution of DE-kupl contigs across different lncRNA annotations: MiTranscriptome (violet), Class 1 (yellow), Class 2 (brown), GENCODE (red) and novel (blue), number of contigs is marked in each section. Proportion of DE-kupl contigs embedded into up-regulated (UP) GENCODE (red bar) and Class 1 (yellow bar) lncRNAs is expressed as a histogram. **(D)** Ving-generated RNA-seq profiling along plus (+) and minus (-) strands of chr5:15,500,295-15,939,910 in tumor and normal prostate specimens: the GENCODE annotated protein-coding gene *FBXL7* (blue), antisense DE-kupl contig ctg_23999 (P22) and antisense HoLdUp Class 1-TU (orange). Arrow-lines and rectangles represent introns and exons, respectively. TU = transcription unit; DE = differentially expressed; RPKM = Reads per kilo base per million mapped reads.

Intersection of DE-kupl contigs with HoLdUp TUs and the recent GENCODE lncRNA annotation showed that 687 DE-kupl contigs out of 1,179 make part of the stand-alone transcripts. Moreover, up to 85.5% and 96.8% DE-kupl contigs embedded into GENCODE and HoLdUp Class 1 lncRNA genes, respectively, were also detected by DESeq as significantly up-regulated transcripts in the same dataset, when the RNA-seq reads were counted within the entire TU (Fig. 3C; Fig. S2C). One such example is the contig ctg_23999 (P22) embedded into a novel HoLdUp assembled Class 1 TU antisense to the protein-coding *FBXL7* gene (Fig. 3D).

In conclusion, the reference-based assembly protocol HoLdUp is complementary to DE-kupl and allows attributing short RNA subsequences to whole transcription units. Nevertheless, DE-kupl was more powerful illuminating much more transcriptomic variations not only within the annotated loci but also within putative new noncoding regions in highly complex and heterogeneous total RNA-seq datasets of clinical origin.

### Selection of a restricted set of 23 PCa RNA contigs showing the highest differential expression

We further leveraged the DE-kupl contig catalog to define a robust PCa signature among putative new lncRNAs using several filters (Fig. S3A). First, contigs were sorted according to their adjusted p-value and, second, were visually selected using the Integrative Genomic Viewer (IGV) applying the following criteria: (i) when several contigs were present within the same genomic region (5 kilobase window) the contig with the lowest adjusted p-value was retained, (ii) contigs antisense to expressed exons, bidirectional or positioned in close vicinity to other transcribed protein-coding genes were filtered out. We also retained contigs assigned to already known PCa associated lncRNAs, such as CTBP1-AS (ctg_25348, P10), PCAT7 (ctg_111158, P6) and PCAT1 (ctg_105149, P18), or lncRNAs referenced elsewhere as ctg_104447 (P11) mapped into LOC283177, ctg_123090 (P5) into AC004066.3, and ctg_73782 (P8) into LINC01006; all of which passed the aforementioned selection criteria. Notably, the RNA-seq visualization of a new contig antisense to the protein-coding *FBP2* gene (ctg_28650, P2) revealed that it most likely makes part of the PCAT7 lncRNA as an extension of its last exon (Fig. S3B). The contig ctg_28650 (P2) was retained in the restricted list as the strongest candidate antisense to *FBP2*, overcoming ctg_111158 (P6) assigned to the *PCAT7* gene. In total, 23 candidates belonging to contiguous (N=21), spliced (N=1) or repeat (N=1) subgroups of contigs were selected for further validation, all being expressed at least 6 times more in tumor tissues comparing to normal prostate (Fig. S3C, Supplemental Table S2). Among them, 12 candidates mapped antisense to annotated protein-coding or lncRNA genes and 11 located to intergenic regions. To facilitate further reading, contigs’ identity (ID) are replaced by probes’ ID from P1 to P23 according to increasing p-values of DE of the *Discovery set* (Supplemental Table S2).

Following the manual filtering we aimed to validate the expression of selected 23 contigs in the extended PAIR cohort of 9 normal and 135 tumor specimens (*Selection Set*) (Supplemental Table S3). For this purpose, an alternative RNA quantification procedure based on the NanoString nCounter^TM^ platform for direct enzyme-free multiplex digital RNA measurements was carried out (Fig. 4A). In addition to DE-kupl contigs, a probe for PCA3 was used as a benchmark lncRNA. We also measured the expression of six housekeeping genes and selected three lowly expressed mRNAs (GPATCH3, ZNF2, ZNF346) as custom internal controls for relative quantifications (Supplemental Table S4, Fig. S4).

**Figure 4.**
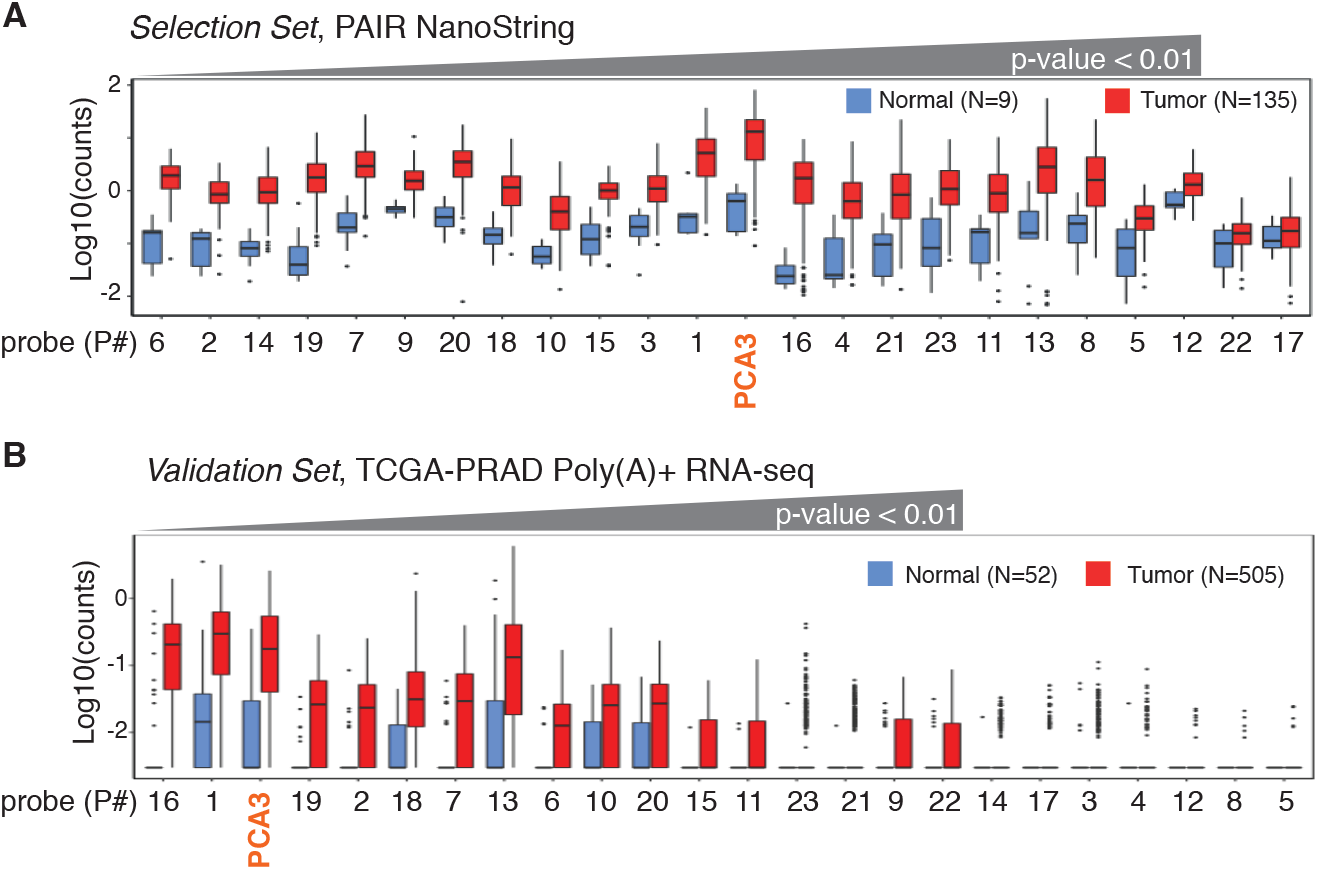
Expression of DE-kupl contigs in PAIR and TCGA-PRAD cohorts. **(A)** Box-plot of Log10(counts) of PCA3 and 23 DE-kupl contigs in 144 PAIR specimens of the *Selection set* by NanoString. **(B)** Box-plot of Log10(counts) of PCA3 and 23 DE-kupl contigs in 557 TCGA-PRAD specimens of the *Validation Set* by poly(A)+ unstranded RNA-seq. Normal tissues - in blue, tumor tissues - in red.

The NanoString assay revealed that all DE-kupl contigs were expressed at a lower level than PCA3, but still 21 out of 23 contigs were significantly overexpressed (Wilcoxon p-value < 0.01) in tumor specimens (Fig. 4A, Supplemental Table S5). Two contigs, intergenic P22 (ctg_119680) and repeat P17 (ctg_36195) did not show significant difference in expression between normal and tumor specimens. Ranking according to p-values revealed 12 contigs better than PCA3. Among the top DE contigs were those embedded into *PCAT1* (ctg_105149, P18), *CTBP1-AS* (ctg_25348, P10) and *PCAT7* (ctg_111158, P6) genes, and the rest were assigned to novel lncRNAs. Notably, apart from P17 (ctg_36195) and P22 (ctg_119680), expression measurements were consistent between the two technologies, total stranded RNA-seq and NanoString, though the p-values ordering was different (Fig. S5, Supplemental Table S6).

Thus, 21 out of 23 contigs were validated in the extended set of RNA specimens using the independent single-molecule measurement technology.

### Validation of contig-based RNA candidates in an independent clinical cohort

Independent validation of DE-kupl contigs was done using the biggest PCa clinical resource of 557 poly(A)+ RNA-seq datasets, including 52 normal and 505 tumor tissues from radical prostatectomy (TCGA-PRAD cohort, *Validation Set*) (Fig. 1, Supplemental Table S7).

The occurrence of sequences representing 23 DE-kupl contigs was measured and compared to PCA3. In total, 16 out of 23 DE-kupl contigs had significant support for overexpression in tumor specimens in the TCGA-PRAD cohort (Wilcoxon p-value < 0.01, FC > 2) (Fig. 4B, Supplemental Table S8). Among the best scored candidates, the two novel DE-kupl contigs, P16 (ctg_111348) antisense to *DLX1* and intergenic P1 (ctg_17297), surpassed PCA3 ranked third. However, important discrepancies were observed between expression counts in poly(A)+ RNA-seq TCGA datasets and NanoString or total RNA-seq PAIR datasets. First, P22 (ctg_119680) was detected as DE in TCGA-PRAD, but failed the DE test when measured by NanoString (Fig. 4, Fig. S5). Second, the expression of nine DE-kupl contigs were near the base line in the TCGA dataset, including those showing relatively high expression and low p-values in the PAIR cohort, such as P14 (ctg_61528) antisense to *TPO* or the intergenic P9 (ctg_9446). Detection of these contigs in TCGA-PRAD was compromised independently of their genomic location (intergenic or antisense) or of the expression level of a sense-paired gene. We hypothesized that it is most likely due to a relatively low RNA-seq coverage and/or to a loss of poorly or non-polyadenylated transcripts during cDNA library preparation in the TCGA experimental setup. Finally, ranking of contigs according to increasing p-values was very different between the *Validation*, *Discovery* and *Selection* Sets highlighting remarkable discrepancies either between technologies or clinical origins.

Regardless all experimental biases, 16 out of 23 DE-kupl contigs were validated in the independent clinical cohort as significantly overexpressed in tumors. This cohort was further used for validation of clinical potency of contigs.

### Expression of DE-kupl contigs is independent on tumor risk and recurrence metrics

Several clinical studies have revealed high heterogeneity of expression and low efficiency of the PCA3 biomarker in detection of high-risk tumors, questioning its robustness and reliability in PCa diagnostics (Alshalalfa et al. 2017), (Fenstermaker et al. 2017). We assessed contig expression in tumors of different clinical metrics. For risk prognosis, the most common metric is a three-group risk stratification system established by D’Amico in 1998 (D’Amico et al. 1998), which takes into account preoperative PSA level, biopsy Gleason score and clinical TNM stage. As mentioned above, this scheme is highly debated due to disagreements over the PSA score in relation to PCa over-diagnosis (Loeb et al. 2014), (Carlsson et al. 2012). To define a molecular signature independent of PSA, we excluded this criterion and categorized tumor specimens into low-, intermediate- and high-risk groups uniquely on the basis of Gleason and TNM features, below referred to as naïve indexing (Fig. S6). In addition to risk assessment, we also separated specimens in two subgroups depending on the tumor recurrence status (Fig. S6B). Then, expression of PCA3 and the 23 DE-kupl contigs were compared for each subgroup of the *Selection Set*.

To evaluate the robustness of contig expression, we ranked probes by decreasing FC for high-risk (HR) against low-risk (LR) tumors and positive versus negative recurrence status (Fig. 5).

**Figure 5.**
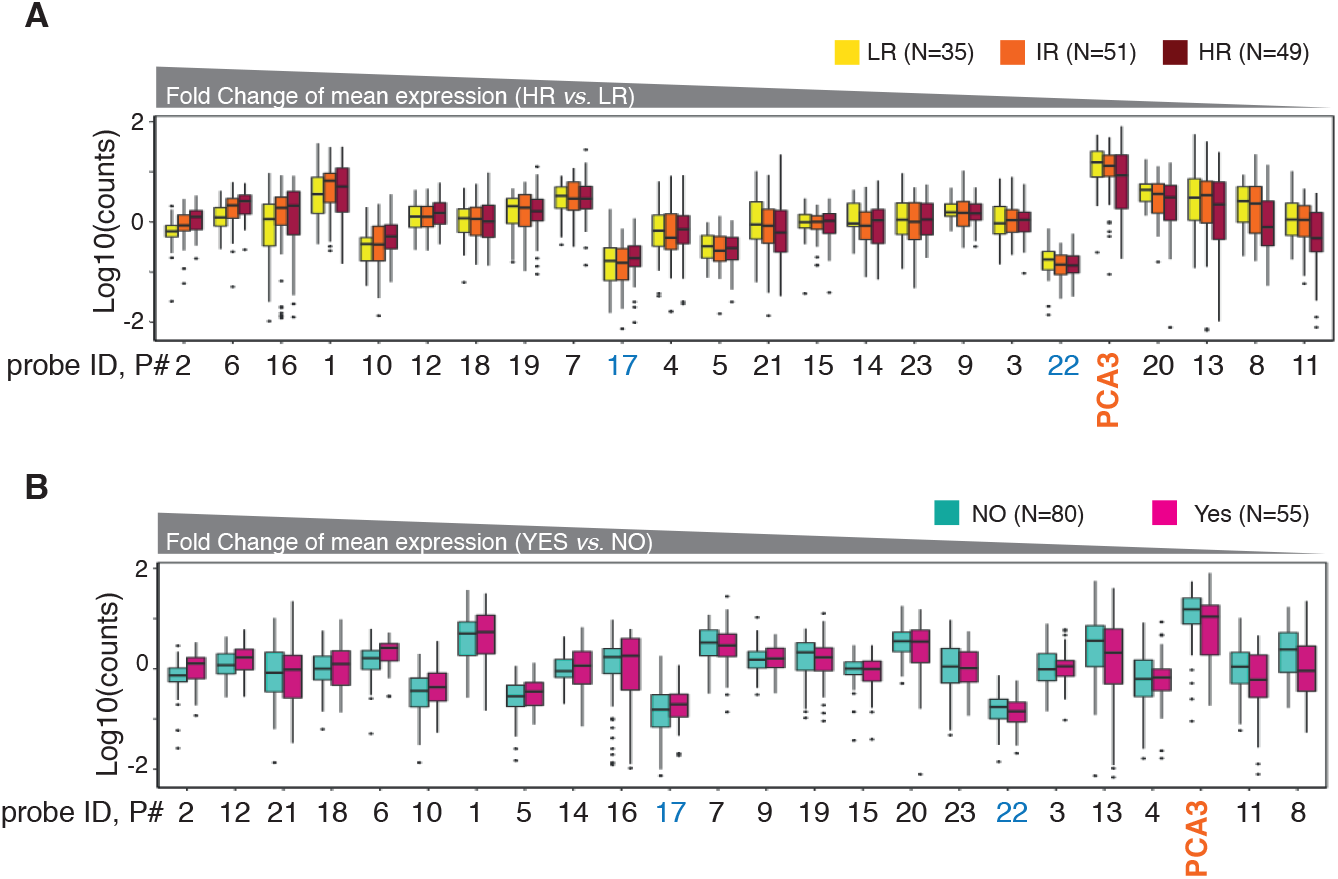
Box-plot of Log10(counts) of PCA3 and DE-kupl contigs in prostate specimens of the PAIR cohort (*Selection Set*) depending on tumor risk. **(A)** and recurrence status **(B)** assessed by NanoString. PCA3 is marked in orange, and the contigs showing insignificant expression change between normal and tumor specimens are in blue. Contigs are ordered by the decreasing FC of mean expression in HR *vs*. LR specimens in the A panel and in Yes *vs.* NO recurrence specimens in the B panel.

The majority of contigs showed robust expression independently of the tumor classification. In contrast, the PCA3 level was more disperse with the lower median and mean expression and higher p-values in high-risk and recurrence positive specimens (Supplemental Table S9). While considering only 21 significantly overexpressed contigs, 17 of them outperformed PCA3 in both contrasts (Supplemental Table S9). Notably, contigs P6 (ctg_111158) and P2 (ctg_28650) both antisense to *FBP2*, P10 (ctg_25348) embedded into CTBP1-AS, but also the novel P16 contig (ctg_111348) antisense to *DLX1* and the intergenic P1 (ctg_17297) performed best.

In conclusion, the majority of DE-kupl contigs showed robust expression independent of tumor metrics. Hence, even if used alone, they may offer a better clinical potency for PCa diagnosis than PCA3.

### Inferring a multiplex RNA-probe panel and evaluation of its performance in PCa diagnosis

To extract parsimonious probe subsets predicting the tumor status, we applied LASSO (Least Absolute Shrinkage and Selection Operator) logistic regression on the *Selection Set* of 144 PAIR specimens (Ghosh and Chinnaiyan 2005). First, the initial 21 DE-kupl contigs and PCA3 validated for expression by NanoString were submitted to LASSO to define the best mixed signature comprised of already known and yet unannotated lncRNA probes for discrimination of tumor from normal tissues (Fig. S7A). Then, LASSO was performed with the probe subset composed uniquely of contigs assigned to putative novel lncRNAs (N=15) to infer the best new-lnc RNA signature. It resulted in two panels of 9 mixed and 9 new-lnc RNA candidates (Fig. 6A, Fig. S7B). Retrieved signatures were then used to predict a tumor status in the *Validation Set* of the TCGA-PRAD cohort using a leave-one-out cross-validated boosted logistic regression. To assess the sensitivity of DE-kupl contigs in PCa diagnosis, a predictive accuracy index, Area Under Curve (AUC) of the receiver-operating characteristic (ROC), was calculated for each signature and PCA3 alone in the PAIR (*Selection Set*) and TCGA-PRAD (*Validation Set*) datasets (Fig. 6B; Fig. S7B). Remarkably, all signatures still hold their predictive capacity in the independent TCGA-PRAD cohort in spite of the important differences in experimental setups between the two studies. Both markedly outperformed PCA3 for tumor detection with AUC of 0.92 for mixed and of 0.91 for new-lnc signatures against AUC of 0.73 for PCA3 (Fig. 6B and 6C). In addition, these signatures were much better in predicting high-risk tumors where PCA3 is particularly inaccurate (Fig. 6C). Remarkably, the new-lnc RNA signature of 9 contigs composed uniquely of yet unannotated lncRNAs predicted the tumor status with the same performance as the mixed signature. Logistic regression did not retain PCA3 within the mixed signature set, instead contigs embedded into the well characterized PCAT1 lncRNA and into two already annotated but yet functionally uncharacterized lncRNAs LOC283177 and LINC01006 were present.

**Figure 6.**
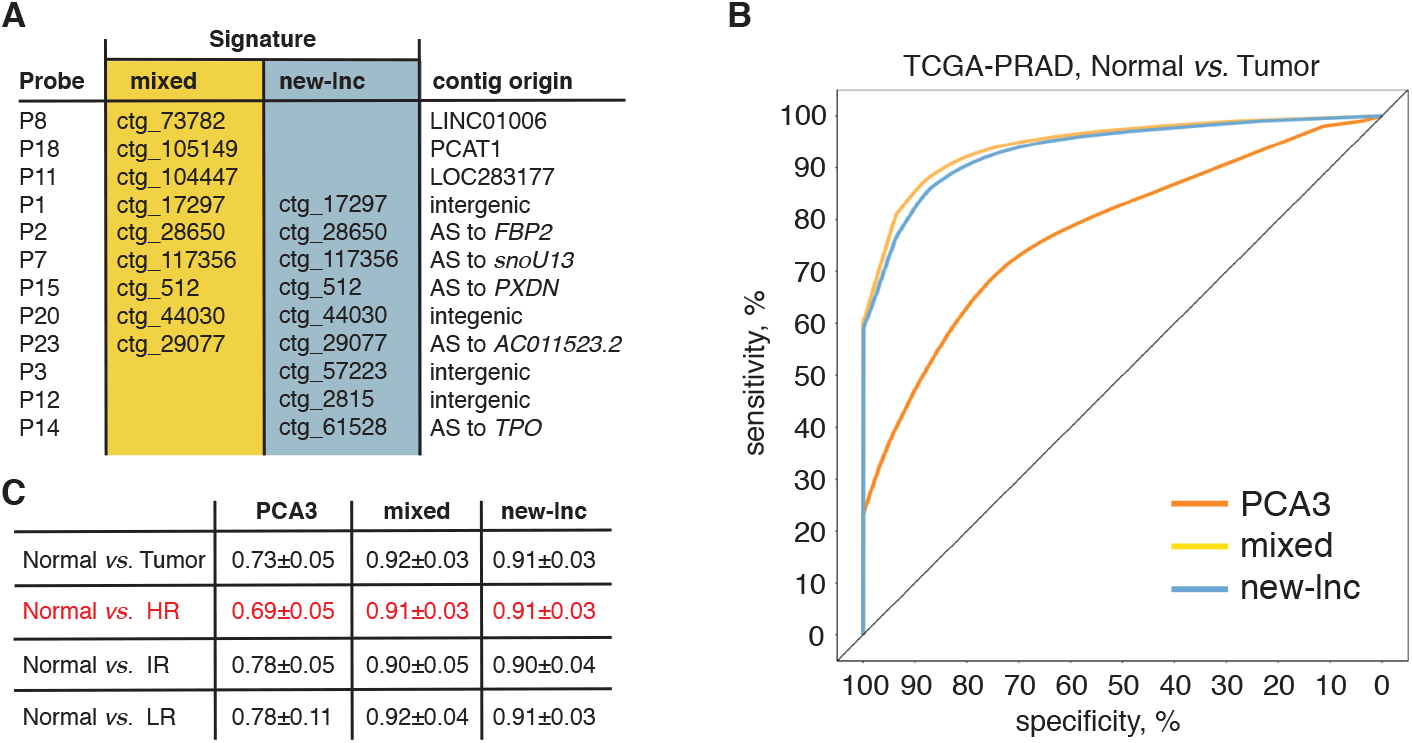
Predictive performance of PCA3 and multiplex mixed and new-lnc RNA signatures inferred from the LASSO penalized logistic regression. **(A)** Multiplex biomarker signatures composed of either known and unannotated RNAs (mixed) or of only unannotated RNAs (new-lnc). **(B)** ROC for the PCa prediction in the TCGA dataset (*Validation Set*) using two signatures and PCA3 alone. **(C)** Mean and standard deviation of AUC computed over 100 samplings of the *Validation Set* for PCA3 and two signatures to classify tumors according to their risk status. AS = antisense; AUC = area under the curve; HR = high-risk, IR = intermediate-risk, LR = low-risk tumors.

This result highlights both the incompleteness of current cancer transcriptome datasets and the biological value of transcript information that can be extracted through adequate experimental (total stranded RNA-seq and NanoString quantification) and computational (DE-kupl) tools. The resulting signature demonstrated a sensitivity and robustness towards tumor risk surpassing the state of the art for discrimination of prostate cancer. Furthermore, the nine-probe RNA signature performed independently of tumor origin and clinico-pathological characteristics, but also independently of the technology used for RNA measurements.

## DISCUSSION

Molecular biomarker assays are invaluable tools in cancer diagnosis, prognosis and treatment follow-up. Within this scope, sequencing technologies unveiled the pervasiveness and diversity of the human transcriptome, promoting lncRNAs as important cancer signatures (Schmitt and Chang 2016). These molecules are highly dynamic and reflect cellular states in a sensitive and specific way due to their involvement in genetic and regulatory flows of information. However, the variety of RNA forms and high heterogeneity of expression present a challenge for their detection and proper quantification in clinical samples. Predominant microarray and unstranded poly(A)+ RNA-seq based approaches allowed identification of numerous lncRNAs with tumorigenic function. However, their clinical performance as biomarkers stays rather poor due to the aforementioned RNA features hindering RNA detection, quantification and clinical validation under conventional experimental setups. Here, we presented an innovative experimental and computational platform that permits discovery of RNA biomarkers of high clinical potency from total stranded RNA-seq datasets of clinical origin.

As a proof-of-concept, we focused on PCa as the only type of cancer using, so far, an RNA-based diagnostic test (Progensa™). The *Discovery Set* based on comparison of 8 normal with 16 tumor specimens from total RNA-seq datasets was processed by DE-kupl to extract the most significant differentially expressed subsequences in the form of k-mer contigs. Further filtering based on contig length, genomic position and expression levels powered the pipeline towards the discovery of putative lncRNAs, for the majority, yet unreferenced in the human transcriptome. Then, the catalog of contigs was manually refined and tested for expression using the NanoString single-molecule RNA counting technology in the extended cohort of 144 specimens. Contig expression was next assessed in the independent, publicly available TCGA-PRAD dataset generated by the poly(A)+ unstranded RNA-seq technology. The expression of contigs was systematically compared to that of the benchmark biomarker lncRNA, PCA3. In total, 16 out of 23 contigs were validated in both setups but with important differences. Primarily, RNA measurements were consistent between two different technologies: NanoString and total stranded RNA-seq. In contrast, the TCGA poly(A)+ unstranded datasets revealed weakness and high heterogeneity of contig counts over the selected regions, resulting in unexpectedly low signals even for PCA3, considered as a highly expressed lncRNA. Our results promote the total stranded RNA-seq as a first-choice strategy for discovery of RNA biomarkers from clinical samples and when searching for transcripts others than highly abundant mRNAs. It reflects far more precisely the transcriptomic landscape of clinical samples and, hence, is more advantageous as a *Discovery Set* for development of clinical tests. At the same time, full-length transcript assembly from short-read sequencing is inaccurate, time and computer memory consuming, and this is aggravated by the added complexity of total (ribo-depleted) RNA-seq libraries (Hayer et al. 2015b). DE-kupl bypasses this issue by directly extracting from raw data RNA subsequences significantly overexpressed in a defined condition. In PCa tissues, this allowed identification of 1,179 lncRNA-hosted candidates. Further analysis isolated a restrained set of 9 contigs either within putative new lncRNAs or mixed annotated and novel lncRNAs allowing PCa diagnosis independently of tumor risk classifications with higher than the actual PCA3. Remarkably, the best performing mixed signature did not include PCA3, consistent with the low potency of this biomarker in detection of aggressive tumors. Instead, both mixed and new-lnc RNA signatures contained contigs embedded into putative novel lncRNA genes whose function in PCa progression will be important to explore. Among them, the P23 contig (ctg_29077) is antisense to AC011523.2, an intergenic lncRNA, co-transcribed with P23 in PCa specimens. This region is part of a super-enhancer, annotated in several PCa cell lines, located between *KLK15* and the PSA encoding *KLK3* genes (Jiang et al. 2019). Moreover, it has also been described as an enhancer bi-directionally transcribed into enhancer (e)RNAs and regulating expression of the neighboring *KLK3* and *KLK2* genes through eRNA and Med1-dependent chromatin looping in androgen-dependent LNCaP and VCaP, and androgen-independent LNCaP-abl cells (Hsieh et al. 2014). Presence of the P23 contig within the mixed and new-lnc RNA signatures supports, in addition to clinical potency, a possible regulatory function of the k-mer containing RNA contigs inferred by DE-kupl. More globally, the majority of DE-kupl contigs within co-transcribed sense-antisense pairs were annotated as super-enhancers in prostate tissues and cell lines or other biosamples, *e.g.* P15 (ctg_512), P7 (ctg_117356), and P4 (ctg_63866) (Jiang et al. 2019). In most cases, their function in gene expression regulation and chromatin configuration has not yet been investigated and experimentally validated, but it is tempting to speculate that defined sense-antisense transcripts may influence a super-enhancer activity and, consequently, may fine-tune the expression of neighboring genes.

In this work, we propose DE-kupl as a tool for discovery of novel disease-associated transcriptomic variations, which can be further explored for biological and clinical relevance. As a pilot project, we oriented the pipeline towards the discovery of novel lncRNAs, but using proper masking and filtering criteria defined by the investigator, other variant transcripts including single nucleotide variations (SNV), novel splice events, gene fusion, circular RNAs or exogenous viral RNAs could be probed. The workflow can be applied to any RNA-seq datasets of any clinical origin to generate a probe panel that may be implemented as a multiplex platform for simultaneous detection of RNAs in clinical samples. Moreover, different experimental contrasts (normal *vs.* pathology, low-*vs.* high-risk grade, treatment resistant *vs.* sensitive, *etc.*) will define the biomarker usage in diagnosis, prognosis or other clinical applications, hence, providing clinicians and researchers with a simple and highly sensitive tool for genomic and personalized medicine.

## METHODS

### Tissue samples

Tumor and normal biopsy specimens were retrospectively collected from prostate cancer patients who provided informed consent and were approved for distribution by the H. Mondor institutional board (PAIR cohort). Tumors classification in low-, intermediate- and high-risk prognosis was performed according to Gleason and TNM scores and regardless PSA values (Supplemental Table S1, S3).

### RNA extraction, quantification and cDNA library production

Total RNA was extracted using the TRizol reagent (ThermoFisher), according to manufacturer’s procedure, quantified and quality controlled using a 2100 Bioanalyzer (Agilent). RNA samples with RNA Integrity Number (RIN) above 6 were depleted for ribosomal RNA and converted into cDNA library using a TruSeq Stranded Total Library Preparation kit (Illumina). cDNA libraries were normalized using an Illumina Duplex-specific Nuclease (DSN) protocol prior to a paired-end sequencing on HiSeq™ 2500 (Illumina). At least 20x coverage per sample was considered as minimum of unique sequences for further data analysis.

### RNA-sequencing data

Raw paired-end strand-specific RNA-seq data was generated by our laboratory from ribo-depleted total RNA samples of prostate tissues (8 normal and 16 tumor specimens; Supplemental Table S1) and can be retrieved from the gene omnibus portal (GEO), accession number <u>GSE115414</u>.

TCGA prostate cancer poly(A)-selected RNA-seq and corresponding clinical data were obtained from publicly available TCGA dataset (http://cancergenome.nih.gov), 557 inputs in total (52 normal and 505 tumors of high-(N=240), intermediate-(N=128) and low-risk (N=132) groups. Among them, 369 patients showed no tumor recurrence, 108 presented a new tumor event (Supplemental Table S7).

### Computational workflow for k-mer contigs discovery from total stranded RNA-seq dataset

DE-kupl run was performed from (June 2017) with parameters ctg_length 31, min_recurrence 6, min_recurrence_abundance 5, pvalue_threshold 0.05, lib_type stranded, diff_method DESeq2. K-mer masking was performed against the GENCODE v24 annotation. DE-kupl analysis of the 8 against 16 PAIR RNA-seq prostate libraries yielded 124,809 DE contigs, in total. Contigs were annotated by alignment on the hg19 human genome assembly using the DE-kupl *annotate* procedure. We further selected contigs of size above 200 nucleotides and classified them into four categories (contiguous, repeat, spliced, unmapped) based on their location and mapping features.

### Computational workflow for reference-based *ab initio* transcripts assembly from total stranded RNA-seq dataset (HoLdUP)

The human genome version hg19 and the GENCODE v14 annotation were used in this study. First, we performed a quality control of all sequencing data by FastQC Babraham Bioinformatics software. Reads were mapped using TopHat 2.0.4, allowing 3 mismatches and requesting uniquely mapped reads which were further assembled using the BedTools suite. Overlapping contigs from all libraries were merged and only contigs supported by at least 10 reads in either library were further assembled in segments if mapped in the same strand and separated by less than 100 nucleotides. We compared segments to the GENCODE v14 annotation to extract antisense and intergenic TU longer than 200 nucleotides. To classify lncRNAs, we applied the following criteria: (i) an expression level above 0.2 quartile of mRNA expression in at least one condition per tissue (Class 2); (ii) within this class, all TUs containing at least one TopHat-identified exon-exon junction and at least one spliced EST from UCSC mapped contigs were assigned to Class 1. The whole catalog, the R code and Data Tables can be provided upon request.

### Overlap between GENCODE, MiTranscriptome, DE-kupl and HoLdUp catalogues

Intersection between transcripts was counted only in case of 50% overlap of nucleotide sequence between genomic coordinates of each fragment.

### Differential expression analysis

Read counting was performed on the compiled annotation (GENCODE v26, HoLdUp Class 1 and Class 2) for each sample, using *featureCounts* 1.6.0 with the following parameters: -F "SAF" -p -s 2 -O and the DESeq R package (Love et al. 2014). Only RNAs with adjusted p-value below 0.01 were retained as differentially expressed to constitute the prostate tumor signature.

### NanoString nCounter Expression Assay

100 ng of total RNA was used for direct digital detection of 29 target transcripts: 6 housekeeping genes (*RPL11, GAPDH, NOL7, GPATCH3, ZNF2* and *ZNF346*), 23 contigs and the one known PCa-associated lncRNA, PCA3. Each target gene of interest was detected in RNA samples of 144 specimens (9 normal and 135 tumor) of the PAIR cohort (Supplemental Table S3) on NanoString nCounter V2 using reporter and capture probes of 35- to 50-nucleotide targeting sequences. Data was normalized through the use of NanoString’s intrinsic positive controls and then contig expression was calculated relative to the average signal of three housekeeping genes (*GPATCH3, ZNF2* and *ZNF346*). Raw and normalized data for each specimen, mean and fold change expression in normal against tumor samples are presented in Supplemental Table S4 and S5.

### Contig expression measurements in TCGA-PRAD datasets

DE-kupl provides representative k-mers for each differentially expressed contig. We converted the TCGA-PRAD FASTQ files to k-mer counts using *Jellyfish count* and counted representative k-mers in each Jellyfish count file using the *Jellyfish query* command. Counts were normalized by total number of reads in corresponding libraries. To determine whether counts of DE-kupl derived representative k-mer were a reliable proxy for evaluating contig expression, we compared representative k-mer counts to average counts from k-mers sampled along each contig. All individual counts were obtained using *Jellyfish Dump* files produced for each TCGA-PRAD library. Sampling was performed as follows: (i) we extracted all k-mers from the contig that were unique in the Ensembl human v91 transcript reference, and (ii) from this list we sampled 10 regularly spaced k-mers, starting from the first 10% and ending in the last 10% of the list. This sampling procedure was repeated four times for each contig. For the whole TCGA library and each contig, the 10 k-mer counts obtained by Jellyfish were averaged, yielding one average count per sample per library. Correlations between sample counts and representative k-mer counts are shown in Fig. S8 for two DE-kupl contigs.

### RNA-sequencing data visualization

RNA-seq reads profiling along a locus of interest was performed using our in-house R script VING (Descrimes et al. 2015). The normal samples were assigned to the group “controls” and the tumor specimens – to the group “cases”, with the assumption that the “cases” should have higher values than “controls”.

### Unsupervised clustering of prostate specimens

Specimens were ranked based on the Log10(expression counts) levels of contigs assessed by the NanoString nCounter assay using a ComplexHeatmap R-package (Gu et al. 2016).

### Variable selection using the LASSO penalized logistic regression and external validation of signatures

Signature inference was performed in R using the normalized *Selection Se*t (23 probes in 144 observations) as a variable selection dataset and contigs counts table of the *Validation Set* (23 probes in 557 observations) as an external validation dataset (R Core Team). First, we performed penalized logistic regression using the *glmnet* R package to select probes predicting the tumor status on the *Selection Set* upsampled to correct the imbalance class distribution (9 normal versus 135 tumor specimens) (Friedman et al. 2010). Selection was performed using all probes (signature_mixed including PCA3) or using only new-lnc RNA contigs only (signature_new-lnc) (Fig. S7). Second, we built predictors using the boosted logistic regression from the *caTools* and *caret* packages (Tuszynski 2008), (Kuhn 2008). AUCs were computed using the *precrec* package on 100 training and testing datasets (Saito and Rehmsmeier 2017), sub-sampled from the initial dataset (Normal *vs.* Tumor, Normal *vs.* HR, Normal *vs.* IR and Normal *vs.* LR) using the *sample.split* function from the *caTools* package.

## DATA ACCESS

TCGA prostate cancer poly(A)-selected RNA-seq and corresponding clinical data can be obtained from TCGA portal (https://www.cancer.gov/tcga).

## ACKNOWLEDGEMENTS

We deeply thank Dominika Foretek and Maxime Wery for editorial suggestions, Camille Gautier, Claire Bertrand and Anna Almeida (Morillon lab, Institut Curie) and Sylvain Baulande for RNA-seq (NGS platform, Institut Curie), Audrey Rapinat and David Gentien for NanoString experiments (Genomic Platform, Institut Curie), Cedric Saule and Jeremy Le Coz for a DE-kupl run (Gautheret lab, I2BC). TCGA RNA-seq data were downloaded from the dbGaP website under authorization phs000178/GRU to D.G..

## AUTHORS’ CONTRIBUTIONS

Conceptualization: D.G., A.M..

Funding acquisition: A.M., D.G..

Data acquisition: M.P..

Clinical samples collection and classification: Y.A., V.F., A.T..

Data analysis and curation: M.D., Mé.G., Ma.G., N.H.N., Z.S., M.P..

Interpretation: Y.A., D.G., V.F., A. L.-V., A.M., M.P., A.T..

Writing, review, and/or revision of the manuscript: D.G., A.M., M.P., Mé.G..

Study supervision: D.G., A.M., M.P..

## SUPPLEMENTAL MATERIAL

**Supplemental Table S1.** Clinico-pathological characteristics and recurrence status of the prostate specimens used for the total stranded RNA-sequencing (PAIR, *Discovery Set*).

**Supplemental Table S2.** DE-kupl contigs, PCA3 and housekeeping protein-coding genes for RNA expression measurements by the NanoString nCounter assay.

**Supplemental Table S3.** Clinico-pathological characteristics, risk classification and recurrence status of the prostate specimens used in NanoString (PAIR, *Selection Set*).

**Supplemental Table S4.** PCA3 and DE-kupl contigs expression measurements by the NanoString nCounter assay in 144 specimens of the PAIR cohort (*Selection Set*). Normalized expression of each probe was calculated as a ratio of the raw value to the mean expression of three housekeeping genes (*GPATCH3, ZNF2, ZNF346*).

**Supplemental Table S5.** Mean and Fold Change of expression of PCA3 and DE-kupl contigs in prostate normal and tumor specimens measured by NanoString across 144 prostate specimens of the *Selection Set*.

**Supplemental Table S6.** PCA3 and DE-kupl contigs expression quantification assessed by the total stranded RNA-seq in 24 prostate specimens from the PAIR cohort (*Discovery Set*).

**Supplemental Table S7.** Clinico-pathological characteristics and recurrence status of the prostate specimens from TCGA-PRAD cohort (*Validation Set*).

**Supplemental Table S8.** PCA3 and DE-kupl contigs quantification of expression assessed by the poly(A)+ unstranded RNA-seq in 557 prostate specimens from the TCGA-PRAD cohort (*Validation Set*).

**Supplemental Table S9.** Mean and Fold Change expression of PCA3, DE-kupl contigs and housekeeping genes in low-risk (LR) and high-risk (HR) tumors and recurrence negative (NO) and positive (YES) specimens of the PAIR cohort (*Selection Set*).

**Figure S1.**
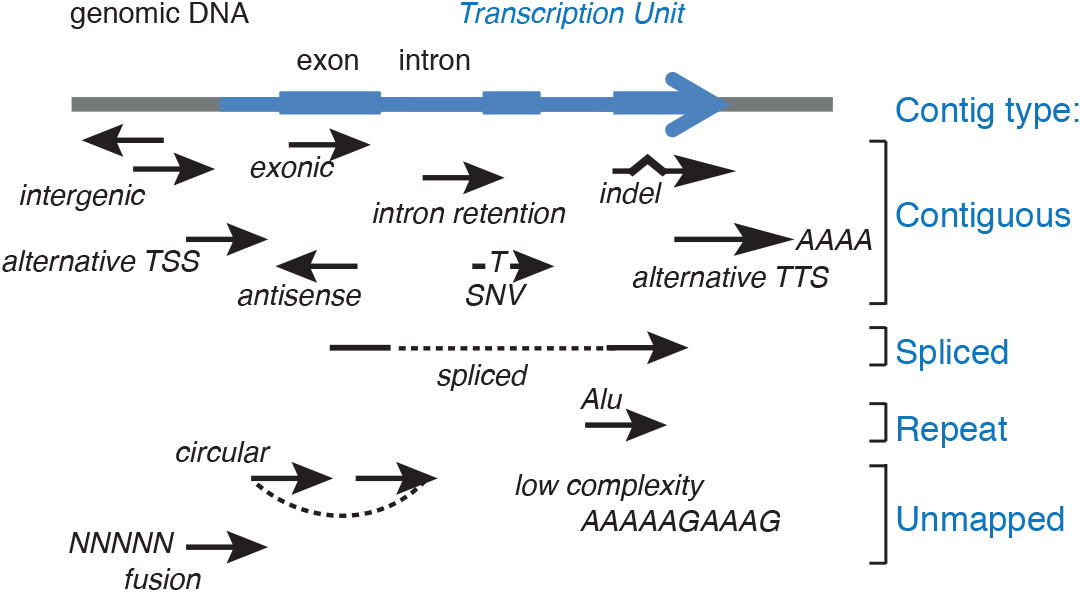
DE-kupl contigs assignment to contiguous, spliced, repeat and unmapped categories according to their genomic location outside or within annotated transcription units (blue). Each black arrow represents a contig.

**Figure S2.**
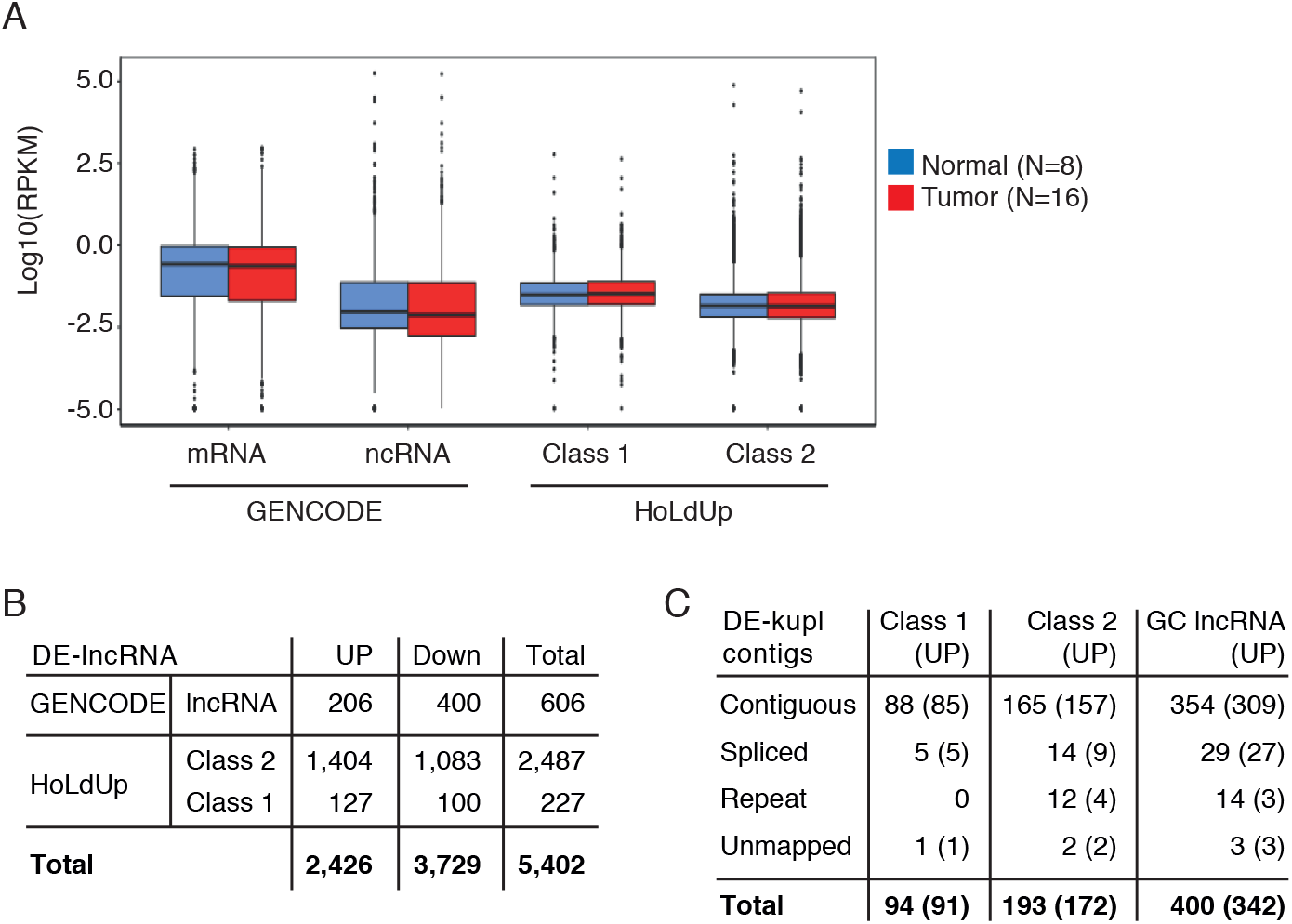
Expression of GENCODE and HoLdUp annotated lncRNAs in the *Discovery Set*. **(A)** Box-plot of Log10(RPKM) of mRNAs (N=21,330), lncRNAs (N=14,533) from the GENCODE annotation, and Class 1 (N=2,967) and Class 2 (N=168,163) TUs assembled by HoLdUp. Expression is measured in RPKM (Reads per kilo base per million mapped reads) by total stranded RNA-seq across 8 normal (bleu) and 16 tumor specimens (red). **(B)** Catalog of DE lncRNAs identified by DESeq within GENCODE lncRNAs and HoLdUp annotated TUs. **(C)** Intersection of DE-kupl contigs with HoLdUp and GENCODE annotated lncRNAs including those embedded into up-regulated transcripts (UP).

**Figure S3.**
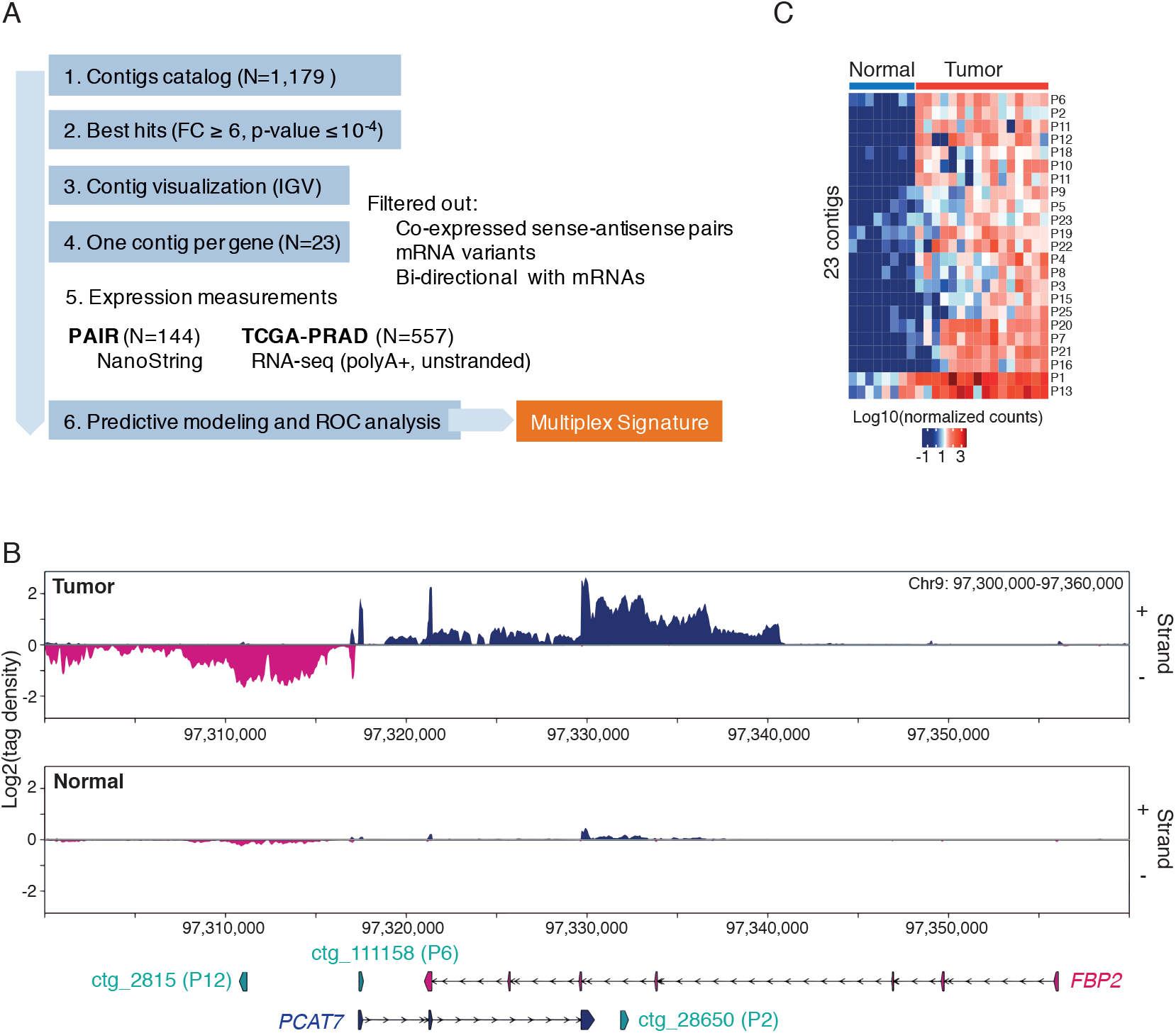
Selection of the most potent DE-kupl contigs. **(A)** Experimental rationale for contigs selection and validation of diagnostic potency. **(B)** RNA-seq profiling by VING along plus (+) and minus (-) strands of chr9: 97,300,000-97,360,000 in tumor and normal prostate specimens: the DE-kupl contig P6 (ctg_111158) assigned to PCAT7 and P2 (ctg_28650) antisense to the *FBP2* gene. Arrow-lines represent introns, rectangles - exons. **(C)** Heatmap representing the expression level and unsupervised clustering of the selected 23 DE-kupl contigs across prostate cancer specimens of the *Discovery Set*.

**Figure S4.**
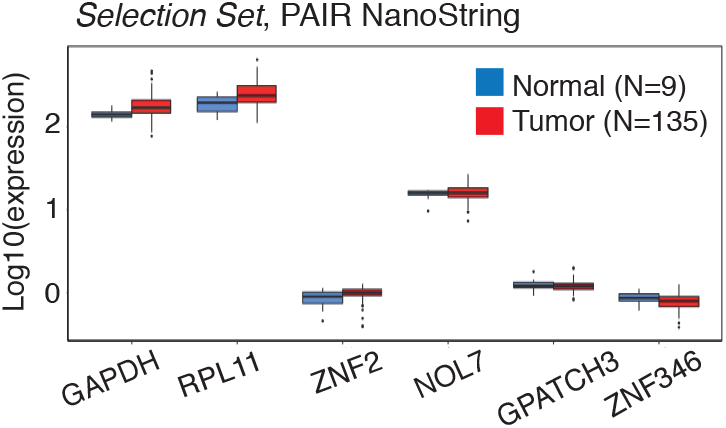
Box-plot of Log10(counts) of housekeeping protein-coding genes in 9 normal and 135 tumor specimens of the PAIR cohort (*Selection Set*) by the NanoString nCounter assay.

**Figure S5.**
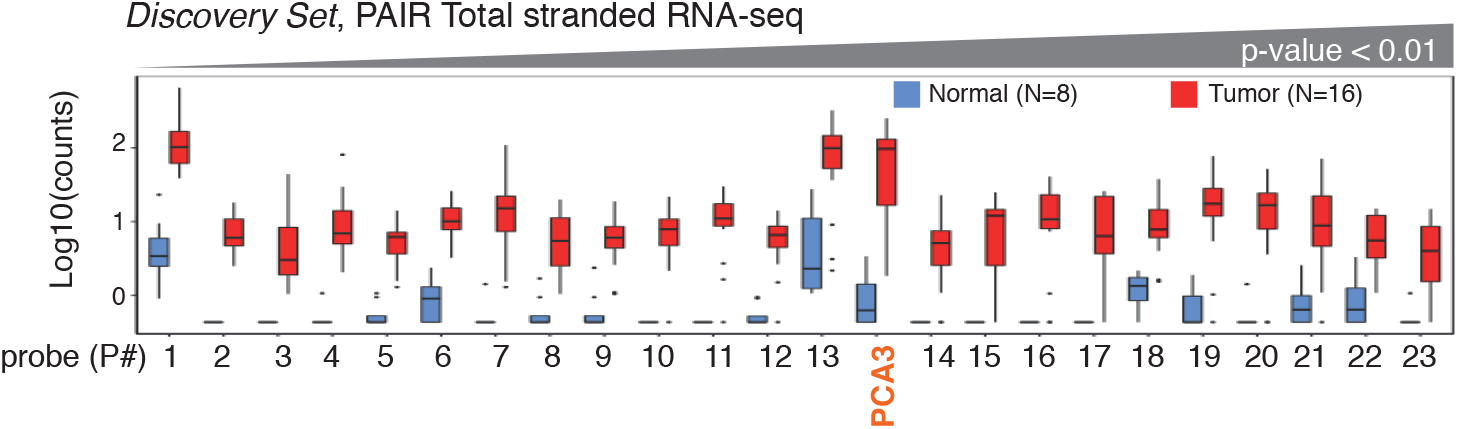
Box plot of PCA3 and DE-kupl contigs expression in the *Discovery Set* measured by the total stranded RNA-seq. Contigs are ordered by increasing adjusted p-values.

**Figure S6.**
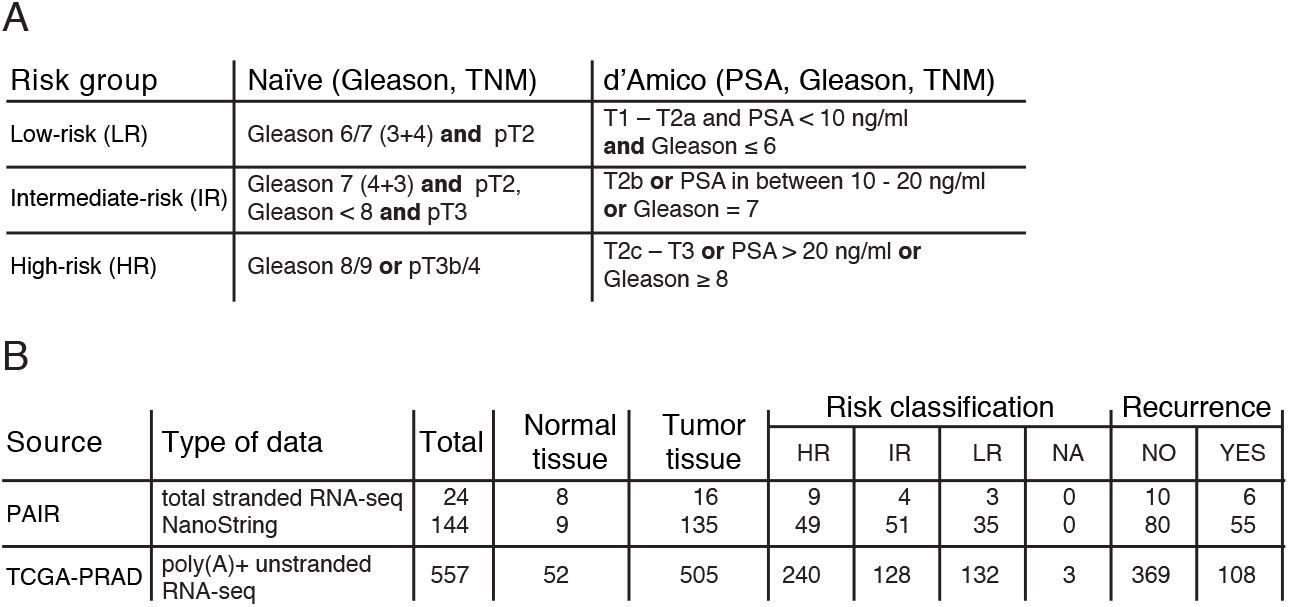
Risk classification of prostate tumors according to their clinico-pathological features. **(A)** Risk prognosis classification criteria according to D’Amico and the PSA independent naïve indexing. **(B)** Risk classification and recurrence status of prostate specimens from TCGA-PRAD and PAIR cohorts used in this study. PSA=prostate specific antigen; TNM=tumor, node, metastasis; HR=high-risk, IR=intermediate-risk, LR=low-risk tumors.

**Figure S7.**
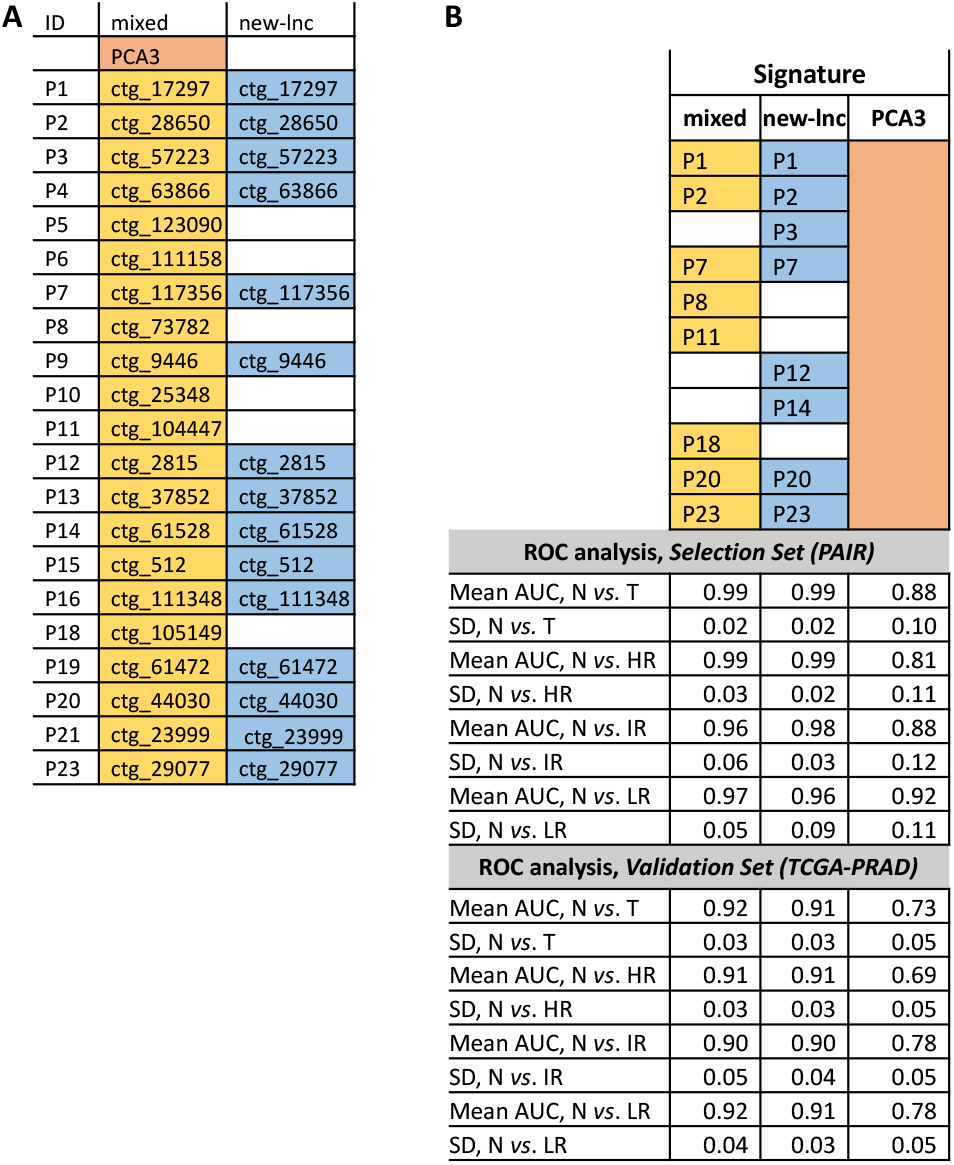
Predictive modeling and clinical performance analysis of DE-kupl contigs. **(A)** LASSO processed list of probes. **(B)** PCA3, mixed and new-lnc RNA signatures and their performance (mean and standard deviation of AUCs) in the PAIR (*Selection Set*) and the TCGA-PRAD (*Validation Set*) datasets. N = normal tissue, T = tumor tissue, HR = high-risk, IR = intermediate-risk, LR = low-risk tumors; AUC = area under the curve; SD = standard deviation.

**Figure S8.**
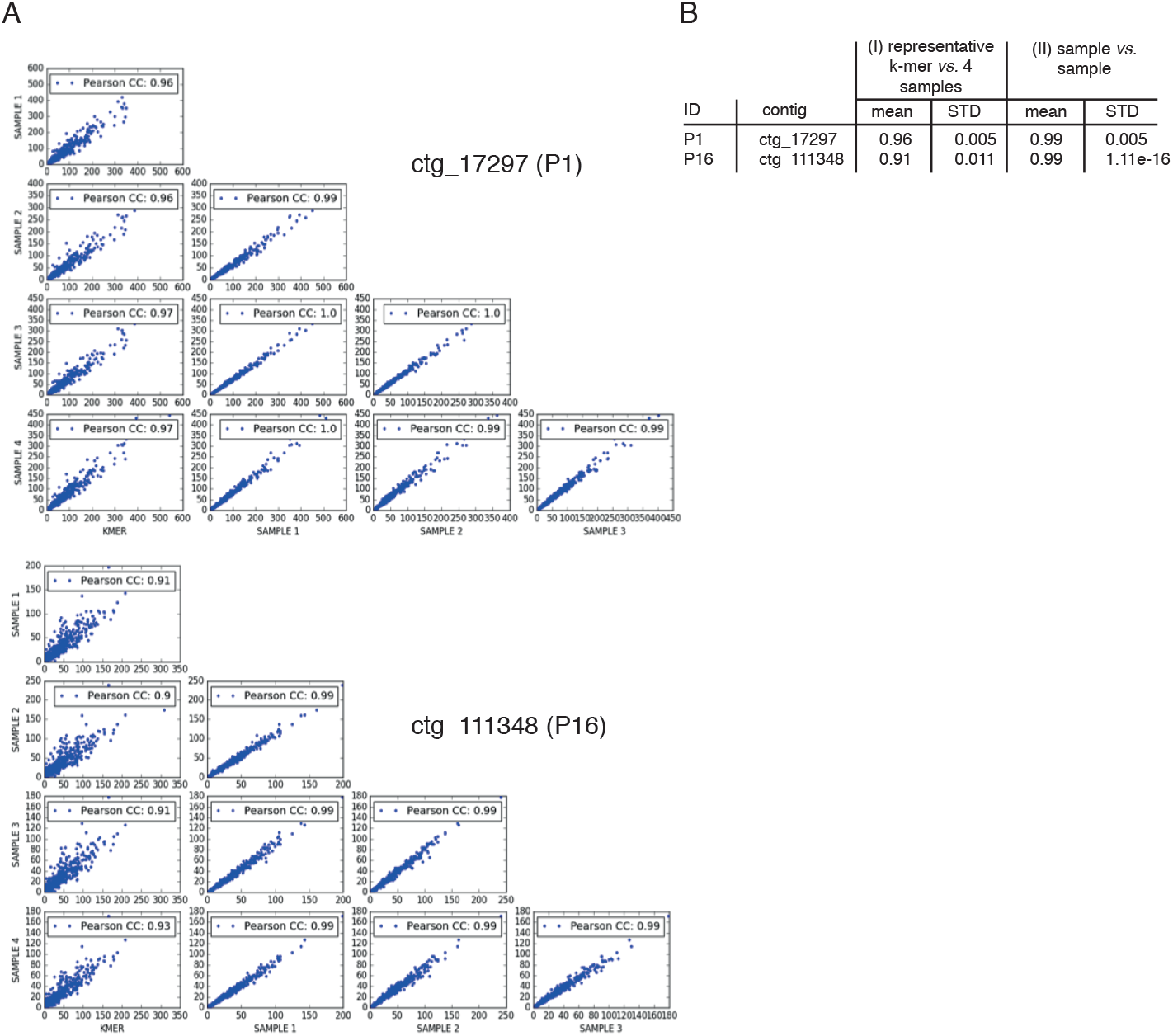
Contig expression measurements in TCGA-PRAD datasets. (A) Stability of k-mer counts for contigs P1 (ctg_17297) and P16 (ctg_111348) across the TCGA-PRAD dataset. **(B)** Pearson correlations between counts of representative k-mers and sampled k-mers from the same contig: for each contig, in the TCGA-PRAD datasets (N=557) the number of occurrences of (I) the representative DE-kupl k-mer and (II) of four sets of 10 k-mers sampled at regular distance along the length of the contig. Each sample (noted SAMPLE1-4) was obtained by changing the starting position of the first k-mer.

