## Supplemental Table S1 for "Blind exploration of the unreferenced transcriptome reveals novel RNAs for prostate cancer diagnosis"

Table S1. Clinico-pathological characteristics and recurrence status of the prostate specimens us

| Sample_ID | Tissue type | Gleason score | TMN score | Risk group | Recurrence |
|---|---|---|---|---|---|
| HMN_PC_102 | tumor | 6 (3 + 3) | pT2a | low | no |
| HMN_PC_228 | tumor | 6 (3 + 3) | pT2a | low | no |
| HMN_PC_089 | tumor | 7 (3 + 4) | pT2c | low | no |
| HMN_PC_167 | tumor | 8 (4 + 4) | pT2c | high | no |
| HMN_PC_077 | tumor | 7 (3 + 4) | pT3a | intermediate | no |
| HMN_PC_092 | tumor | 7 (4 + 3) | pT3a | intermediate | no |
| HMN_PC_124 | tumor | 8 (4 + 4) | pT3a | high | yes |
| HMN_PC_131 | tumor | 8 (4 + 4) | pT3a | high | yes |
| HMN_PC_195 | tumor | 8 (4 + 4) | pT3a | high | no |
| HMN_PC_139 | tumor | 6 (3 + 3) | pT3a | intermediate | yes |
| HMN_PC_033 | tumor | 7 (3 + 4) | pT3a | intermediate | no |
| HMN_PC_085 | tumor | 7 (4 + 3) | pT3b | high | no |
| HMN_PC_121 | tumor | 8 (4 + 4) | pT3b | high | yes |
| HMN_PC_009 | tumor | 8 (4 + 4) | pT3b | high | no |
| HMN_PC_148 | tumor | 8 (4 + 4) | pT4 | high | yes |
| HMN_PC_130 | tumor | 6 (3 + 3) | pT4 | high | yes |
| HMN_PC_78(-) | normal | | | | |
| HMN_PC_093(-) | normal | | | | |
| HMN_PC_125(-) | normal | | | | |
| HMN_PC_132(-) | normal | | | | |
| HMN_PC_149(-) | normal | | | | |
| HMN_PC_202(-) | normal | | | | |
| HMN_PC_086(-) | normal | | | | |
| HMN_PC_061(-) | normal | | | | |

pecimens used for the total stranded RNA-sequencing (PAIR, Discovery Set).
