## Supplemental Table S2 for "Blind exploration of the unreferenced transcriptome reveals novel RNAs for prostate cancer diagnosis"

**Table S2.** DE-kupl contigs, PCA3 and housekeeping protein-coding genes for RNA expression measurements by the NanoString nCounter assay.

| | | RNA-seq counts | | DE-kupl | | HUGO_ID | | |
|---|---|---|---|---|---|---|---|---|
| probe_ID | contig_ID | log2FC | pvalue | log2FC | pvalue | gene | paired_gene | Type of k-mer |
| P1 | ctg_17297 | 4,68 | 1,36E-06 | 3,90 | 3,86E-13 | | none | contiguous |
| P2 | ctg_28650 | Inf | 3,72E-05 | 4,46 | 6,80E-12 | | FBP2 | contiguous |
| P3 | ctg_57223 | Inf | 3,72E-05 | 3,87 | 2,23E-08 | | none | contiguous |
| P4 | ctg_63866 | 6,66 | 4,13E-05 | 4,03 | 1,15E-09 | | PDLIM5 | contiguous |
| P5 | ctg_123090 | 4,54 | 4,46E-05 | 3,33 | 2,96E-08 | AC004066.3 | none | contiguous |
| P6 | ctg_111158 | 3,79 | 4,97E-05 | 3,21 | 3,26E-10 | PCAT7 | FBP2 | contiguous |
| P7 | ctg_117356 | 6,85 | 5,34E-05 | 4,35 | 9,73E-12 | | snoU13 | unmapped |
| P8 | ctg_73782 | 4,51 | 5,74E-05 | 3,32 | 8,57E-08 | LINC01006 | none | contiguous |
| P9 | ctg_9446 | 4,05 | 7,38E-05 | 3,07 | 1,37E-07 | | none | contiguous |
| P10 | ctg_25348 | Inf | 9,11E-05 | 4,33 | 6,66E-11 | CTBP1-AS | CTBP1 | contiguous |
| P11 | ctg_104447 | Inf | 9,11E-05 | 4,68 | 5,55E-13 | RP11-627G | none | contiguous |
| P12 | ctg_2815 | 4,88 | 1,44E-04 | 3,46 | 7,46E-09 | | none | contiguous |
| P13 | ctg_37852 | 3,86 | 1,85E-04 | 2,95 | 2,85E-06 | | ABCC4 | contiguous |
| PCA3 | - | 6,42 | 2,09E-04 | - | - | PCA3 | PRUNE2 | - |
| P14 | ctg_61528 | Inf | 2,10E-04 | 3,82 | 2,91E-08 | | TPO | contiguous |
| P15 | ctg_512 | Inf | 2,10E-04 | 4,07 | 3,18E-09 | | PXDN | spliced |
| P16 | ctg_111348 | Inf | 2,10E-04 | 4,34 | 1,71E-10 | | DLX1 | contiguous |
| P17 | ctg_36195 | Inf | 2,10E-04 | 4,07 | 3,21E-09 | | none | repeat |
| P18 | ctg_105149 | 3,32 | 2,14E-04 | 2,74 | 3,86E-07 | PCAT1 | none | contiguous |
| P19 | ctg_61472 | 5,51 | 2,75E-04 | 3,85 | 1,15E-09 | | AP006748.1 | contiguous |
| P20 | ctg_44030 | 6,57 | 3,04E-04 | 4,12 | 2,92E-10 | | none | contguous |
| P21 | ctg_23999 | 4,70 | 3,21E-04 | 3,40 | 8,80E-08 | | FBXL7 | contiguous |
| P22 | ctg_119680 | 2,96 | 3,30E-04 | 2,38 | 1,46E-05 | | none | contiguous |
| P23 | ctg_29077 | 5,38 | 7,42E-04 | 3,05 | 1,64E-05 | | AC011523.2 | contiguous |
| RPL11 | - | - | - | - | - | RPL11 | none | - |
| GAPDH | - | - | - | - | - | GAPDH | none | - |
| NOL7 | - | - | - | - | - | NOL7 | none | - |
| GPATCH3 | - | - | - | - | - | GPATCH3 | none | - |
| ZNF2 | - | - | - | - | - | ZNF2 | none | - |
| ZNF346 | - | - | - | - | - | ZNF346 | none | - |
