## Supplemental Table S3 for "Blind exploration of the unreferenced transcriptome reveals novel RNAs for prostate cancer diagnosis"

**Table S3. Clinico-pathological characteristics, risk classification and recurrence status of the prostate specimens used in NanoString (PAIR, Selection Set).**

| ID_sample | Tissue type | TNM | Gleason score | Recurrence | Risk group |
|---|---|---|---|---|---|
| HMN_PC_001 | Tumor | pT2c | 7 (4 + 3) | no | intermediate risk |
| HMN_PC_010 | Tumor | pT3a | 7 (4 + 3) | no | intermediate risk |
| HMN_PC_013 | Tumor | pT2c | 6 (3 + 3) | no | low risk |
| HMN_PC_014 | Tumor | pT3a | 7 (3 + 4) | yes | intermediate risk |
| HMN_PC_016 | Tumor | pT3b | 9 (4 + 5) | yes | high risk |
| HMN_PC_019 | Tumor | pT2c | 7 (4 + 3) | yes | intermediate risk |
| HMN_PC_020 | Tumor | pT3a | 7 (3 + 4) | no | intermediate risk |
| HMN_PC_021 | Tumor | pT3a | 7 (3 + 4) | no | intermediate risk |
| HMN_PC_022 | Tumor | pT3a | 8 (4 + 4) | yes | high risk |
| HMN_PC_023 | Tumor | pT3a | 7 (4 + 3) | no | intermediate risk |
| HMN_PC_024 | Tumor | pT2c | 6 (3 + 3) | no | low risk |
| HMN_PC_026 | Tumor | pT3a | 7 (4 + 3) | no | intermediate risk |
| HMN_PC_027 | Tumor | pT2c | 7 (3 + 4) | no | low risk |
| HMN_PC_028 | Tumor | pT3a | 8 (4 + 4) | yes | high risk |
| HMN_PC_030 | Tumor | pT3b | 7 (3 + 4) | no | high risk |
| HMN_PC_031 | Tumor | pT2c | 7 (3 + 4) | no | low risk |
| HMN_PC_032 | Tumor | pT2c | 7 (4 + 3) | no | intermediate risk |
| HMN_PC_033 | Tumor | pT3a | 7 (3 + 4) | no | intermediate risk |
| HMN_PC_036 | Tumor | pT4 | 7 (4 + 3) | no | high risk |
| HMN_PC_037 | Tumor | pT3a | 7 (4 + 3) | yes | intermediate risk |
| HMN_PC_038 | Tumor | pT3b | 7 (4 + 3) | no | high risk |
| HMN_PC_040 | Tumor | pT3a | 7 (4 + 3) | no | intermediate risk |
| HMN_PC_041 | Tumor | pT3a | 7 (4 + 3) | no | intermediate risk |
| HMN_PC_043 | Tumor | pT2c | 6 (3 + 3) | no | low risk |
| HMN_PC_044 | Tumor | pT2a | 7 (3 + 4) | no | low risk |
| HMN_PC_045 | Tumor | pT2c | 7 (4 + 3) | no | intermediate risk |
| HMN_PC_047 | Tumor | pT3b | 7 (4 + 3) | no | high risk |
| HMN_PC_049 | Tumor | pT3b | 8 (4 + 4) | yes | high risk |
| HMN_PC_050 | Tumor | pT3a | 7 (4 + 3) | yes | intermediate risk |
| HMN_PC_051 | Tumor | pT2c | 6 (3 + 3) | no | low risk |
| HMN_PC_052 | Tumor | pT3a | 8 (4 + 4) | yes | high risk |
| HMN_PC_056 | Tumor | pT3a | 8 (4 + 4) | yes | high risk |
| HMN_PC_057 | Tumor | pT3b | 8 (4 + 4) | yes | high risk |
| HMN_PC_058(-) | Normal | - | - | - | - |
| HMN_PC_059(-) | Normal | - | - | - | - |
| HMN_PC_060(-) | Normal | - | - | - | - |
| HMN_PC_063 | Tumor | pT3a | 8 (4 + 4) | yes | high risk |
| HMN_PC_066 | Tumor | pT3b | 7 (4 + 3) | no | high risk |
| HMN_PC_067 | Tumor | pT2a | 8 (4 + 4) | yes | high risk |
| HMN_PC_068 | Tumor | pT3b | 9 (4 + 5) | yes | high risk |
| HMN_PC_070 | Tumor | pT3a | 7 (4 + 3) | yes | intermediate risk |
| HMN_PC_071 | Tumor | pT3b | 7 (4 + 3) | yes | high risk |
| HMN_PC_072 | Tumor | pT3b | 9 (4 + 5) | no | high risk |
| HMN_PC_075 | Tumor | pT3b | 7 (4 + 3) | no | high risk |
| HMN_PC_076 | Tumor | pT2c | 8 (4 + 4) | yes | high risk |

| | | | | | |
|---|---|---|---|---|---|
| HMN_PC_077.1 | Tumor | pT3a | 7 (3 + 4) | no | intermediate risk |
| HMN_PC_077.2 | Tumor | pT3a | 7 (4 + 3) | yes | intermediate risk |
| HMN_PC_078(-) | Normal | - | - | - | - |
| HMN_PC_081 | Tumor | pT3a | 7 (3 + 4) | no | intermediate risk |
| HMN_PC_082 | Tumor | pT2c | 7 (3 + 4) | no | low risk |
| HMN_PC_084 | Tumor | pT2c | 7 (3 + 4) | no | low risk |
| HMN_PC_085 | Tumor | pT3b | 7 (4 + 3) | no | high risk |
| HMN_PC_086(-) | Normal | - | - | - | - |
| HMN_PC_087 | Tumor | pT3a | 7 (4 + 3) | no | intermediate risk |
| HMN_PC_088(-) | Normal | - | - | - | - |
| HMN_PC_089 | Tumor | pT2c | 7 (3 + 4) | no | low risk |
| HMN_PC_092 | Tumor | pT3a | 7 (4 + 3) | no | intermediate risk |
| HMN_PC_093(-) | Normal | - | - | - | - |
| HMN_PC_096 | Tumor | pT2a | 7 (3 + 4) | no | low risk |
| HMN_PC_097 | Tumor | pT2c | 7 (3 + 4) | no | low risk |
| HMN_PC_099 | Tumor | pT2c | 6 (3 + 3) | no | low risk |
| HMN_PC_100 | Tumor | pT3a | 7 (4 + 3) | no | intermediate risk |
| HMN_PC_101 | Tumor | pT2a | 6 (3 + 3) | no | low risk |
| HMN_PC_102 | Tumor | pT2a | 6 (3 + 3) | no | low risk |
| HMN_PC_103 | Tumor | pT3a | 7 (4 + 3) | no | intermediate risk |
| HMN_PC_104 | Tumor | pT3a | 7 (3 + 4) | no | intermediate risk |
| HMN_PC_105 | Tumor | pT2c | 7 (3 + 4) | no | low risk |
| HMN_PC_106 | Tumor | pT3a | 7 (3 + 4) | no | intermediate risk |
| HMN_PC_110 | Tumor | pT3a | 7 (4 + 3) | yes | intermediate risk |
| HMN_PC_111 | Tumor | pT2c | 6 (3 + 3) | no | low risk |
| HMN_PC_113 | Tumor | pT2c | 7 (3 + 4) | no | low risk |
| HMN_PC_114 | Tumor | pT2b | 6 (3 + 3) | no | low risk |
| HMN_PC_115 | Tumor | pT2c | 6 (3 + 3) | no | low risk |
| HMN_PC_117 | Tumor | pT3a | 6 (3 + 3) | no | intermediate risk |
| HMN_PC_118 | Tumor | pT3a | 7 (4 + 3) | no | intermediate risk |
| HMN_PC_119 | Tumor | pT2c | 7 (3 + 4) | no | low risk |
| HMN_PC_121 | Tumor | pT3b | 8 (4 + 4) | yes | high risk |
| HMN_PC_124 | Tumor | pT3a | 8 (4 + 4) | yes | high risk |
| HMN_PC_125(-) | Normal | - | - | - | - |
| HMN_PC_126 | Tumor | pT4 | 7 (4 + 3) | yes | high risk |
| HMN_PC_127 | Tumor | pT3a | 8 (4 + 4) | yes | high risk |
| HMN_PC_128 | Tumor | pT3a | 7 (4 + 3) | yes | intermediate risk |
| HMN_PC_129 | Tumor | pT2c | 6 (3 + 3) | no | low risk |
| HMN_PC_130 | Tumor | pT4 | 6 (3 + 3) | yes | high risk |
| HMN_PC_131 | Tumor | pT3a | 8 (4 + 4) | yes | high risk |
| HMN_PC_134 | Tumor | pT3a | 8 (4 + 4) | yes | high risk |
| HMN_PC_135 | Tumor | pT3a | 7 (4 + 3) | yes | intermediate risk |
| HMN_PC_137 | Tumor | pT3b | 9 (4 + 5) | yes | high risk |
| HMN_PC_138 | Tumor | pT2c | 7 (3 + 4) | yes | low risk |
| HMN_PC_139 | Tumor | pT3a | 6 (3 + 3) | yes | intermediate risk |
| HMN_PC_140 | Tumor | pT3b | 7 (4 + 3) | yes | high risk |
| HMN_PC_141 | Tumor | pT3a | 8 (4 + 4) | yes | high risk |
| HMN_PC_142 | Tumor | pT3a | 8 (4 + 4) | yes | high risk |

| | | | | | |
|---|---|---|---|---|---|
| HMN_PC_143 | Tumor | pT3b | 8 (4 + 4) | yes | high risk |
| HMN_PC_145 | Tumor | pT3b | 9 (4 + 5) | no | high risk |
| HMN_PC_146 | Tumor | pT3b | 7 (4 + 3) | yes | high risk |
| HMN_PC_147 | Tumor | pT2c | 7 (3 + 4) | yes | low risk |
| HMN_PC_148 | Tumor | pT4 | 8 (4 + 4) | yes | high risk |
| HMN_PC_149(-) | Normal | - | - | - | - |
| HMN_PC_152 | Tumor | pT3a | 7 (3 + 4) | no | intermediate risk |
| HMN_PC_153 | Tumor | pT3a | 6 (3 + 3) | no | intermediate risk |
| HMN_PC_156 | Tumor | pT2c | 6 (3 + 3) | no | low risk |
| HMN_PC_157 | Tumor | pT3b | 7 (4 + 3) | no | high risk |
| HMN_PC_158 | Tumor | pT2c | 7 (3 + 4) | no | low risk |
| HMN_PC_160 | Tumor | pT3a | 7 (3 + 4) | no | intermediate risk |
| HMN_PC_161 | Tumor | pT2c | 6 (3 + 3) | no | low risk |
| HMN_PC_162 | Tumor | pT3a | 7 (4 + 3) | no | intermediate risk |
| HMN_PC_164 | Tumor | pT3a | 7 (4 + 3) | no | intermediate risk |
| HMN_PC_166 | Tumor | pT3a | 7 (4 + 3) | no | intermediate risk |
| HMN_PC_167 | Tumor | pT2c | 8 (4 + 4) | no | high risk |
| HMN_PC_168 | Tumor | pT2a | 7 (4 + 3) | yes | intermediate risk |
| HMN_PC_169 | Tumor | pT3a | 7 (4 + 3) | no | intermediate risk |
| HMN_PC_170 | Tumor | pT3a | 7 (4 + 3) | yes | intermediate risk |
| HMN_PC_171 | Tumor | pT2c | 7 (3 + 4) | no | low risk |
| HMN_PC_172 | Tumor | pT3a | 8 (4 + 4) | yes | high risk |
| HMN_PC_173 | Tumor | pT2c | 6 (3 + 3) | no | low risk |
| HMN_PC_174 | Tumor | pT2c | 7 (3 + 4) | no | low risk |
| HMN_PC_209 | Tumor | pT3a | 7 (3 + 4) | yes | intermediate risk |
| HMN_PC_217 | Tumor | pT3b | 7 (4 + 3) | yes | high risk |
| HMN_PC_218 | Tumor | pT3a | 7 (4 + 3) | yes | intermediate risk |
| HMN_PC_219 | Tumor | pT3a | 8 (4 + 4) | yes | high risk |
| HMN_PC_220 | Tumor | pT4 | 8 (4 + 4) | yes | high risk |
| HMN_PC_221 | Tumor | pT3b | 7 (4 + 3) | yes | high risk |
| HMN_PC_222 | Tumor | pT2c | 8 (3 + 5) | no | high risk |
| HMN_PC_223 | Tumor | pT4 | 8 (4 + 4) | yes | high risk |
| HMN_PC_224 | Tumor | pT3b | 8 (4 + 4) | yes | high risk |
| HMN_PC_225 | Tumor | pT3a | 8 (4 + 4) | yes | high risk |
| HMN_PC_226 | Tumor | pT3a | 8 (4 + 4) | yes | high risk |
| HMN_PC_227 | Tumor | pT2c | 7 (4 + 3) | no | intermediate risk |
| HMN_PC_228 | Tumor | pT2a | 6 (3 + 3) | no | low risk |
| HMN_PC_229 | Tumor | pT3a | 7 (4 + 3) | no | intermediate risk |
| HMN_PC_232 | Tumor | pT3a | 7 (4 + 3) | yes | intermediate risk |
| HMN_PC_233 | Tumor | pT2c | 7 (3 + 4) | no | low risk |
| HMN_PC_234 | Tumor | pT3a | 8 (4 + 4) | yes | high risk |
| HMN_PC_235 | Tumor | pT2c | 6 (3 + 3) | no | low risk |
| HMN_PC_236 | Tumor | pT3a | 7 (4 + 3) | no | intermediate risk |
| HMN_PC_239 | Tumor | pT2c | 6 (3 + 3) | no | low risk |
| HMN_PC_241 | Tumor | pT3a | 7 (4 + 3) | no | intermediate risk |
| HMN_PC_242 | Tumor | pT3a | 7 (3 + 4) | no | intermediate risk |
| HMN_PC_243 | Tumor | pT3a | 7 (4 + 3) | no | intermediate risk |
| HMN_PC_244 | Tumor | pT4 | 8 (4 + 4) | yes | high risk |

| | | | | | |
|---|---|---|---|---|---|
| HMN_PC_245 | Tumor | pT3a | 7 (4 + 3) | yes | intermediate risk |
| HMN_PC_246 | Tumor | pT3a | 7 (4 + 3) | no | intermediate risk |
| HMN_PC_247 | Tumor | pT2c | 6 (3 + 3) | no | low risk |
