## Supplemental Table S4 for "Blind exploration of the unreferenced transcriptome reveals novel RNAs for prostate cancer diagnosis"

**Table S4. PCA3 and DE-kupl contigs expression measurements by NanoString nCounter assay in 144 specimens of the PAIR cohort (Selection Set). Normalized expression of each probe was calculated as a ratio of the raw**

| ID_sample | Tissue | ctg_123090 | ctg_25348 | ctg_73782 | ctg_104447 | PCA3 | ctg_105149 | ctg_111158 |
|---|---|---|---|---|---|---|---|---|
| HMN_PC_001 | tumor | 0,43 | 0,84 | 6,08 | 1,90 | 28,06 | 1,89 | 2,15 |
| HMN_PC_010 | tumor | 0,10 | 0,18 | 0,17 | 0,54 | 20,14 | 1,36 | 1,77 |
| HMN_PC_013 | tumor | 0,11 | 0,27 | 2,65 | 0,60 | 21,05 | 0,37 | 1,08 |
| HMN_PC_014 | tumor | 0,12 | 0,39 | 3,94 | 0,17 | 35,01 | 0,17 | 0,65 |
| HMN_PC_016 | tumor | 0,10 | 0,34 | 3,37 | 1,37 | 15,16 | 1,30 | 0,86 |
| HMN_PC_019 | tumor | 0,52 | 2,29 | 0,80 | 0,22 | 22,95 | 3,06 | 2,82 |
| HMN_PC_020 | tumor | 0,26 | 0,43 | 4,23 | 0,48 | 15,25 | 1,55 | 5,64 |
| HMN_PC_021 | tumor | 0,30 | 0,16 | 1,19 | 1,15 | 13,16 | 1,29 | 0,87 |
| HMN_PC_022 | tumor | 0,16 | 0,18 | 0,32 | 0,23 | 1,34 | 2,45 | 0,86 |
| HMN_PC_023 | tumor | 0,14 | 0,11 | 1,22 | 1,38 | 8,50 | 0,52 | 2,15 |
| HMN_PC_024 | tumor | 0,18 | 0,06 | 4,20 | 2,27 | 9,97 | 0,49 | 0,93 |
| HMN_PC_026 | tumor | 0,51 | 0,24 | 0,97 | 3,50 | 22,19 | 3,54 | 2,46 |
| HMN_PC_027 | tumor | 0,17 | 0,52 | 7,17 | 0,97 | 16,19 | 0,65 | 1,39 |
| HMN_PC_028 | tumor | 0,38 | 0,76 | 3,43 | 0,48 | 11,18 | 0,48 | 2,81 |
| HMN_PC_030 | tumor | 0,45 | 0,79 | 0,94 | 0,08 | 37,51 | 0,72 | 5,61 |
| HMN_PC_031 | tumor | 0,20 | 0,97 | 0,43 | 1,12 | 21,32 | 4,79 | 2,18 |
| HMN_PC_032 | tumor | 0,14 | 0,63 | 5,66 | 0,13 | 9,40 | 0,23 | 3,64 |
| HMN_PC_033 | tumor | 0,12 | 0,92 | 0,51 | 2,20 | 17,19 | 1,73 | 0,82 |
| HMN_PC_036 | tumor | 0,03 | 0,28 | 0,78 | 1,31 | 18,64 | 0,82 | 0,68 |
| HMN_PC_037 | tumor | 0,08 | 0,15 | 2,79 | 0,26 | 23,40 | 0,24 | 0,90 |
| HMN_PC_038 | tumor | 0,17 | 0,65 | 0,18 | 2,19 | 2,92 | 0,71 | 2,69 |
| HMN_PC_040 | tumor | 0,01 | 0,01 | 2,08 | 0,46 | 0,51 | 0,57 | 0,79 |
| HMN_PC_041 | tumor | 0,24 | 0,11 | 8,98 | 0,74 | 28,16 | 1,05 | 0,99 |
| HMN_PC_043 | tumor | 0,40 | 1,76 | 0,58 | 10,35 | 11,37 | 2,44 | 0,96 |
| HMN_PC_044 | tumor | 0,32 | 0,42 | 0,21 | 0,28 | 2,08 | 0,77 | 1,69 |
| HMN_PC_045 | tumor | 0,08 | 0,29 | 0,18 | 0,66 | 2,57 | 0,55 | 1,98 |
| HMN_PC_047 | tumor | 0,04 | 0,13 | 1,46 | 0,14 | 35,75 | 0,61 | 2,01 |
| HMN_PC_049 | tumor | 0,17 | 0,24 | 2,61 | 0,33 | 0,31 | 0,28 | 1,92 |
| HMN_PC_050 | tumor | 1,32 | 0,32 | 22,44 | 3,00 | 4,06 | 1,33 | 1,44 |
| HMN_PC_051 | tumor | 0,77 | 0,76 | 0,68 | 1,12 | 41,45 | 1,37 | 1,93 |
| HMN_PC_052 | tumor | 0,62 | 0,51 | 1,51 | 0,51 | 6,45 | 1,66 | 2,58 |
| HMN_PC_056 | tumor | 0,27 | 0,39 | 0,72 | 0,45 | 16,22 | 9,64 | 4,12 |
| HMN_PC_057 | tumor | 0,15 | 1,53 | 0,36 | 1,31 | 0,18 | 0,13 | 2,60 |
| HMN_PC_058(-) | normal | 0,02 | 0,06 | 0,10 | 0,19 | 0,88 | 0,07 | 0,02 |
| HMN_PC_059(-) | normal | 0,03 | 0,09 | 0,03 | 0,04 | 0,14 | 0,12 | 0,03 |
| HMN_PC_060(-) | normal | 0,02 | 0,09 | 0,10 | 0,02 | 0,17 | 0,04 | 0,35 |
| HMN_PC_063 | tumor | 0,17 | 0,29 | 6,19 | 1,07 | 2,31 | 0,44 | 2,96 |
| HMN_PC_066 | tumor | 0,21 | 0,73 | 0,36 | 0,05 | 35,19 | 0,55 | 2,01 |
| HMN_PC_067 | tumor | 0,35 | 0,35 | 0,21 | 0,01 | 1,62 | 5,52 | 0,73 |
| HMN_PC_068 | tumor | 0,49 | 1,00 | 2,88 | 0,19 | 0,23 | 1,33 | 5,25 |
| HMN_PC_070 | tumor | 0,67 | 0,40 | 5,89 | 3,30 | 0,64 | 1,42 | 2,61 |
| HMN_PC_071 | tumor | 0,39 | 0,15 | 2,01 | 1,54 | 16,16 | 3,27 | 3,42 |
| HMN_PC_072 | tumor | 0,39 | 0,18 | 0,12 | 0,35 | 21,68 | 0,31 | 0,28 |
| HMN_PC_075 | tumor | 0,24 | 0,11 | 0,71 | 0,03 | 16,78 | 0,58 | 3,04 |
| HMN_PC_076 | tumor | 0,35 | 0,63 | 0,34 | 2,79 | 81,11 | 1,87 | 4,63 |
| HMN_PC_077.1 | tumor | 0,68 | 0,28 | 10,36 | 1,97 | 8,13 | 2,03 | 1,36 |
| HMN_PC_077.2 | tumor | 0,12 | 0,10 | 4,98 | 0,46 | 1,01 | 0,50 | 1,43 |
| HMN_PC_078(-) | normal | 0,22 | 0,04 | 0,18 | 0,12 | 0,64 | 0,16 | 0,21 |
| HMN_PC_081 | tumor | 0,32 | 0,39 | 5,89 | 1,96 | 21,82 | 2,43 | 1,17 |
| HMN_PC_082 | tumor | 0,62 | 1,89 | 0,99 | 8,32 | 12,76 | 2,17 | 4,18 |
| HMN_PC_084 | tumor | 0,35 | 0,10 | 2,25 | 0,37 | 4,54 | 0,82 | 0,34 |
| HMN_PC_085 | tumor | 0,80 | 1,26 | 9,46 | 0,39 | 1,85 | 0,39 | 5,93 |
| HMN_PC_086(-) | normal | 0,29 | 0,05 | 0,92 | 0,35 | 1,30 | 0,40 | 0,17 |
| HMN_PC_087 | tumor | 0,46 | 0,15 | 2,97 | 3,38 | 26,84 | 2,68 | 1,48 |
| HMN_PC_088(-) | normal | 0,12 | 0,12 | 0,34 | 0,18 | 1,35 | 0,19 | 0,17 |
| HMN_PC_089 | tumor | 0,74 | 1,98 | 7,29 | 4,11 | 27,50 | 1,18 | 1,79 |
| HMN_PC_092 | tumor | 0,19 | 0,29 | 6,89 | 1,68 | 13,16 | 1,03 | 4,05 |
| HMN_PC_093(-) | normal | 0,08 | 0,06 | 0,24 | 0,33 | 0,83 | 0,15 | 0,09 |
| HMN_PC_096 | tumor | 0,12 | 0,38 | 2,42 | 0,41 | 6,22 | 1,00 | 0,59 |

| ID_sample | Tissue | ctg_123090 | ctg_25348 | ctg_73782 | ctg_104447 | PCA3 | ctg_105149 | ctg_111158 |
|---|---|---|---|---|---|---|---|---|
| HMN_PC_097 | tumor | 0,27 | 0,36 | 5,47 | 0,65 | 51,87 | 0,81 | 2,24 |
| HMN_PC_099 | tumor | 0,19 | 0,16 | 0,92 | 0,73 | 8,18 | 0,66 | 1,94 |
| HMN_PC_100 | tumor | 0,39 | 0,17 | 4,16 | 1,10 | 8,32 | 0,70 | 1,72 |
| HMN_PC_101 | tumor | 0,33 | 0,35 | 4,16 | 0,54 | 5,64 | 0,39 | 0,30 |
| HMN_PC_102 | tumor | 0,70 | 0,07 | 5,92 | 2,93 | 16,67 | 1,81 | 0,54 |
| HMN_PC_103 | tumor | 1,14 | 0,42 | 9,88 | 1,51 | 48,94 | 4,05 | 6,22 |
| HMN_PC_104 | tumor | 0,40 | 1,10 | 0,34 | 3,07 | 26,24 | 1,65 | 1,11 |
| HMN_PC_105 | tumor | 0,56 | 0,45 | 6,19 | 0,82 | 2,76 | 1,68 | 2,46 |
| HMN_PC_106 | tumor | 0,87 | 0,44 | 17,06 | 2,63 | 16,24 | 5,06 | 2,55 |
| HMN_PC_110 | tumor | 0,52 | 0,34 | 12,52 | 2,58 | 10,56 | 2,59 | 3,25 |
| HMN_PC_111 | tumor | 0,14 | 0,13 | 5,27 | 0,40 | 27,58 | 0,38 | 2,36 |
| HMN_PC_113 | tumor | 0,53 | 1,31 | 0,72 | 2,19 | 15,49 | 1,57 | 3,58 |
| HMN_PC_114 | tumor | 0,22 | 0,33 | 7,28 | 1,52 | 20,42 | 1,43 | 1,08 |
| HMN_PC_115 | tumor | 0,26 | 0,40 | 1,36 | 0,50 | 26,76 | 0,40 | 1,23 |
| HMN_PC_117 | tumor | 0,38 | 0,24 | 4,59 | 1,91 | 21,26 | 2,05 | 1,54 |
| HMN_PC_118 | tumor | 0,21 | 0,22 | 0,56 | 0,63 | 0,09 | 0,53 | 0,25 |
| HMN_PC_119 | tumor | 0,49 | 0,13 | 3,47 | 1,56 | 54,92 | 1,83 | 4,16 |
| HMN_PC_121 | tumor | 0,75 | 0,41 | 2,94 | 0,59 | 0,65 | 1,42 | 5,11 |
| HMN_PC_124 | tumor | 0,26 | 0,09 | 0,51 | 2,46 | 1,85 | 1,02 | 5,21 |
| HMN_PC_125(-) | normal | 0,18 | 0,03 | 0,40 | 0,04 | 0,31 | 0,10 | 0,04 |
| HMN_PC_126 | tumor | 0,23 | 0,41 | 0,74 | 0,92 | 27,96 | 0,56 | 2,59 |
| HMN_PC_127 | tumor | 0,15 | 0,58 | 0,08 | 2,24 | 0,64 | 1,69 | 4,06 |
| HMN_PC_128 | tumor | 0,68 | 0,17 | 2,81 | 1,45 | 17,90 | 2,21 | 2,25 |
| HMN_PC_129 | tumor | 0,45 | 0,12 | 8,78 | 1,24 | 5,63 | 1,39 | 1,06 |
| HMN_PC_130 | tumor | 0,51 | 3,57 | 0,97 | 3,80 | 5,81 | 3,57 | 3,44 |
| HMN_PC_131 | tumor | 0,58 | 0,99 | 0,48 | 1,79 | 0,76 | 6,93 | 4,85 |
| HMN_PC_134 | tumor | 0,45 | 1,75 | 0,82 | 3,64 | 2,16 | 1,17 | 2,81 |
| HMN_PC_135 | tumor | 0,45 | 0,82 | 2,33 | 0,30 | 10,83 | 1,05 | 3,11 |
| HMN_PC_137 | tumor | 0,47 | 0,30 | 1,42 | 0,07 | 0,75 | 2,96 | 1,88 |
| HMN_PC_138 | tumor | 0,86 | 0,45 | 1,13 | 1,94 | 29,27 | 4,68 | 0,95 |
| HMN_PC_139 | tumor | 0,27 | 0,83 | 0,64 | 4,61 | 13,47 | 0,25 | 3,01 |
| HMN_PC_140 | tumor | 1,08 | 0,06 | 0,85 | 0,77 | 1,14 | 4,82 | 2,67 |
| HMN_PC_141 | tumor | 0,76 | 2,16 | 1,35 | 1,66 | 1,13 | 1,20 | 3,59 |
| HMN_PC_142 | tumor | 0,80 | 0,88 | 9,09 | 1,75 | 1,94 | 1,87 | 3,16 |
| HMN_PC_143 | tumor | 0,73 | 0,79 | 0,49 | 0,25 | 17,91 | 3,96 | 1,58 |
| HMN_PC_145 | tumor | 0,30 | 0,40 | 4,03 | 0,34 | 0,27 | 0,52 | 2,04 |
| HMN_PC_146 | tumor | 0,52 | 1,08 | 0,79 | 2,40 | 3,11 | 0,45 | 3,34 |
| HMN_PC_147 | tumor | 0,08 | 0,05 | 2,62 | 0,54 | 4,43 | 0,21 | 0,29 |
| HMN_PC_148 | tumor | 0,30 | 0,49 | 0,20 | 0,99 | 8,53 | 1,84 | 3,89 |
| HMN_PC_149(-) | normal | 0,01 | 0,04 | 0,30 | 0,17 | 0,14 | 0,22 | 0,16 |
| HMN_PC_152 | tumor | 0,77 | 0,83 | 1,15 | 1,10 | 36,87 | 1,32 | 1,48 |
| HMN_PC_153 | tumor | 0,14 | 0,03 | 0,49 | 1,34 | 2,18 | 0,64 | 0,05 |
| HMN_PC_156 | tumor | 0,99 | 0,56 | 4,80 | 5,41 | 12,14 | 1,73 | 1,17 |
| HMN_PC_157 | tumor | 0,20 | 0,87 | 0,25 | 0,38 | 28,07 | 2,49 | 1,39 |
| HMN_PC_158 | tumor | 0,63 | 0,18 | 0,99 | 2,51 | 24,89 | 2,65 | 0,74 |
| HMN_PC_160 | tumor | 0,22 | 0,11 | 1,64 | 0,66 | 13,12 | 0,16 | 0,71 |
| HMN_PC_161 | tumor | 0,15 | 0,19 | 0,92 | 0,18 | 7,59 | 1,74 | 0,64 |
| HMN_PC_162 | tumor | 0,38 | 0,19 | 4,70 | 0,06 | 11,46 | 0,16 | 4,74 |
| HMN_PC_164 | tumor | 0,13 | 0,08 | 2,55 | 1,98 | 8,07 | 0,42 | 1,80 |
| HMN_PC_166 | tumor | 0,22 | 1,19 | 0,38 | 0,08 | 35,96 | 0,66 | 3,58 |
| HMN_PC_167 | tumor | 0,24 | 0,56 | 4,35 | 0,36 | 31,87 | 0,61 | 1,10 |
| HMN_PC_168 | tumor | 1,21 | 1,11 | 1,33 | 3,69 | 15,09 | 3,02 | 3,15 |
| HMN_PC_169 | tumor | 0,30 | 1,77 | 2,59 | 0,90 | 0,95 | 0,65 | 2,15 |
| HMN_PC_170 | tumor | 0,58 | 1,35 | 0,31 | 0,79 | 11,70 | 1,15 | 4,44 |
| HMN_PC_171 | tumor | 0,15 | 0,14 | 3,02 | 0,21 | 0,25 | 0,06 | 1,55 |
| HMN_PC_172 | tumor | 0,18 | 0,96 | 0,10 | 0,20 | 53,80 | 0,24 | 1,51 |
| HMN_PC_173 | tumor | 0,28 | 0,44 | 3,43 | 0,52 | 23,91 | 0,34 | 2,22 |
| HMN_PC_174 | tumor | 0,42 | 0,46 | 1,59 | 3,39 | 26,36 | 1,15 | 1,63 |
| HMN_PC_209 | tumor | 0,22 | 0,65 | 0,26 | 2,18 | 8,65 | 3,69 | 0,76 |
| HMN_PC_217 | tumor | 0,19 | 0,10 | 0,40 | 0,27 | 21,73 | 0,33 | 1,01 |
| HMN_PC_218 | tumor | 0,19 | 0,09 | 0,68 | 0,70 | 12,26 | 1,48 | 2,87 |

| ID_sample | Tissue | ctg_123090 | ctg_25348 | ctg_73782 | ctg_104447 | PCA3 | ctg_105149 | ctg_111158 |
|---|---|---|---|---|---|---|---|---|
| HMN_PC_219 | tumor | 0,14 | 0,53 | 0,16 | 0,01 | 27,06 | 0,40 | 2,25 |
| HMN_PC_220 | tumor | 0,11 | 0,46 | 13,81 | 0,28 | 2,09 | 2,16 | 1,43 |
| HMN_PC_221 | tumor | 0,14 | 1,19 | 0,36 | 3,08 | 11,10 | 0,87 | 1,27 |
| HMN_PC_222 | tumor | 0,18 | 0,76 | 0,27 | 0,26 | 29,31 | 0,88 | 1,37 |
| HMN_PC_223 | tumor | 0,23 | 0,15 | 7,43 | 0,62 | 1,90 | 0,37 | 1,40 |
| HMN_PC_224 | tumor | 0,46 | 0,37 | 4,47 | 0,16 | 13,24 | 0,49 | 2,38 |
| HMN_PC_225 | tumor | 0,74 | 1,32 | 1,35 | 1,86 | 12,64 | 2,72 | 3,71 |
| HMN_PC_226 | tumor | 0,36 | 0,41 | 0,05 | 0,58 | 53,74 | 1,17 | 0,96 |
| HMN_PC_227 | tumor | 0,14 | 0,66 | 0,23 | 0,70 | 3,59 | 0,86 | 4,03 |
| HMN_PC_228 | tumor | 0,36 | 0,60 | 4,37 | 0,69 | 35,65 | 0,88 | 1,35 |
| HMN_PC_229 | tumor | 0,81 | 0,50 | 12,17 | 0,87 | 3,17 | 0,83 | 1,21 |
| HMN_PC_232 | tumor | 0,20 | 0,57 | 0,17 | 2,43 | 19,95 | 1,44 | 2,64 |
| HMN_PC_233 | tumor | 0,24 | 0,17 | 7,83 | 1,26 | 13,25 | 0,59 | 0,53 |
| HMN_PC_234 | tumor | 0,42 | 0,19 | 2,04 | 0,39 | 26,57 | 0,23 | 2,66 |
| HMN_PC_235 | tumor | 0,54 | 0,36 | 1,65 | 2,41 | 12,14 | 1,85 | 1,85 |
| HMN_PC_236 | tumor | 0,21 | 1,88 | 2,93 | 1,05 | 14,01 | 0,49 | 2,25 |
| HMN_PC_239 | tumor | 0,22 | 0,42 | 1,69 | 2,90 | 15,70 | 2,49 | 0,96 |
| HMN_PC_241 | tumor | 0,48 | 0,85 | 0,27 | 2,33 | 4,22 | 5,76 | 1,70 |
| HMN_PC_242 | tumor | 0,21 | 1,30 | 0,17 | 0,44 | 21,87 | 0,95 | 4,35 |
| HMN_PC_243 | tumor | 0,07 | 0,22 | 1,76 | 0,27 | 19,70 | 0,14 | 0,37 |
| HMN_PC_244 | tumor | 0,82 | 0,69 | 0,18 | 0,17 | 2,36 | 2,15 | 2,78 |
| HMN_PC_245 | tumor | 0,26 | 0,09 | 2,29 | 0,46 | 12,65 | 0,33 | 2,16 |
| HMN_PC_246 | tumor | 0,90 | 0,36 | 5,19 | 2,05 | 4,82 | 1,54 | 2,47 |
| HMN_PC_247 | tumor | 0,36 | 0,34 | 0,47 | 2,33 | 14,05 | 2,57 | 0,72 |

| ID_sample | ctg_37852 | ctg_23999 | ctg_111348 | ctg_117356 | ctg_119680 | ctg_17297 | ctg_2815 | ctg_28650 |
|---|---|---|---|---|---|---|---|---|
| HMN_PC_001 | 4,24 | 0,83 | 2,52 | 8,40 | 0,36 | 10,61 | 2,29 | 1,76 |
| HMN_PC_010 | 3,79 | 2,18 | 1,06 | 7,17 | 0,20 | 5,62 | 0,81 | 1,01 |
| HMN_PC_013 | 6,55 | 0,67 | 2,04 | 4,85 | 0,18 | 0,72 | 0,73 | 0,68 |
| HMN_PC_014 | 1,63 | 0,78 | 3,06 | 2,46 | 0,08 | 8,35 | 0,34 | 0,12 |
| HMN_PC_016 | 0,78 | 1,63 | 0,02 | 1,01 | 0,20 | 15,11 | 1,68 | 0,47 |
| HMN_PC_019 | 6,19 | 1,40 | 5,35 | 3,11 | 0,28 | 1,65 | 2,39 | 0,61 |
| HMN_PC_020 | 22,79 | 0,42 | 3,20 | 7,69 | 0,20 | 31,29 | 1,14 | 1,85 |
| HMN_PC_021 | 2,55 | 0,90 | 0,45 | 3,45 | 0,31 | 15,32 | 0,91 | 0,44 |
| HMN_PC_022 | 0,13 | 0,48 | 0,14 | 2,63 | 0,44 | 16,04 | 1,91 | 0,34 |
| HMN_PC_023 | 3,51 | 1,79 | 1,57 | 1,80 | 0,38 | 7,70 | 1,63 | 0,67 |
| HMN_PC_024 | 18,52 | 2,59 | 1,55 | 4,04 | 0,32 | 9,58 | 1,46 | 0,31 |
| HMN_PC_026 | 0,74 | 0,69 | 1,50 | 2,26 | 0,37 | 26,17 | 2,43 | 1,03 |
| HMN_PC_027 | 3,62 | 0,51 | 2,51 | 4,25 | 0,23 | 1,84 | 1,12 | 0,03 |
| HMN_PC_028 | 8,45 | 1,00 | 3,42 | 2,60 | 0,03 | 5,83 | 1,23 | 0,47 |
| HMN_PC_030 | 11,94 | 0,76 | 3,92 | 2,67 | 0,11 | 4,86 | 0,99 | 1,87 |
| HMN_PC_031 | 7,08 | 1,24 | 0,31 | 6,29 | 0,17 | 1,62 | 1,18 | 0,58 |
| HMN_PC_032 | 0,47 | 0,08 | 3,79 | 0,63 | 0,07 | 10,75 | 0,53 | 2,19 |
| HMN_PC_033 | 3,28 | 1,40 | 0,07 | 7,95 | 0,26 | 9,32 | 1,15 | 0,72 |
| HMN_PC_036 | 4,51 | 0,25 | 1,08 | 0,69 | 0,19 | 0,53 | 0,36 | 0,51 |
| HMN_PC_037 | 40,50 | 0,04 | 0,02 | 1,52 | 0,05 | 4,14 | 0,72 | 0,35 |
| HMN_PC_038 | 4,38 | 0,48 | 8,35 | 2,12 | 0,14 | 0,92 | 0,94 | 1,01 |
| HMN_PC_040 | 1,39 | 0,01 | 0,01 | 0,98 | 0,14 | 2,54 | 0,42 | 0,54 |
| HMN_PC_041 | 2,04 | 0,62 | 4,20 | 5,73 | 0,27 | 8,12 | 0,47 | 0,83 |
| HMN_PC_043 | 6,11 | 10,30 | 0,34 | 6,62 | 0,41 | 0,63 | 3,75 | 0,73 |
| HMN_PC_044 | 0,99 | 1,16 | 1,27 | 3,48 | 0,11 | 7,68 | 1,50 | 1,07 |
| HMN_PC_045 | 1,52 | 0,70 | 2,73 | 5,51 | 0,09 | 8,21 | 0,27 | 0,77 |
| HMN_PC_047 | 5,80 | 0,23 | 2,53 | 0,83 | 0,08 | 5,13 | 0,91 | 0,55 |
| HMN_PC_049 | 0,93 | 0,26 | 4,26 | 2,92 | 0,19 | 5,89 | 0,54 | 1,26 |
| HMN_PC_050 | 3,65 | 1,27 | 4,11 | 0,88 | 0,10 | 6,67 | 1,90 | 0,82 |
| HMN_PC_051 | 9,76 | 5,18 | 0,84 | 8,36 | 0,15 | 8,42 | 2,09 | 0,72 |
| HMN_PC_052 | 18,23 | 0,26 | 1,72 | 3,78 | 0,20 | 0,33 | 2,83 | 0,97 |
| HMN_PC_056 | 16,58 | 2,97 | 5,49 | 1,96 | 0,18 | 1,49 | 1,65 | 1,55 |
| HMN_PC_057 | 0,01 | 1,30 | 0,23 | 3,70 | 0,29 | 23,64 | 0,91 | 1,37 |
| HMN_PC_058(-) | 0,23 | 0,02 | 0,02 | 0,37 | 0,16 | 0,36 | 0,28 | 0,02 |
| HMN_PC_059(-) | 0,16 | 0,18 | 0,03 | 0,04 | 0,04 | 0,15 | 0,39 | 0,03 |
| HMN_PC_060(-) | 0,13 | 0,02 | 0,02 | 0,07 | 0,02 | 0,22 | 0,93 | 0,15 |
| HMN_PC_063 | 0,25 | 0,22 | 4,05 | 1,83 | 0,14 | 0,56 | 0,89 | 1,29 |
| HMN_PC_066 | 18,15 | 0,30 | 4,95 | 3,21 | 0,12 | 0,25 | 1,10 | 0,79 |
| HMN_PC_067 | 2,24 | 1,01 | 0,01 | 6,50 | 0,16 | 4,60 | 1,52 | 0,51 |
| HMN_PC_068 | 2,25 | 0,14 | 3,63 | 2,42 | 0,23 | 5,09 | 2,33 | 3,37 |
| HMN_PC_070 | 1,51 | 0,55 | 2,37 | 1,70 | 0,12 | 5,16 | 1,81 | 1,30 |
| HMN_PC_071 | 2,81 | 0,53 | 0,40 | 4,86 | 0,21 | 24,41 | 3,06 | 1,43 |
| HMN_PC_072 | 0,01 | 0,58 | 3,10 | 5,15 | 0,04 | 1,88 | 0,68 | 0,19 |
| HMN_PC_075 | 4,36 | 0,24 | 5,23 | 1,77 | 0,07 | 11,28 | 0,77 | 1,01 |
| HMN_PC_076 | 17,47 | 8,49 | 1,12 | 3,45 | 0,08 | 12,90 | 1,76 | 2,66 |
| HMN_PC_077.1 | 2,68 | 1,36 | 2,45 | 5,09 | 0,19 | 3,35 | 2,51 | 1,12 |
| HMN_PC_077.2 | 0,25 | 0,10 | 0,25 | 0,42 | 0,16 | 0,33 | 0,74 | 0,54 |
| HMN_PC_078(-) | 0,41 | 0,04 | 0,08 | 0,34 | 0,23 | 0,32 | 0,86 | 0,04 |
| HMN_PC_081 | 26,45 | 4,80 | 0,01 | 7,68 | 0,31 | 14,35 | 1,66 | 0,61 |
| HMN_PC_082 | 7,07 | 5,75 | 0,58 | 1,85 | 0,51 | 6,93 | 4,37 | 2,07 |
| HMN_PC_084 | 3,04 | 0,08 | 0,18 | 2,08 | 0,11 | 3,59 | 1,14 | 0,24 |
| HMN_PC_085 | 0,29 | 0,20 | 2,08 | 2,51 | 0,10 | 5,05 | 0,82 | 1,68 |
| HMN_PC_086(-) | 0,12 | 0,12 | 0,04 | 0,20 | 0,22 | 0,38 | 1,09 | 0,16 |
| HMN_PC_087 | 6,53 | 4,37 | 0,49 | 1,76 | 0,22 | 6,65 | 1,90 | 0,67 |
| HMN_PC_088(-) | 0,41 | 0,15 | 0,02 | 0,54 | 0,18 | 0,32 | 0,52 | 0,17 |
| HMN_PC_089 | 23,97 | 5,51 | 3,54 | 7,69 | 0,38 | 9,53 | 1,52 | 0,56 |
| HMN_PC_092 | 3,61 | 0,23 | 5,61 | 3,59 | 0,29 | 8,18 | 0,91 | 0,98 |
| HMN_PC_093(-) | 1,52 | 0,38 | 0,01 | 0,81 | 0,01 | 2,18 | 0,54 | 0,07 |
| HMN_PC_096 | 1,01 | 0,25 | 0,61 | 0,35 | 0,07 | 0,46 | 0,77 | 0,30 |

| ID_sample | ctg_37852 | ctg_23999 | ctg_111348 | ctg_117356 | ctg_119680 | ctg_17297 | ctg_2815 | ctg_28650 |
|---|---|---|---|---|---|---|---|---|
| HMN_PC_097 | 15,69 | 0,06 | 3,43 | 5,41 | 0,13 | 5,96 | 1,13 | 0,88 |
| HMN_PC_099 | 0,91 | 0,38 | 0,78 | 1,52 | 0,08 | 0,49 | 1,18 | 0,65 |
| HMN_PC_100 | 4,01 | 0,29 | 0,82 | 1,23 | 0,08 | 8,08 | 1,10 | 0,66 |
| HMN_PC_101 | 1,44 | 0,84 | 1,36 | 1,38 | 0,15 | 2,55 | 0,51 | 0,19 |
| HMN_PC_102 | 6,52 | 1,58 | 1,14 | 7,55 | 0,19 | 4,46 | 1,32 | 0,54 |
| HMN_PC_103 | 12,57 | 4,15 | 8,52 | 6,08 | 0,13 | 1,48 | 2,71 | 2,87 |
| HMN_PC_104 | 21,66 | 7,61 | 0,79 | 2,05 | 0,09 | 0,73 | 2,04 | 0,75 |
| HMN_PC_105 | 0,94 | 0,59 | 0,82 | 1,81 | 0,01 | 2,94 | 2,43 | 1,59 |
| HMN_PC_106 | 6,57 | 1,48 | 2,20 | 4,39 | 0,09 | 2,09 | 3,07 | 1,10 |
| HMN_PC_110 | 21,83 | 1,04 | 1,36 | 4,52 | 0,23 | 11,79 | 2,17 | 1,98 |
| HMN_PC_111 | 1,28 | 0,26 | 2,21 | 0,49 | 0,07 | 10,08 | 0,74 | 0,92 |
| HMN_PC_113 | 0,64 | 3,14 | 0,83 | 2,63 | 0,24 | 5,76 | 3,13 | 1,72 |
| HMN_PC_114 | 56,36 | 0,36 | 3,48 | 3,32 | 0,27 | 2,10 | 0,79 | 0,56 |
| HMN_PC_115 | 2,81 | 0,58 | 0,30 | 1,83 | 0,17 | 0,94 | 0,69 | 0,49 |
| HMN_PC_117 | 12,31 | 1,79 | 1,72 | 6,47 | 0,30 | 14,69 | 1,47 | 0,75 |
| HMN_PC_118 | 3,95 | 0,23 | 0,01 | 0,33 | 0,08 | 2,71 | 2,05 | 0,32 |
| HMN_PC_119 | 13,17 | 1,11 | 9,47 | 1,91 | 0,74 | 37,18 | 2,67 | 1,75 |
| HMN_PC_121 | 23,70 | 0,25 | 4,46 | 2,69 | 0,12 | 4,47 | 2,41 | 2,57 |
| HMN_PC_124 | 0,12 | 3,29 | 0,56 | 15,28 | 0,35 | 31,70 | 1,23 | 2,40 |
| HMN_PC_125(-) | 0,02 | 0,02 | 0,05 | 0,17 | 0,04 | 0,15 | 0,48 | 0,12 |
| HMN_PC_126 | 6,03 | 0,61 | 4,13 | 7,30 | 0,41 | 14,44 | 0,97 | 1,94 |
| HMN_PC_127 | 0,45 | 2,13 | 3,08 | 0,91 | 0,20 | 11,63 | 1,22 | 2,07 |
| HMN_PC_128 | 0,72 | 1,80 | 1,91 | 2,93 | 0,25 | 6,82 | 2,65 | 1,04 |
| HMN_PC_129 | 1,03 | 2,91 | 1,76 | 4,67 | 0,26 | 5,86 | 2,28 | 0,74 |
| HMN_PC_130 | 7,04 | 22,27 | 6,12 | 27,73 | 0,40 | 11,90 | 4,07 | 2,29 |
| HMN_PC_131 | 24,80 | 3,18 | 4,00 | 5,12 | 0,58 | 18,61 | 3,79 | 1,78 |
| HMN_PC_134 | 1,91 | 2,27 | 0,05 | 7,07 | 0,11 | 0,76 | 4,12 | 1,53 |
| HMN_PC_135 | 0,37 | 0,84 | 4,18 | 1,14 | 0,21 | 4,16 | 3,26 | 1,79 |
| HMN_PC_137 | 0,01 | 0,10 | 0,54 | 0,63 | 0,15 | 10,49 | 3,07 | 1,61 |
| HMN_PC_138 | 21,29 | 5,69 | 0,01 | 2,83 | 0,02 | 0,36 | 1,96 | 0,54 |
| HMN_PC_139 | 6,51 | 8,64 | 0,02 | 0,63 | 0,19 | 5,45 | 1,92 | 1,55 |
| HMN_PC_140 | 6,62 | 2,02 | 3,74 | 2,92 | 0,26 | 20,98 | 3,14 | 1,59 |
| HMN_PC_141 | 0,42 | 1,27 | 5,77 | 9,10 | 0,21 | 23,11 | 4,49 | 2,03 |
| HMN_PC_142 | 4,42 | 3,66 | 1,53 | 2,92 | 0,27 | 7,91 | 3,10 | 1,66 |
| HMN_PC_143 | 1,03 | 0,97 | 0,47 | 2,29 | 0,12 | 0,23 | 2,35 | 1,42 |
| HMN_PC_145 | 0,90 | 0,54 | 1,59 | 1,50 | 0,10 | 7,73 | 1,69 | 0,94 |
| HMN_PC_146 | 2,27 | 0,91 | 2,34 | 0,74 | 0,24 | 20,26 | 1,44 | 1,97 |
| HMN_PC_147 | 2,46 | 0,07 | 0,32 | 2,76 | 0,11 | 2,58 | 0,28 | 0,27 |
| HMN_PC_148 | 0,14 | 0,32 | 4,77 | 0,14 | 0,06 | 3,87 | 1,76 | 2,65 |
| HMN_PC_149(-) | 0,01 | 0,10 | 0,01 | 0,16 | 0,10 | 0,16 | 1,09 | 0,19 |
| HMN_PC_152 | 17,25 | 2,79 | 1,30 | 11,24 | 0,13 | 3,83 | 1,34 | 0,88 |
| HMN_PC_153 | 0,86 | 0,12 | 0,02 | 0,32 | 0,09 | 0,40 | 0,80 | 0,06 |
| HMN_PC_156 | 1,14 | 2,45 | 3,75 | 7,17 | 0,28 | 7,86 | 2,51 | 0,68 |
| HMN_PC_157 | 11,61 | 2,38 | 1,97 | 8,34 | 0,06 | 3,88 | 2,45 | 0,72 |
| HMN_PC_158 | 6,07 | 0,65 | 4,31 | 7,03 | 0,18 | 6,10 | 1,28 | 0,52 |
| HMN_PC_160 | 5,76 | 0,26 | 1,50 | 1,34 | 0,10 | 2,17 | 0,42 | 0,53 |
| HMN_PC_161 | 0,12 | 0,43 | 0,10 | 1,40 | 0,16 | 1,44 | 0,75 | 0,39 |
| HMN_PC_162 | 0,13 | 0,11 | 3,89 | 0,36 | 0,05 | 0,95 | 0,61 | 1,46 |
| HMN_PC_164 | 11,81 | 0,15 | 2,23 | 1,90 | 0,11 | 2,36 | 0,63 | 0,85 |
| HMN_PC_166 | 3,98 | 0,86 | 4,69 | 3,65 | 0,11 | 23,27 | 0,79 | 1,07 |
| HMN_PC_167 | 7,38 | 0,50 | 2,21 | 2,55 | 0,18 | 4,13 | 0,62 | 0,59 |
| HMN_PC_168 | 0,62 | 8,42 | 1,11 | 6,69 | 0,15 | 3,49 | 2,55 | 1,71 |
| HMN_PC_169 | 0,01 | 0,57 | 2,39 | 2,72 | 0,06 | 9,16 | 1,72 | 1,27 |
| HMN_PC_170 | 0,01 | 0,81 | 1,35 | 1,88 | 0,11 | 7,84 | 4,62 | 2,52 |
| HMN_PC_171 | 2,13 | 0,17 | 2,48 | 3,02 | 0,10 | 11,18 | 0,30 | 0,75 |
| HMN_PC_172 | 2,07 | 1,58 | 0,03 | 3,60 | 0,04 | 0,15 | 0,65 | 0,94 |
| HMN_PC_173 | 1,82 | 0,89 | 2,30 | 4,83 | 0,12 | 13,89 | 0,90 | 1,03 |
| HMN_PC_174 | 0,24 | 1,27 | 1,84 | 3,26 | 0,22 | 0,74 | 1,69 | 0,82 |
| HMN_PC_209 | 6,95 | 1,31 | 2,92 | 3,63 | 0,09 | 2,02 | 1,27 | 0,59 |
| HMN_PC_217 | 0,52 | 0,08 | 2,12 | 2,06 | 0,07 | 3,31 | 0,56 | 0,74 |
| HMN_PC_218 | 3,80 | 1,17 | 0,23 | 10,27 | 0,07 | 22,47 | 0,98 | 1,45 |

### PAIR_NanoString

| ID_sample | ctg_37852 | ctg_23999 | ctg_111348 | ctg_117356 | ctg_119680 | ctg_17297 | ctg_2815 | ctg_28650 |
|---|---|---|---|---|---|---|---|---|
| HMN_PC_219 | 2,76 | 0,03 | 0,72 | 3,61 | 0,05 | 1,58 | 0,76 | 1,25 |
| HMN_PC_220 | 0,18 | 0,24 | 3,03 | 0,30 | 0,03 | 1,34 | 1,25 | 0,59 |
| HMN_PC_221 | 0,72 | 3,03 | 0,03 | 8,06 | 0,12 | 0,43 | 1,31 | 0,64 |
| HMN_PC_222 | 5,82 | 1,02 | 0,33 | 6,57 | 0,24 | 8,94 | 1,89 | 0,70 |
| HMN_PC_223 | 0,11 | 0,13 | 4,10 | 1,30 | 0,13 | 2,23 | 0,83 | 0,52 |
| HMN_PC_224 | 1,10 | 0,26 | 1,46 | 1,08 | 0,10 | 5,34 | 1,89 | 1,44 |
| HMN_PC_225 | 1,08 | 11,65 | 1,53 | 12,28 | 0,29 | 5,67 | 3,45 | 1,43 |
| HMN_PC_226 | 6,27 | 1,70 | 0,19 | 3,09 | 0,04 | 0,90 | 0,67 | 0,57 |
| HMN_PC_227 | 1,84 | 3,43 | 3,76 | 8,40 | 0,09 | 7,79 | 1,02 | 1,54 |
| HMN_PC_228 | 9,24 | 0,85 | 0,10 | 4,90 | 0,22 | 8,69 | 1,19 | 0,77 |
| HMN_PC_229 | 0,13 | 0,84 | 1,42 | 2,66 | 0,19 | 6,43 | 1,96 | 0,70 |
| HMN_PC_232 | 3,19 | 4,73 | 3,89 | 10,31 | 0,03 | 9,40 | 0,79 | 1,64 |
| HMN_PC_233 | 2,18 | 0,49 | 1,12 | 5,10 | 0,19 | 2,85 | 0,50 | 0,44 |
| HMN_PC_234 | 0,26 | 0,18 | 1,26 | 2,95 | 0,19 | 2,27 | 1,55 | 1,04 |
| HMN_PC_235 | 4,69 | 1,05 | 2,12 | 3,90 | 0,10 | 1,54 | 1,28 | 1,00 |
| HMN_PC_236 | 0,44 | 0,34 | 2,14 | 2,00 | 0,21 | 0,93 | 1,24 | 0,85 |
| HMN_PC_239 | 7,44 | 2,47 | 0,03 | 2,16 | 0,25 | 0,40 | 2,66 | 0,58 |
| HMN_PC_241 | 0,51 | 4,08 | 0,92 | 10,39 | 0,10 | 0,27 | 1,94 | 0,67 |
| HMN_PC_242 | 9,68 | 2,08 | 2,07 | 5,39 | 0,18 | 14,39 | 0,84 | 1,31 |
| HMN_PC_243 | 1,06 | 0,28 | 2,75 | 1,74 | 0,03 | 1,85 | 0,32 | 0,27 |
| HMN_PC_244 | 1,02 | 0,95 | 6,20 | 6,34 | 0,11 | 13,19 | 6,10 | 1,13 |
| HMN_PC_245 | 2,11 | 0,49 | 3,39 | 2,66 | 0,16 | 4,95 | 1,03 | 0,89 |
| HMN_PC_246 | 3,43 | 0,68 | 1,25 | 11,89 | 0,18 | 10,05 | 2,06 | 0,84 |
| HMN_PC_247 | 0,31 | 3,75 | 0,04 | 1,57 | 0,29 | 4,57 | 2,01 | 0,54 |

| ID_sample | ctg_29077 | ctg_36195 | ctg_44030 | ctg_512 | ctg_57223 | ctg_61472 | ctg_61528 | ctg_63866 | ctg_9446 |
|---|---|---|---|---|---|---|---|---|---|
| HMN_PC_001 | 0,27 | 0,25 | 7,59 | 1,39 | 0,41 | 3,64 | 3,15 | 0,29 | 5,83 |
| HMN_PC_010 | 1,48 | 0,13 | 1,77 | 1,26 | 0,22 | 3,69 | 0,30 | 0,52 | 0,78 |
| HMN_PC_013 | 0,33 | 0,17 | 1,35 | 0,68 | 0,57 | 2,13 | 0,61 | 0,31 | 0,84 |
| HMN_PC_014 | 1,01 | 0,11 | 5,85 | 1,17 | 1,22 | 1,68 | 1,40 | 0,42 | 1,66 |
| HMN_PC_016 | 0,20 | 0,05 | 3,09 | 0,92 | 0,49 | 0,34 | 0,82 | 0,02 | 1,34 |
| HMN_PC_019 | 3,67 | 0,61 | 5,80 | 1,76 | 1,29 | 1,70 | 1,65 | 3,10 | 0,92 |
| HMN_PC_020 | 4,91 | 0,28 | 5,72 | 1,72 | 2,06 | 6,31 | 1,53 | 5,69 | 1,57 |
| HMN_PC_021 | 0,59 | 0,52 | 1,38 | 1,14 | 0,49 | 2,85 | 0,84 | 0,19 | 0,89 |
| HMN_PC_022 | 0,19 | 0,65 | 0,59 | 0,34 | 0,24 | 2,24 | 0,19 | 0,43 | 0,55 |
| HMN_PC_023 | 2,59 | 0,49 | 2,44 | 0,38 | 0,62 | 2,99 | 0,73 | 1,81 | 0,98 |
| HMN_PC_024 | 0,27 | 0,30 | 5,33 | 0,99 | 2,11 | 2,06 | 0,81 | 0,28 | 1,55 |
| HMN_PC_026 | 0,05 | 0,08 | 1,15 | 0,56 | 1,38 | 1,27 | 0,44 | 0,10 | 1,27 |
| HMN_PC_027 | 4,20 | 0,14 | 4,81 | 1,20 | 3,19 | 2,17 | 2,31 | 1,73 | 2,06 |
| HMN_PC_028 | 2,55 | 0,13 | 3,50 | 0,76 | 1,26 | 1,72 | 1,73 | 4,78 | 1,19 |
| HMN_PC_030 | 2,31 | 0,07 | 1,08 | 0,39 | 2,72 | 1,58 | 0,26 | 1,33 | 1,39 |
| HMN_PC_031 | 0,41 | 0,43 | 4,03 | 2,86 | 0,22 | 3,29 | 0,54 | 0,11 | 0,70 |
| HMN_PC_032 | 0,92 | 0,18 | 4,74 | 0,95 | 0,90 | 0,41 | 1,30 | 1,46 | 0,73 |
| HMN_PC_033 | 0,50 | 0,15 | 2,15 | 1,05 | 0,53 | 4,27 | 0,64 | 0,45 | 0,49 |
| HMN_PC_036 | 0,79 | 0,03 | 0,75 | 1,25 | 0,27 | 0,26 | 0,22 | 0,03 | 0,42 |
| HMN_PC_037 | 2,11 | 0,02 | 1,29 | 1,07 | 0,63 | 0,46 | 0,08 | 0,34 | 0,83 |
| HMN_PC_038 | 0,95 | 0,01 | 3,53 | 0,98 | 1,38 | 1,09 | 0,98 | 1,34 | 0,67 |
| HMN_PC_040 | 0,27 | 0,06 | 2,43 | 0,85 | 0,64 | 0,38 | 0,75 | 0,07 | 0,66 |
| HMN_PC_041 | 0,19 | 0,04 | 4,46 | 0,97 | 1,03 | 2,60 | 0,82 | 0,36 | 1,94 |
| HMN_PC_043 | 0,65 | 0,31 | 12,60 | 2,06 | 0,87 | 3,53 | 4,39 | 0,53 | 1,85 |
| HMN_PC_044 | 0,22 | 0,30 | 1,98 | 0,77 | 0,30 | 2,69 | 0,79 | 0,67 | 1,26 |
| HMN_PC_045 | 1,99 | 0,05 | 1,20 | 1,33 | 2,97 | 2,44 | 0,43 | 0,48 | 0,30 |
| HMN_PC_047 | 3,61 | 0,59 | 1,76 | 1,05 | 0,73 | 1,06 | 0,37 | 2,20 | 1,14 |
| HMN_PC_049 | 3,83 | 0,33 | 6,11 | 1,38 | 0,77 | 1,13 | 1,45 | 1,63 | 3,51 |
| HMN_PC_050 | 2,95 | 0,20 | 6,07 | 0,68 | 2,28 | 0,38 | 2,47 | 0,81 | 4,26 |
| HMN_PC_051 | 2,39 | 0,27 | 3,36 | 1,35 | 1,86 | 4,62 | 1,25 | 2,02 | 2,12 |
| HMN_PC_052 | 0,30 | 0,14 | 7,17 | 1,75 | 3,88 | 1,57 | 2,98 | 0,10 | 2,35 |
| HMN_PC_056 | 2,83 | 0,09 | 3,06 | 1,72 | 0,79 | 1,46 | 1,78 | 1,92 | 1,15 |
| HMN_PC_057 | 0,48 | 0,20 | 1,87 | 0,38 | 0,09 | 2,22 | 0,65 | 0,45 | 0,52 |
| HMN_PC_058(-) | 0,08 | 0,12 | 0,24 | 0,49 | 0,34 | 0,02 | 0,10 | 0,02 | 0,45 |
| HMN_PC_059(-) | 0,08 | 0,05 | 0,45 | 0,06 | 0,03 | 0,03 | 0,08 | 0,03 | 0,50 |
| HMN_PC_060(-) | 0,05 | 0,08 | 0,16 | 0,19 | 0,14 | 0,02 | 0,02 | 0,02 | 0,48 |
| HMN_PC_063 | 2,70 | 0,02 | 5,55 | 2,05 | 2,08 | 0,56 | 1,66 | 1,81 | 2,35 |
| HMN_PC_066 | 3,38 | 0,14 | 2,93 | 1,36 | 0,58 | 1,15 | 0,68 | 1,00 | 0,98 |
| HMN_PC_067 | 0,20 | 0,14 | 0,18 | 1,05 | 0,38 | 3,31 | 0,11 | 0,10 | 1,82 |
| HMN_PC_068 | 5,37 | 0,16 | 9,10 | 1,09 | 4,68 | 1,64 | 5,62 | 8,57 | 2,80 |
| HMN_PC_070 | 0,70 | 0,25 | 1,93 | 0,35 | 1,01 | 0,54 | 0,67 | 0,57 | 3,54 |
| HMN_PC_071 | 0,70 | 0,05 | 4,63 | 1,47 | 2,53 | 2,34 | 1,46 | 0,28 | 1,63 |
| HMN_PC_072 | 0,41 | 0,67 | 5,11 | 0,83 | 2,02 | 2,19 | 1,08 | 0,72 | 2,05 |
| HMN_PC_075 | 0,55 | 0,23 | 2,87 | 0,96 | 0,33 | 1,14 | 0,59 | 0,83 | 0,85 |
| HMN_PC_076 | 0,55 | 0,24 | 0,66 | 0,33 | 1,93 | 1,79 | 0,24 | 0,40 | 1,40 |
| HMN_PC_077.1 | 3,43 | 0,10 | 6,71 | 0,76 | 6,47 | 2,38 | 3,54 | 2,53 | 3,87 |
| HMN_PC_077.2 | 0,22 | 0,39 | 3,82 | 0,93 | 1,42 | 0,27 | 0,76 | 0,42 | 0,78 |
| HMN_PC_078(-) | 0,04 | 0,21 | 0,32 | 0,04 | 0,46 | 0,04 | 0,04 | 0,04 | 0,38 |
| HMN_PC_081 | 0,65 | 0,06 | 2,24 | 1,30 | 0,84 | 4,94 | 0,91 | 0,34 | 1,52 |
| HMN_PC_082 | 0,96 | 0,02 | 4,94 | 1,31 | 0,59 | 0,83 | 3,14 | 0,74 | 2,51 |
| HMN_PC_084 | 0,73 | 0,23 | 1,35 | 0,04 | 0,71 | 0,78 | 0,23 | 0,04 | 1,45 |
| HMN_PC_085 | 1,98 | 0,42 | 0,70 | 0,38 | 0,62 | 0,91 | 0,20 | 0,07 | 1,71 |
| HMN_PC_086(-) | 0,74 | 0,26 | 0,48 | 0,07 | 0,21 | 0,04 | 0,19 | 0,35 | 0,50 |
| HMN_PC_087 | 0,13 | 0,06 | 2,62 | 1,01 | 0,78 | 0,98 | 0,99 | 0,22 | 1,60 |
| HMN_PC_088(-) | 0,34 | 0,11 | 0,66 | 0,29 | 0,30 | 0,57 | 0,15 | 0,12 | 0,68 |
| HMN_PC_089 | 3,53 | 0,69 | 17,82 | 2,16 | 7,36 | 4,66 | 4,05 | 1,53 | 2,28 |
| HMN_PC_092 | 3,33 | 0,16 | 4,34 | 1,97 | 7,90 | 2,26 | 0,81 | 8,36 | 1,56 |
| HMN_PC_093(-) | 0,01 | 0,08 | 0,79 | 0,23 | 0,09 | 0,20 | 0,11 | 0,01 | 0,39 |
| HMN_PC_096 | 1,09 | 0,42 | 5,06 | 0,51 | 0,50 | 0,16 | 1,50 | 0,54 | 1,03 |

| ID_sample | ctg_29077 | ctg_36195 | ctg_44030 | ctg_512 | ctg_57223 | ctg_61472 | ctg_61528 | ctg_63866 | ctg_9446 |
|---|---|---|---|---|---|---|---|---|---|
| HMN_PC_097 | 3,95 | 0,04 | 5,38 | 2,95 | 1,26 | 3,16 | 0,71 | 1,99 | 1,54 |
| HMN_PC_099 | 1,45 | 0,03 | 2,56 | 1,27 | 0,14 | 0,86 | 0,80 | 0,59 | 1,35 |
| HMN_PC_100 | 1,08 | 0,02 | 2,72 | 1,00 | 1,69 | 0,78 | 0,83 | 0,23 | 1,10 |
| HMN_PC_101 | 3,24 | 0,26 | 5,20 | 0,77 | 1,85 | 0,94 | 0,90 | 3,50 | 1,40 |
| HMN_PC_102 | 0,73 | 0,14 | 6,35 | 0,92 | 1,11 | 3,65 | 0,87 | 0,35 | 3,36 |
| HMN_PC_103 | 3,37 | 0,74 | 12,91 | 2,21 | 6,96 | 2,86 | 4,40 | 2,52 | 10,58 |
| HMN_PC_104 | 0,65 | 0,11 | 3,67 | 1,77 | 0,98 | 0,98 | 1,39 | 0,27 | 1,36 |
| HMN_PC_105 | 1,32 | 0,40 | 7,93 | 0,68 | 0,61 | 0,66 | 2,54 | 0,88 | 2,74 |
| HMN_PC_106 | 1,19 | 0,37 | 8,28 | 1,16 | 1,43 | 2,31 | 3,15 | 0,63 | 5,45 |
| HMN_PC_110 | 0,48 | 0,02 | 8,52 | 0,69 | 1,56 | 2,27 | 2,88 | 0,36 | 4,50 |
| HMN_PC_111 | 1,53 | 0,04 | 4,37 | 1,60 | 0,33 | 0,28 | 0,90 | 0,76 | 1,80 |
| HMN_PC_113 | 0,67 | 0,03 | 2,59 | 0,78 | 1,17 | 1,78 | 0,85 | 0,72 | 2,29 |
| HMN_PC_114 | 3,26 | 0,57 | 5,36 | 1,38 | 2,31 | 2,66 | 1,08 | 1,50 | 1,46 |
| HMN_PC_115 | 1,11 | 0,09 | 2,66 | 0,95 | 0,93 | 0,17 | 0,84 | 0,37 | 1,25 |
| HMN_PC_117 | 0,91 | 0,02 | 5,20 | 1,76 | 2,22 | 3,41 | 1,40 | 0,49 | 2,81 |
| HMN_PC_118 | 0,15 | 0,29 | 2,36 | 0,32 | 0,31 | 0,10 | 0,79 | 0,27 | 0,71 |
| HMN_PC_119 | 1,48 | 1,81 | 3,52 | 1,01 | 2,01 | 2,16 | 0,93 | 5,97 | 4,73 |
| HMN_PC_121 | 2,34 | 0,15 | 6,04 | 1,62 | 1,09 | 5,31 | 1,27 | 0,53 | 1,11 |
| HMN_PC_124 | 0,78 | 0,18 | 2,10 | 2,63 | 0,92 | 6,62 | 0,51 | 0,37 | 1,49 |
| HMN_PC_125(-) | 0,29 | 0,34 | 0,21 | 0,05 | 0,35 | 0,08 | 0,06 | 0,02 | 0,30 |
| HMN_PC_126 | 1,31 | 0,19 | 5,05 | 1,77 | 1,42 | 2,83 | 2,24 | 1,00 | 1,79 |
| HMN_PC_127 | 1,35 | 0,41 | 0,39 | 0,17 | 0,26 | 1,26 | 0,09 | 0,77 | 1,07 |
| HMN_PC_128 | 3,81 | 0,22 | 8,02 | 0,51 | 2,53 | 1,43 | 2,83 | 0,78 | 2,51 |
| HMN_PC_129 | 0,24 | 0,04 | 7,88 | 0,75 | 2,06 | 2,23 | 4,31 | 0,03 | 3,89 |
| HMN_PC_130 | 1,94 | 1,18 | 11,63 | 2,89 | 2,43 | 12,67 | 4,39 | 0,73 | 3,14 |
| HMN_PC_131 | 1,05 | 0,52 | 3,81 | 2,94 | 0,50 | 3,61 | 1,22 | 0,71 | 1,17 |
| HMN_PC_134 | 1,12 | 0,09 | 2,15 | 1,00 | 0,79 | 5,13 | 1,27 | 0,71 | 1,42 |
| HMN_PC_135 | 2,33 | 0,25 | 7,41 | 0,98 | 6,06 | 0,73 | 5,01 | 1,76 | 3,54 |
| HMN_PC_137 | 0,46 | 0,11 | 13,38 | 1,73 | 0,75 | 0,09 | 6,72 | 0,29 | 2,92 |
| HMN_PC_138 | 1,92 | 0,06 | 3,48 | 1,59 | 0,89 | 1,74 | 2,47 | 0,98 | 1,40 |
| HMN_PC_139 | 0,64 | 0,30 | 1,01 | 0,14 | 1,14 | 0,30 | 0,28 | 0,02 | 2,40 |
| HMN_PC_140 | 0,31 | 0,15 | 7,15 | 0,89 | 5,70 | 1,69 | 2,96 | 0,14 | 4,08 |
| HMN_PC_141 | 1,34 | 0,66 | 8,49 | 0,71 | 1,30 | 3,98 | 4,76 | 0,27 | 3,61 |
| HMN_PC_142 | 0,94 | 0,07 | 15,35 | 1,28 | 0,85 | 1,31 | 4,89 | 0,56 | 4,78 |
| HMN_PC_143 | 4,62 | 0,15 | 1,93 | 0,40 | 3,08 | 1,39 | 0,83 | 0,71 | 1,13 |
| HMN_PC_145 | 3,86 | 0,31 | 4,04 | 0,59 | 1,34 | 0,53 | 1,64 | 1,68 | 1,28 |
| HMN_PC_146 | 1,97 | 0,27 | 0,30 | 0,04 | 1,95 | 0,62 | 0,22 | 0,86 | 4,89 |
| HMN_PC_147 | 0,12 | 0,05 | 1,34 | 0,36 | 1,14 | 1,39 | 0,40 | 0,12 | 0,71 |
| HMN_PC_148 | 0,43 | 0,50 | 0,01 | 0,18 | 0,33 | 0,13 | 0,07 | 1,94 | 0,70 |
| HMN_PC_149(-) | 0,01 | 0,11 | 0,10 | 0,12 | 0,20 | 0,09 | 0,08 | 0,23 | 0,39 |
| HMN_PC_152 | 1,33 | 0,35 | 3,63 | 1,32 | 1,77 | 5,70 | 1,45 | 0,47 | 1,98 |
| HMN_PC_153 | 0,06 | 0,10 | 0,97 | 0,14 | 0,25 | 0,14 | 0,35 | 0,02 | 0,48 |
| HMN_PC_156 | 2,95 | 0,32 | 11,82 | 1,75 | 5,03 | 3,28 | 3,38 | 0,72 | 3,68 |
| HMN_PC_157 | 0,56 | 0,19 | 2,95 | 1,24 | 1,54 | 4,27 | 1,79 | 0,16 | 1,95 |
| HMN_PC_158 | 2,08 | 0,07 | 6,26 | 1,44 | 1,79 | 3,88 | 1,53 | 1,47 | 1,61 |
| HMN_PC_160 | 2,07 | 0,21 | 1,07 | 0,94 | 0,68 | 1,00 | 0,37 | 1,68 | 0,60 |
| HMN_PC_161 | 1,24 | 0,17 | 1,53 | 0,45 | 0,56 | 0,90 | 0,64 | 0,61 | 0,65 |
| HMN_PC_162 | 0,27 | 0,09 | 1,34 | 0,58 | 1,98 | 0,30 | 0,44 | 0,43 | 2,62 |
| HMN_PC_164 | 1,24 | 0,09 | 3,88 | 0,59 | 1,59 | 0,86 | 1,16 | 0,25 | 1,04 |
| HMN_PC_166 | 3,73 | 0,01 | 3,43 | 1,58 | 2,58 | 2,85 | 0,55 | 4,62 | 0,69 |
| HMN_PC_167 | 3,57 | 0,28 | 3,93 | 2,19 | 1,56 | 1,91 | 0,75 | 1,58 | 1,71 |
| HMN_PC_168 | 1,06 | 0,30 | 4,31 | 2,18 | 1,09 | 3,65 | 2,54 | 0,24 | 3,02 |
| HMN_PC_169 | 0,41 | 0,05 | 4,89 | 0,79 | 1,74 | 1,35 | 2,10 | 0,79 | 2,68 |
| HMN_PC_170 | 1,17 | 0,08 | 0,16 | 0,20 | 0,47 | 1,18 | 0,14 | 1,10 | 0,94 |
| HMN_PC_171 | 0,95 | 0,20 | 2,39 | 0,56 | 0,84 | 1,26 | 0,82 | 0,62 | 1,04 |
| HMN_PC_172 | 1,23 | 0,05 | 0,34 | 0,47 | 1,17 | 1,93 | 0,17 | 2,38 | 1,60 |
| HMN_PC_173 | 2,39 | 0,07 | 2,25 | 0,82 | 0,68 | 2,73 | 0,92 | 1,28 | 1,48 |
| HMN_PC_174 | 0,35 | 0,10 | 4,66 | 0,86 | 0,35 | 1,53 | 2,40 | 0,23 | 2,42 |
| HMN_PC_209 | 0,62 | 0,35 | 1,85 | 1,37 | 0,56 | 2,13 | 0,83 | 0,41 | 0,81 |
| HMN_PC_217 | 1,28 | 0,18 | 6,63 | 0,98 | 0,74 | 0,95 | 1,86 | 0,58 | 1,01 |
| HMN_PC_218 | 0,18 | 0,05 | 0,96 | 1,01 | 1,25 | 4,50 | 0,32 | 0,22 | 0,52 |

| ID_sample | ctg_29077 | ctg_36195 | ctg_44030 | ctg_512 | ctg_57223 | ctg_61472 | ctg_61528 | ctg_63866 | ctg_9446 |
|---|---|---|---|---|---|---|---|---|---|
| HMN_PC_219 | 0,82 | 0,44 | 3,72 | 1,21 | 3,36 | 1,54 | 1,54 | 0,47 | 0,69 |
| HMN_PC_220 | 0,61 | 0,15 | 5,35 | 1,49 | 0,32 | 0,19 | 2,20 | 0,27 | 2,11 |
| HMN_PC_221 | 0,70 | 0,20 | 1,07 | 0,38 | 0,64 | 4,54 | 0,53 | 0,63 | 1,59 |
| HMN_PC_222 | 0,53 | 0,27 | 0,51 | 0,18 | 0,98 | 3,97 | 0,25 | 0,15 | 0,32 |
| HMN_PC_223 | 1,96 | 0,31 | 3,10 | 1,41 | 1,02 | 0,80 | 0,57 | 1,29 | 1,79 |
| HMN_PC_224 | 1,23 | 0,23 | 6,31 | 0,93 | 1,17 | 0,73 | 2,46 | 0,83 | 2,75 |
| HMN_PC_225 | 0,71 | 0,24 | 1,78 | 1,16 | 1,28 | 8,92 | 1,08 | 0,83 | 1,09 |
| HMN_PC_226 | 2,40 | 0,43 | 0,98 | 1,34 | 1,10 | 2,54 | 0,23 | 0,98 | 1,07 |
| HMN_PC_227 | 1,36 | 0,39 | 0,96 | 1,88 | 0,38 | 6,01 | 0,36 | 1,91 | 1,10 |
| HMN_PC_228 | 9,40 | 0,71 | 6,08 | 2,41 | 2,18 | 3,42 | 1,06 | 6,44 | 1,85 |
| HMN_PC_229 | 3,07 | 0,44 | 8,42 | 1,02 | 0,57 | 1,93 | 3,03 | 1,48 | 2,30 |
| HMN_PC_232 | 0,73 | 0,07 | 1,08 | 0,91 | 0,87 | 4,85 | 0,45 | 0,78 | 2,30 |
| HMN_PC_233 | 4,22 | 0,09 | 3,51 | 1,01 | 2,73 | 2,20 | 0,84 | 1,09 | 1,94 |
| HMN_PC_234 | 5,56 | 0,52 | 3,33 | 1,50 | 1,30 | 1,92 | 1,23 | 4,61 | 2,92 |
| HMN_PC_235 | 0,55 | 0,20 | 8,95 | 1,18 | 5,14 | 1,59 | 2,41 | 0,49 | 4,82 |
| HMN_PC_236 | 2,50 | 0,40 | 6,62 | 0,75 | 0,47 | 1,87 | 1,21 | 1,44 | 1,13 |
| HMN_PC_239 | 0,18 | 0,09 | 2,53 | 0,50 | 0,94 | 1,63 | 1,63 | 0,27 | 1,16 |
| HMN_PC_241 | 0,41 | 0,07 | 0,51 | 0,49 | 0,91 | 6,32 | 0,20 | 0,20 | 1,69 |
| HMN_PC_242 | 0,41 | 0,15 | 1,63 | 1,63 | 1,11 | 4,40 | 0,42 | 0,90 | 0,66 |
| HMN_PC_243 | 3,23 | 0,16 | 4,24 | 1,25 | 0,94 | 0,73 | 1,34 | 1,39 | 1,38 |
| HMN_PC_244 | 0,34 | 0,13 | 3,47 | 1,36 | 1,26 | 4,18 | 0,75 | 0,52 | 2,29 |
| HMN_PC_245 | 0,55 | 0,24 | 3,99 | 0,82 | 0,58 | 1,76 | 0,96 | 0,48 | 2,01 |
| HMN_PC_246 | 4,37 | 0,12 | 8,38 | 1,15 | 4,75 | 5,60 | 3,59 | 4,08 | 5,33 |
| HMN_PC_247 | 0,58 | 0,14 | 3,14 | 0,78 | 0,48 | 0,92 | 1,04 | 0,23 | 1,34 |
