## Supplemental Table S5 for "Blind exploration of the unreferenced transcriptome reveals novel RNAs for prostate cancer diagnosis"

**Table S5. Mean and Fold Change of expression of PCA3 and DE-kupl contigs in prostate normal and tumor specimens measured by NanoString across 144 prostate specimens of the Selection Set.**

| Probe_ID | Contig_ID | mean_normal | mean_tumor | FC | log2(FC) | wilcoxon_pvalue |
|---|---|---|---|---|---|---|
| P6 | ctg_111158 | 0,14 | 2,19 | 15,81 | 3,98 | 4,35E-07 |
| P2 | ctg_28650 | 0,10 | 1,06 | 10,09 | 3,33 | 6,09E-07 |
| P14 | ctg_61528 | 0,09 | 1,44 | 15,77 | 3,98 | 6,89E-07 |
| P19 | ctg_61472 | 0,12 | 2,27 | 18,72 | 4,23 | 8,47E-07 |
| P7 | ctg_117356 | 0,30 | 4,05 | 13,46 | 3,75 | 1,13E-06 |
| P9 | ctg_9446 | 0,45 | 1,92 | 4,26 | 2,09 | 1,22E-06 |
| P20 | ctg_44030 | 0,38 | 4,16 | 10,98 | 3,46 | 1,83E-06 |
| P18 | ctg_105149 | 0,16 | 1,53 | 9,49 | 3,25 | 1,90E-06 |
| P10 | ctg_25348 | 0,07 | 0,57 | 8,78 | 3,13 | 2,14E-06 |
| P15 | ctg_512 | 0,17 | 1,11 | 6,46 | 2,69 | 2,41E-06 |
| P3 | ctg_57223 | 0,23 | 1,54 | 6,58 | 2,72 | 3,17E-06 |
| P1 | ctg_17297 | 0,47 | 7,15 | 15,17 | 3,92 | 3,85E-06 |
| PCA3 | - | 0,64 | 15,55 | 24,36 | 4,61 | 5,24E-06 |
| P16 | ctg_111348 | 0,03 | 2,16 | 68,18 | 6,09 | 7,09E-06 |
| P21 | ctg_23999 | 0,11 | 1,78 | 15,75 | 3,98 | 1,19E-05 |
| P23 | ctg_29077 | 0,18 | 1,59 | 8,66 | 3,11 | 1,43E-05 |
| P11 | ctg_104447 | 0,16 | 1,39 | 8,74 | 3,13 | 2,20E-05 |
| P13 | ctg_37852 | 0,33 | 5,86 | 17,60 | 4,14 | 5,43E-05 |
| P8 | ctg_73782 | 0,29 | 3,00 | 10,38 | 3,38 | 7,59E-05 |
| P5 | ctg_123090 | 0,11 | 0,38 | 3,54 | 1,82 | 1,55E-04 |
| P12 | ctg_2815 | 0,69 | 1,63 | 2,37 | 1,24 | 6,82E-04 |
| P4 | ctg_63866 | 0,36 | 1,16 | 3,23 | 1,69 | 6,34E-03 |
| P22 | ctg_119680 | 0,11 | 0,18 | 1,59 | 0,67 | 3,87E-02 |
| P17 | ctg_36195 | 0,15 | 0,24 | 1,57 | 0,65 | 1,80E-01 |
| GAPDH | - | 140,66 | 183,16 | 1,30 | 0,38 | 4,55E-03 |
| RPL11 | - | 194,19 | 265,00 | 1,36 | 0,45 | 1,41E-02 |
| ZNF2 | - | 0,85 | 0,99 | 1,16 | 0,22 | 3,35E-02 |
| NOL7 | - | 15,26 | 16,30 | 1,07 | 0,10 | 3,07E-01 |
| GPATCH3 | - | 1,27 | 1,22 | 0,96 | -0,06 | 7,16E-01 |
| ZNF346 | - | 0,88 | 0,79 | 0,90 | -0,15 | 9,60E-01 |
