## Supplemental Table S6 for "Blind exploration of the unreferenced transcriptome reveals novel RNAs for prostate cancer diagnosis"

Table S6. PCA3 and DE-kupl contigs expression quantification assessed by the total stranded RNA-seq in 24 prostate specimens from the PAIR cohort (Discovery Set).

| Probe_ID | Contig_ID | mean_normal | mean_tumor | log2FC | wilcoxon_pvalue |
|---|---|---|---|---|---|
| P1 | ctg_17297 | 0,06 | 1,61 | 4,68 | 1,36E-06 |
| P2 | ctg_28650 | 0,00 | 0,08 | Inf | 3,72E-05 |
| P3 | ctg_57223 | 0,00 | 0,08 | Inf | 3,72E-05 |
| P4 | ctg_63866 | 0,00 | 0,13 | 6,66 | 4,13E-05 |
| P5 | ctg_123090 | 0,00 | 0,06 | 4,54 | 4,46E-05 |
| P6 | ctg_111158 | 0,01 | 0,12 | 3,79 | 4,97E-05 |
| P7 | ctg_117356 | 0,00 | 0,20 | 6,85 | 5,34E-05 |
| P8 | ctg_73782 | 0,00 | 0,08 | 4,51 | 5,74E-05 |
| P9 | ctg_9446 | 0,00 | 0,07 | 4,05 | 7,38E-05 |
| P10 | ctg_25348 | 0,00 | 0,09 | Inf | 9,11E-05 |
| P11 | ctg_104447 | 0,00 | 0,12 | Inf | 9,11E-05 |
| P12 | ctg_2815 | 0,00 | 0,07 | 4,88 | 1,44E-04 |
| P13 | ctg_37852 | 0,08 | 1,19 | 3,86 | 1,85E-04 |
| PCA3 | - | 0,01 | 0,93 | 6,42 | 2,09E-04 |
| P14 | ctg_61528 | 0,00 | 0,07 | Inf | 2,10E-04 |
| P15 | ctg_512 | 0,00 | 0,10 | Inf | 2,10E-04 |
| P16 | ctg_111348 | 0,00 | 0,15 | Inf | 2,10E-04 |
| P17 | ctg_36195 | 0,00 | 0,11 | Inf | 2,10E-04 |
| P18 | ctg_105149 | 0,01 | 0,12 | 3,32 | 2,14E-04 |
| P19 | ctg_61472 | 0,00 | 0,22 | 5,51 | 2,75E-04 |
| P20 | ctg_44030 | 0,00 | 0,17 | 6,57 | 3,04E-04 |
| P21 | ctg_23999 | 0,01 | 0,18 | 4,70 | 3,21E-04 |
| P22 | ctg_119680 | 0,01 | 0,07 | 2,96 | 3,30E-04 |
| P23 | ctg_29077 | 0,00 | 0,06 | 5,38 | 7,42E-04 |
| GAPDH | - | 8,62 | 10,54 | 0,29 | 1,32E-01 |
| RPL11 | - | 59,18 | 70,05 | 0,24 | 2,08E-01 |
| GPATCH3 | - | 0,06 | 0,07 | 0,18 | 3,05E-01 |
| NOL7 | - | 0,88 | 0,85 | -0,06 | 5,12E-01 |
| ZNF346 | - | 0,12 | 0,13 | 0,01 | 6,06E-01 |
| ZNF2 | - | 0,11 | 0,08 | -0,50 | 9,97E-01 |
