## Supplemental Table S7 for "Blind exploration of the unreferenced transcriptome reveals novel RNAs for prostate cancer diagnosis"

**Table S7. Clinico-pathological characteristics and recurrence status of the prostate specimens from the TCGA-PRAD cohort (Validation Set).**

| Barcode | Tissue_type | State | gleason_sco | TNM | Risk group | Recurrence |
|---|---|---|---|---|---|---|
| TCGA-CH-5761-11A-01R-1580-07 | normal | Live | - | - | - | - |
| TCGA-CH-5767-11B-01R-1789-07 | normal | Live | - | - | - | - |
| TCGA-CH-5768-11A-01R-1580-07 | normal | Live | - | - | - | - |
| TCGA-CH-5769-11A-01R-1580-07 | normal | Live | - | - | - | - |
| TCGA-EJ-7115-11A-01R-2118-07 | normal | Live | - | - | - | - |
| TCGA-EJ-7123-11A-01R-1965-07 | normal | Live | - | - | - | - |
| TCGA-EJ-7125-11A-01R-1965-07 | normal | Live | - | - | - | - |
| TCGA-EJ-7314-11A-01R-2118-07 | normal | Live | - | - | - | - |
| TCGA-EJ-7315-11A-01R-2118-07 | normal | Live | - | - | - | - |
| TCGA-EJ-7317-11A-01R-2118-07 | normal | Live | - | - | - | - |
| TCGA-EJ-7321-11A-01R-2263-07 | normal | Live | - | - | - | - |
| TCGA-EJ-7327-11A-01R-2118-07 | normal | Live | - | - | - | - |
| TCGA-EJ-7328-11A-01R-2118-07 | normal | Live | - | - | - | - |
| TCGA-EJ-7330-11A-01R-2118-07 | normal | Live | - | - | - | - |
| TCGA-EJ-7331-11A-01R-2118-07 | normal | Live | - | - | - | - |
| TCGA-EJ-7781-11A-01R-2118-07 | normal | Live | - | - | - | - |
| TCGA-EJ-7782-11A-01R-2118-07 | normal | Live | - | - | - | - |
| TCGA-EJ-7783-11A-01R-2118-07 | normal | Live | - | - | - | - |
| TCGA-EJ-7784-11A-01R-2118-07 | normal | Live | - | - | - | - |
| TCGA-EJ-7785-11A-01R-2118-07 | normal | Live | - | - | - | - |
| TCGA-EJ-7786-11A-01R-2118-07 | normal | Live | - | - | - | - |
| TCGA-EJ-7789-11A-01R-2118-07 | normal | Live | - | - | - | - |
| TCGA-EJ-7792-11A-01R-2118-07 | normal | Live | - | - | - | - |
| TCGA-EJ-7793-11A-01R-2263-07 | normal | Live | - | - | - | - |
| TCGA-EJ-7794-11A-01R-2118-07 | normal | Live | - | - | - | - |
| TCGA-EJ-7797-11A-01R-2263-07 | normal | Live | - | - | - | - |
| TCGA-EJ-A8FO-11A-11R-A36G-07 | normal | Live | - | - | - | - |
| TCGA-G9-6333-11A-01R-1965-07 | normal | Live | - | - | - | - |
| TCGA-G9-6342-11A-02R-1965-07 | normal | Live | - | - | - | - |
| TCGA-G9-6348-11A-01R-1789-07 | normal | Live | - | - | - | - |
| TCGA-G9-6351-11A-01R-1965-07 | normal | Live | - | - | - | - |
| TCGA-G9-6356-11A-01R-1789-07 | normal | Live | - | - | - | - |
| TCGA-G9-6362-11A-01R-1789-07 | normal | Live | - | - | - | - |
| TCGA-G9-6363-11A-01R-1789-07 | normal | Live | - | - | - | - |
| TCGA-G9-6365-11A-01R-1789-07 | normal | Live | - | - | - | - |
| TCGA-G9-6384-11A-01R-1858-07 | normal | Live | - | - | - | - |
| TCGA-G9-6496-11A-01R-1789-07 | normal | Live | - | - | - | - |
| TCGA-G9-6499-11A-02R-1965-07 | normal | Live | - | - | - | - |
| TCGA-HC-7211-11A-01R-2118-07 | normal | Live | - | - | - | - |
| TCGA-HC-7737-11A-02R-2118-07 | normal | Live | - | - | - | - |
| TCGA-HC-7738-11A-01R-2118-07 | normal | Live | - | - | - | - |
| TCGA-HC-7740-11A-01R-2118-07 | normal | Live | - | - | - | - |

| | | | | | | |
|---|---|---|---|---|---|---|
| TCGA-HC-7742-11A-01R-2118-07 | normal | Live | - | - | - | - |
| **Barcode** | **Tissue_type** | **State** | **gleason_sc** | **TNM** | **Risk group** | **Recurrence** |
| TCGA-HC-7745-11A-01R-2118-07 | normal | Live | - | - | - | - |
| TCGA-HC-7747-11A-01R-2118-07 | normal | Live | - | - | - | - |
| TCGA-HC-7752-11A-01R-2118-07 | normal | Live | - | - | - | - |
| TCGA-HC-7819-11A-01R-2118-07 | normal | Live | - | - | - | - |
| TCGA-HC-8258-11A-01R-2263-07 | normal | Live | - | - | - | - |
| TCGA-HC-8259-11A-01R-2263-07 | normal | Live | - | - | - | - |
| TCGA-HC-8260-11A-01R-2263-07 | normal | Live | - | - | - | - |
| TCGA-HC-8262-11A-01R-2263-07 | normal | Live | - | - | - | - |
| TCGA-J4-A83J-11A-11R-A36G-07 | normal | Live | - | - | - | - |
| TCGA-2A-A8VT-01A-11R-A37L-07 | tumor | Live | 9 | T4 | high | NO |
| TCGA-2A-A8VX-01A-11R-A37L-07 | tumor | Live | 8 | T3b | high | NO |
| TCGA-4L-AA1F-01A-11R-A41O-07 | tumor | Live | 8 | T3b | high | NO |
| TCGA-CH-5741-01A-11R-1580-07 | tumor | Live | 9 | T3b | high | NO |
| TCGA-CH-5752-01A-11R-1580-07 | tumor | Live | 8 | T3a | high | NO |
| TCGA-CH-5762-01A-11R-1580-07 | tumor | Live | 7 | T3b | high | NO |
| TCGA-CH-5772-01A-11R-1580-07 | tumor | Live | 9 | T3a | high | NO |
| TCGA-EJ-5495-01A-01R-1580-07 | tumor | Live | 8 | T3b | high | NO |
| TCGA-EJ-5501-01A-01R-1580-07 | tumor | Live | 7 | T3b | high | NO |
| TCGA-EJ-5503-01A-01R-1580-07 | tumor | Live | 8 | T3a | high | NO |
| TCGA-EJ-5506-01A-01R-1580-07 | tumor | Live | 8 | T3b | high | NO |
| TCGA-EJ-5507-01A-01R-1580-07 | tumor | Live | 9 | T3b | high | NO |
| TCGA-EJ-5514-01A-01R-1580-07 | tumor | Live | 9 | T2c | high | NO |
| TCGA-EJ-5519-01A-01R-1580-07 | tumor | Live | 8 | T3b | high | NO |
| TCGA-EJ-7312-01B-21R-A32O-07 | tumor | Live | 7 | T3b | high | NO |
| TCGA-EJ-7314-01A-31R-2118-07 | tumor | Live | 7 | T3b | high | NO |
| TCGA-EJ-7325-01B-11R-A32O-07 | tumor | Live | 7 | T3b | high | NO |
| TCGA-EJ-7327-01A-11R-2118-07 | tumor | Live | 7 | T3b | high | NO |
| TCGA-EJ-7782-01A-11R-2118-07 | tumor | Live | 8 | T2c | high | NO |
| TCGA-EJ-7788-01A-11R-2118-07 | tumor | Live | 7 | T3b | high | NO |
| TCGA-EJ-8468-01A-21R-2403-07 | tumor | Live | 8 | T3a | high | NO |
| TCGA-EJ-8469-01A-11R-2403-07 | tumor | Live | 9 | T3a | high | NO |
| TCGA-EJ-8474-01A-11R-2403-07 | tumor | Live | 8 | T3a | high | NO |
| TCGA-EJ-A46B-01A-31R-A250-07 | tumor | Live | 8 | T3a | high | NO |
| TCGA-EJ-A46D-01A-21R-A32Y-07 | tumor | Live | 8 | T2c | high | NO |
| TCGA-EJ-A46E-01A-31R-A250-07 | tumor | Live | 8 | [Not Available] | high | NO |
| TCGA-EJ-A46G-01A-31R-A26U-07 | tumor | Live | 8 | T3a | high | NO |
| TCGA-EJ-A65B-01A-12R-A30B-07 | tumor | Live | 9 | T3b | high | NO |
| TCGA-EJ-A65D-01A-11R-A30B-07 | tumor | Live | 8 | T3a | high | NO |
| TCGA-EJ-A65E-01A-11R-A29R-07 | tumor | Live | 7 | T3b | high | NO |
| TCGA-EJ-A65G-01A-21R-A29R-07 | tumor | Live | 8 | T2c | high | NO |
| TCGA-EJ-A65J-01A-11R-A311-07 | tumor | Live | 9 | T3a | high | NO |
| TCGA-EJ-A7NJ-01A-22R-A352-07 | tumor | Live | 8 | T2c | high | NO |
| TCGA-EJ-A7NM-01A-21R-A33R-07 | tumor | Live | 9 | T3b | high | NO |

| | | | | | | |
|---|---|---|---|---|---|---|
| TCGA-EJ-A8FU-01A-11R-A36G-07 | tumor | Live | 8 | T3a | high | NO |
| **Barcode** | **Tissue_type** | **State** | **gleason_sc** | **TNM** | **Risk group** | **Recurrence** |
| TCGA-FC-7961-01A-11R-A29R-07 | tumor | Live | 9 | T3a | high | NO |
| TCGA-FC-A4JI-01A-11R-A250-07 | tumor | Live | 8 | T3a | high | NO |
| TCGA-FC-A5OB-01A-11R-A29R-07 | tumor | Live | 9 | T3b | high | NO |
| TCGA-FC-A66V-01A-21R-A30B-07 | tumor | Live | 7 | T3b | high | NO |
| TCGA-G9-6356-01A-11R-1789-07 | tumor | Live | 9 | T3b | high | NO |
| TCGA-G9-6363-01A-21R-1789-07 | tumor | Live | 7 | T4 | high | NO |
| TCGA-G9-6365-01A-11R-1789-07 | tumor | Live | 7 | T4 | high | NO |
| TCGA-G9-6367-01A-11R-1789-07 | tumor | Live | 9 | T3a | high | NO |
| TCGA-G9-6370-01A-11R-1789-07 | tumor | Live | 7 | T3b | high | NO |
| TCGA-G9-6379-01A-11R-A31N-07 | tumor | Live | 7 | T3b | high | NO |
| TCGA-G9-6494-01A-11R-1789-07 | tumor | Live | 7 | T4 | high | NO |
| TCGA-G9-6499-01A-12R-1965-07 | tumor | Live | 9 | T3a | high | NO |
| TCGA-G9-7510-01A-11R-2263-07 | tumor | Live | 8 | T3a | high | NO |
| TCGA-G9-7521-01A-11R-2263-07 | tumor | Live | 8 | T3a | high | NO |
| TCGA-G9-7523-01A-11R-2263-07 | tumor | Live | 10 | T3a | high | NO |
| TCGA-G9-A9S4-01A-11R-A41O-07 | tumor | Live | 8 | T3b | high | NO |
| TCGA-G9-A9S7-01A-11R-A41O-07 | tumor | Live | 8 | T3a | high | NO |
| TCGA-HC-7081-01A-11R-1965-07 | tumor | Live | 9 | T3b | high | NO |
| TCGA-HC-7744-01A-11R-2118-07 | tumor | Live | 7 | T3b | high | NO |
| TCGA-HC-7745-01A-11R-2118-07 | tumor | Live | 7 | T3b | high | NO |
| TCGA-HC-7817-01B-11R-A29R-07 | tumor | Live | 7 | T3b | high | NO |
| TCGA-HC-7819-01A-11R-2118-07 | tumor | Live | 8 | T2c | high | NO |
| TCGA-HC-8257-01A-11R-2263-07 | tumor | Live | 7 | T3b | high | NO |
| TCGA-HC-8262-01A-11R-2263-07 | tumor | Live | 8 | T2c | high | NO |
| TCGA-HC-8264-01B-11R-2403-07 | tumor | Live | 9 | T3b | high | NO |
| TCGA-HC-8265-01A-11R-2263-07 | tumor | Live | 8 | T3a | high | NO |
| TCGA-HC-8265-01B-04R-2302-07 | tumor | Live | 8 | T3a | high | NO |
| TCGA-HC-8265-01B-04R-2302-07 | tumor | Live | 8 | T3a | high | NO |
| TCGA-HC-8266-01A-11R-2263-07 | tumor | Live | 8 | T3b | high | NO |
| TCGA-HC-A48F-01A-11R-A250-07 | tumor | Live | 8 | T2c | high | NO |
| TCGA-HC-A4ZV-01A-11R-A26U-07 | tumor | Live | 9 | T3b | high | NO |
| TCGA-HC-A631-01A-11R-A29R-07 | tumor | Live | 9 | T2c | high | NO |
| TCGA-HC-A632-01A-11R-A29R-07 | tumor | Live | 9 | T3a | high | NO |
| TCGA-HC-A76W-01A-11R-A33R-07 | tumor | Live | 7 | T3b | high | NO |
| TCGA-J4-8198-01A-11R-2263-07 | tumor | Live | 7 | T3b | high | NO |
| TCGA-J4-A6G1-01A-11R-A311-07 | tumor | Live | 8 | T3a | high | NO |
| TCGA-J9-A52C-01A-11R-A26U-07 | tumor | Live | 9 | T3b | high | NO |
| TCGA-J9-A52D-01A-11R-A29R-07 | tumor | Live | 9 | T3a | high | NO |
| TCGA-J9-A8CK-01A-11R-A352-07 | tumor | Live | 9 | T3a | high | NO |
| TCGA-KK-A59V-01A-11R-A29R-07 | tumor | Live | 9 | T3b | high | NO |
| TCGA-KK-A59Y-01A-11R-A26U-07 | tumor | Live | 9 | T3a | high | NO |
| TCGA-KK-A6E1-01A-11R-A311-07 | tumor | Live | 9 | T3b | high | NO |
| TCGA-KK-A6E5-01A-11R-A311-07 | tumor | Live | 7 | T3b | high | NO |

| | | | | | | |
|---|---|---|---|---|---|---|
| TCGA-KK-A6E6-01A-11R-A311-07 | tumor | Live | 9 | T3a | high | NO |
| **Barcode** | **Tissue_type** | **State** | **gleason_sc** | **TNM** | **Risk group** | **Recurrence** |
| TCGA-KK-A6E8-01A-11R-A31N-07 | tumor | Live | 9 | T2c | high | NO |
| TCGA-KK-A8I8-01A-11R-A36G-07 | tumor | Live | 9 | T3b | high | NO |
| TCGA-KK-A8IA-01A-11R-A36G-07 | tumor | Live | 9 | T3b | high | NO |
| TCGA-KK-A8ID-01A-11R-A36G-07 | tumor | Live | 9 | T3b | high | NO |
| TCGA-KK-A8IK-01A-11R-A36G-07 | tumor | Live | 9 | T2c | high | NO |
| TCGA-M7-A723-01A-12R-A32O-07 | tumor | Live | 7 | T3b | high | NO |
| TCGA-M7-A725-01A-12R-A32O-07 | tumor | Live | 7 | T3b | high | NO |
| TCGA-MG-AAMC-01A-11R-A41O-07 | tumor | Live | 9 | T3a | high | NO |
| TCGA-SU-A7E7-01A-22R-A33R-07 | tumor | Live | 8 | T2b | high | NO |
| TCGA-V1-A8MG-01A-11R-A36G-07 | tumor | Live | 7 | T3b | high | NO |
| TCGA-V1-A8WV-01A-11R-A37L-07 | tumor | Live | 9 | T3b | high | NO |
| TCGA-V1-A9O9-01A-11R-A41O-07 | tumor | Live | 8 | T3b | high | NO |
| TCGA-V1-A9OA-01A-11R-A41O-07 | tumor | Live | 9 | T3b | high | NO |
| TCGA-V1-A9OH-01A-11R-A41O-07 | tumor | Live | 8 | T2b | high | NO |
| TCGA-V1-A9OX-01A-11R-A41O-07 | tumor | Live | 8 | T2c | high | NO |
| TCGA-V1-A9Z8-01A-11R-A41O-07 | tumor | Live | 9 | T3b | high | NO |
| TCGA-V1-A9Z9-01A-21R-A41O-07 | tumor | Live | 9 | T2c | high | NO |
| TCGA-V1-A9ZG-01A-11R-A41O-07 | tumor | Live | 9 | T2c | high | NO |
| TCGA-V1-A9ZK-01A-11R-A41O-07 | tumor | Live | 8 | T3b | high | NO |
| TCGA-VN-A88I-01A-11R-A352-07 | tumor | Live | 8 | T3a | high | NO |
| TCGA-VN-A88K-01A-11R-A352-07 | tumor | Live | 8 | T3a | high | NO |
| TCGA-VN-A88Q-01A-11R-A352-07 | tumor | Live | 8 | T2c | high | NO |
| TCGA-VN-A943-01A-11R-A41O-07 | tumor | Live | 8 | T2c | high | NO |
| TCGA-VP-A872-01A-11R-A352-07 | tumor | Live | 8 | T3a | high | NO |
| TCGA-VP-A876-01A-11R-A352-07 | tumor | Live | 8 | T2c | high | NO |
| TCGA-VP-A879-01A-11R-A352-07 | tumor | Live | 9 | T2b | high | NO |
| TCGA-VP-A87H-01A-11R-A352-07 | tumor | Live | 9 | T3b | high | NO |
| TCGA-VP-AA1N-01A-31R-A41O-07 | tumor | Live | 9 | T3a | high | NO |
| TCGA-WW-A8ZI-01A-11R-A37L-07 | tumor | Live | 8 | T2c | high | NO |
| TCGA-XJ-A83G-01A-11R-A352-07 | tumor | Live | 7 | T3b | high | NO |
| TCGA-XJ-A9DI-01A-11R-A37L-07 | tumor | Live | 9 | T3b | high | NO |
| TCGA-XJ-A9DK-01A-11R-A37L-07 | tumor | Live | 8 | T2c | high | NO |
| TCGA-XK-AAIR-01A-11R-A41O-07 | tumor | Live | 8 | T2c | high | NO |
| TCGA-XK-AAIV-01A-11R-A41O-07 | tumor | Live | 10 | T3b | high | NO |
| TCGA-XK-AAJ3-01A-11R-A41O-07 | tumor | Live | 8 | T3a | high | NO |
| TCGA-XQ-A8TA-01A-11R-A36G-07 | tumor | Live | 10 | [Not Available] | high | NO |
| TCGA-XQ-A8TB-01A-11R-A36G-07 | tumor | Live | 9 | T3b | high | NO |
| TCGA-Y6-A9XI-01A-11R-A41O-07 | tumor | Live | 8 | [Discrepancy] | high | NO |
| TCGA-YL-A8HJ-01A-11R-A36G-07 | tumor | Live | 9 | T3b | high | NO |
| TCGA-YL-A8HL-01A-11R-A36G-07 | tumor | Live | 9 | T3b | high | NO |
| TCGA-YL-A8SA-01A-21R-A37L-07 | tumor | Live | 8 | T3b | high | NO |
| TCGA-YL-A8SF-01A-11R-A37L-07 | tumor | Live | 8 | T3b | high | NO |
| TCGA-YL-A8SK-01B-21R-A37L-07 | tumor | Live | 9 | T3b | high | NO |

| | | | | | | |
|---|---|---|---|---|---|---|
| TCGA-YL-A8SL-01B-21R-A37L-07 | tumor | Live | 8 | T3b | high | NO |
| **Barcode** | **Tissue_type** | **State** | **gleason_sc** | **TNM** | **Risk group** | **Recurrence** |
| TCGA-YL-A8SO-01B-31R-A37L-07 | tumor | Live | 9 | T3a | high | NO |
| TCGA-YL-A8SR-01B-11R-A37L-07 | tumor | Live | 9 | T3a | high | NO |
| TCGA-YL-A9WH-01A-11R-A37L-07 | tumor | Live | 9 | T3b | high | NO |
| TCGA-YL-A9WI-01A-11R-A37L-07 | tumor | Live | 9 | T3a | high | NO |
| TCGA-ZG-A8QW-01A-11R-A37L-07 | tumor | Live | 9 | T3b | high | NO |
| TCGA-ZG-A8QY-01A-11R-A37L-07 | tumor | Live | 9 | T3b | high | NO |
| TCGA-ZG-A8QZ-01A-11R-A37L-07 | tumor | Live | 9 | T3b | high | NO |
| TCGA-ZG-A9KY-01A-11R-A41O-07 | tumor | Live | 9 | T4 | high | NO |
| TCGA-ZG-A9L0-01A-11R-A41O-07 | tumor | Live | 9 | T3b | high | NO |
| TCGA-ZG-A9L1-01A-11R-A41O-07 | tumor | Live | 9 | T3b | high | NO |
| TCGA-ZG-A9L4-01A-11R-A41O-07 | tumor | Live | 9 | T3b | high | NO |
| TCGA-ZG-A9L5-01A-12R-A41O-07 | tumor | Live | 9 | T4 | high | NO |
| TCGA-ZG-A9LB-01A-11R-A41O-07 | tumor | Live | 9 | T3b | high | NO |
| TCGA-ZG-A9LM-01A-11R-A41O-07 | tumor | Live | 9 | T3a | high | NO |
| TCGA-ZG-A9LN-01A-11R-A41O-07 | tumor | Live | 9 | T3b | high | NO |
| TCGA-ZG-A9LS-01A-12R-A41O-07 | tumor | Live | 9 | T3b | high | NO |
| TCGA-ZG-A9LU-01A-11R-A41O-07 | tumor | Live | 9 | T3a | high | NO |
| TCGA-ZG-A9LY-01A-11R-A41O-07 | tumor | Live | 9 | T3b | high | NO |
| TCGA-ZG-A9M4-01A-11R-A41O-07 | tumor | Live | 9 | T3b | high | NO |
| TCGA-ZG-A9MC-01A-31R-A41O-07 | tumor | Live | 9 | T3b | high | NO |
| TCGA-ZG-A9N3-01A-11R-A41O-07 | tumor | Live | 9 | T3b | high | NO |
| TCGA-ZG-A9ND-01A-11R-A41O-07 | tumor | Live | 9 | T3a | high | NO |
| TCGA-ZG-A9NI-01A-11R-A41O-07 | tumor | Live | 9 | T3b | high | NO |
| TCGA-2A-A8VO-01A-11R-A37L-07 | tumor | Live | 6 | T3a | intermediate | NO |
| TCGA-2A-A8W1-01A-11R-A37L-07 | tumor | Live | 7 | T3a | intermediate | NO |
| TCGA-2A-AAYF-01A-11R-A41O-07 | tumor | Live | 7 | T3a | intermediate | NO |
| TCGA-2A-AAYU-01A-11R-A41O-07 | tumor | Live | 6 | T3a | intermediate | NO |
| TCGA-CH-5739-01A-11R-1580-07 | tumor | Live | 7 | T3a | intermediate | NO |
| TCGA-CH-5763-01A-11R-1580-07 | tumor | Live | 7 | T3a | intermediate | NO |
| TCGA-CH-5765-01A-11R-1580-07 | tumor | Live | 7 | T3a | intermediate | NO |
| TCGA-CH-5767-01A-11R-1789-07 | tumor | Live | 7 | T2c | intermediate | NO |
| TCGA-CH-5768-01A-11R-1580-07 | tumor | Live | 6 | T3a | intermediate | NO |
| TCGA-CH-5771-01A-21R-1580-07 | tumor | Live | 7 | T2c | intermediate | NO |
| TCGA-CH-5789-01A-11R-1580-07 | tumor | Live | 7 | T3a | intermediate | NO |
| TCGA-EJ-5494-01A-01R-1580-07 | tumor | Live | 7 | T3a | intermediate | NO |
| TCGA-EJ-5498-01A-01R-1580-07 | tumor | Live | 7 | T2c | intermediate | NO |
| TCGA-EJ-5499-01A-01R-1580-07 | tumor | Live | 7 | T3a | intermediate | NO |
| TCGA-EJ-5502-01A-01R-1580-07 | tumor | Live | 7 | T2c | intermediate | NO |
| TCGA-EJ-5505-01A-01R-1580-07 | tumor | Live | 7 | T2c | intermediate | NO |
| TCGA-EJ-5508-01A-02R-1580-07 | tumor | Live | 7 | T3a | intermediate | NO |
| TCGA-EJ-5510-01A-01R-1580-07 | tumor | Live | 7 | T2c | intermediate | NO |
| TCGA-EJ-5511-01A-01R-1580-07 | tumor | Live | 7 | T3a | intermediate | NO |
| TCGA-EJ-5515-01A-01R-1580-07 | tumor | Live | 7 | T3a | intermediate | NO |

| | | | | | | |
|---|---|---|---|---|---|---|
| TCGA-EJ-5516-01A-01R-1580-07 | tumor | Live | 7 | T3a | intermediate | NO |
| **Barcode** | **Tissue_type** | **State** | **gleason_sc** | **TNM** | **Risk group** | **Recurrence** |
| TCGA-EJ-5521-01A-01R-1580-07 | tumor | Live | 7 | T3a | intermediate | NO |
| TCGA-EJ-5527-01A-01R-1580-07 | tumor | Live | 7 | T3a | intermediate | NO |
| TCGA-EJ-5531-01A-01R-1580-07 | tumor | Live | 7 | T3a | intermediate | NO |
| TCGA-EJ-5532-01A-01R-1580-07 | tumor | Live | 7 | T3a | intermediate | NO |
| TCGA-EJ-5542-01A-01R-1580-07 | tumor | Live | 7 | T3a | intermediate | NO |
| TCGA-EJ-7115-01A-11R-2118-07 | tumor | Live | 7 | T3a | intermediate | NO |
| TCGA-EJ-7315-01A-31R-2118-07 | tumor | Live | 7 | T3a | intermediate | NO |
| TCGA-EJ-7321-01A-31R-2263-07 | tumor | Live | 6 | T3a | intermediate | NO |
| TCGA-EJ-7328-01A-31R-2118-07 | tumor | Live | 7 | T3a | intermediate | NO |
| TCGA-EJ-7783-01A-11R-2118-07 | tumor | Live | 7 | T3a | intermediate | NO |
| TCGA-EJ-7784-01A-11R-2118-07 | tumor | Live | 7 | T2c | intermediate | NO |
| TCGA-EJ-7785-01A-11R-2118-07 | tumor | Live | 7 | T3a | intermediate | NO |
| TCGA-EJ-7789-01A-11R-2118-07 | tumor | Live | 7 | T3a | intermediate | NO |
| TCGA-EJ-8470-01A-11R-2403-07 | tumor | Live | 7 | T2c | intermediate | NO |
| TCGA-EJ-A46I-01A-12R-A26U-07 | tumor | Live | 7 | T3a | intermediate | NO |
| TCGA-EJ-A7NG-01A-31R-A33R-07 | tumor | Live | 7 | T3a | intermediate | NO |
| TCGA-EJ-A7NK-01A-12R-A352-07 | tumor | Live | 7 | T3a | intermediate | NO |
| TCGA-EJ-A8FO-01A-21R-A36G-07 | tumor | Live | 7 | T3a | intermediate | NO |
| TCGA-FC-7708-01A-11R-2118-07 | tumor | Live | 7 | T3a | intermediate | NO |
| TCGA-FC-A6HD-01A-11R-A31N-07 | tumor | Live | 7 | T3a | intermediate | NO |
| TCGA-FC-A8O0-01A-41R-A37L-07 | tumor | Live | 6 | T3a | intermediate | NO |
| TCGA-G9-6333-01A-12R-1965-07 | tumor | Live | 7 | T2c | intermediate | NO |
| TCGA-G9-6338-01A-12R-1965-07 | tumor | Live | 7 | T3a | intermediate | NO |
| TCGA-G9-6342-01A-11R-1965-07 | tumor | Live | 6 | T3a | intermediate | NO |
| TCGA-G9-6353-01A-11R-1965-07 | tumor | Live | 7 | T3a | intermediate | NO |
| TCGA-G9-6361-01A-21R-1965-07 | tumor | Live | 7 | T3a | intermediate | NO |
| TCGA-G9-6364-01A-21R-1789-07 | tumor | Live | 7 | T3a | intermediate | NO |
| TCGA-G9-6369-01A-21R-1965-07 | tumor | Live | 7 | T2c | intermediate | NO |
| TCGA-G9-6373-01A-11R-1789-07 | tumor | Live | 7 | T3a | intermediate | NO |
| TCGA-G9-6384-01A-11R-1789-07 | tumor | Live | 7 | T3a | intermediate | NO |
| TCGA-G9-6496-01A-11R-1789-07 | tumor | Live | 7 | T2c | intermediate | NO |
| TCGA-G9-7525-01A-31R-2263-07 | tumor | Live | 7 | T2c | intermediate | NO |
| TCGA-H9-A6BY-01A-11R-A30B-07 | tumor | Live | 7 | T3a | intermediate | NO |
| TCGA-HC-7077-01A-11R-1965-07 | tumor | Live | 6 | T3a | intermediate | NO |
| TCGA-HC-7078-01A-11R-2118-07 | tumor | Live | 7 | T3a | intermediate | NO |
| TCGA-HC-7233-01A-11R-2118-07 | tumor | Live | 7 | T2a | intermediate | NO |
| TCGA-HC-7749-01A-11R-2118-07 | tumor | Live | 7 | T3a | intermediate | NO |
| TCGA-HC-8216-01A-11R-A29R-07 | tumor | Live | 7 | T3a | intermediate | NO |
| TCGA-HC-A6AN-01A-11R-A30B-07 | tumor | Live | 7 | T3a | intermediate | NO |
| TCGA-HC-A6AO-01A-11R-A30B-07 | tumor | Live | 7 | T2c | intermediate | NO |
| TCGA-HC-A6HX-01A-11R-A31N-07 | tumor | Live | 7 | T2c | intermediate | NO |
| TCGA-HI-7169-01A-11R-2118-07 | tumor | Live | 7 | T3a | intermediate | NO |
| TCGA-J4-A67K-01A-21R-A30B-07 | tumor | Live | 7 | T2c | intermediate | NO |

| | | | | | | |
|---|---|---|---|---|---|---|
| TCGA-J4-A67L-01A-11R-A30B-07 | tumor | Live | 7 | T3a | intermediate | NO |
| **Barcode** | **Tissue_type** | **State** | **gleason_sc** | **TNM** | **Risk group** | **Recurrence** |
| TCGA-J4-A67M-01A-11R-A30B-07 | tumor | Live | 7 | T3a | intermediate | NO |
| TCGA-J4-A67R-01A-21R-A30B-07 | tumor | Live | 7 | T2c | intermediate | NO |
| TCGA-J4-A83I-01A-11R-A36G-07 | tumor | Live | 7 | T3a | intermediate | NO |
| TCGA-J9-A8CN-01A-11R-A352-07 | tumor | Live | 6 | T3a | intermediate | NO |
| TCGA-J9-A8CP-01A-11R-A352-07 | tumor | Live | 7 | T3a | intermediate | NO |
| TCGA-KC-A7F5-01A-11R-A33R-07 | tumor | Live | 7 | T2c | intermediate | NO |
| TCGA-KC-A7FA-01A-21R-A33R-07 | tumor | Live | 7 | T3a | intermediate | NO |
| TCGA-KC-A7FD-01A-11R-A33R-07 | tumor | Live | 7 | T3a | intermediate | NO |
| TCGA-KK-A59Z-01A-12R-A26U-07 | tumor | Live | 7 | T2c | intermediate | NO |
| TCGA-KK-A8I6-01A-11R-A36G-07 | tumor | Live | 7 | T3a | intermediate | NO |
| TCGA-KK-A8IG-01A-11R-A36G-07 | tumor | Live | 7 | T3a | intermediate | NO |
| TCGA-KK-A8IH-01A-11R-A36G-07 | tumor | Live | 7 | T3a | intermediate | NO |
| TCGA-KK-A8IM-01A-11R-A36G-07 | tumor | Live | 7 | T2c | intermediate | NO |
| TCGA-M7-A71Z-01A-12R-A32O-07 | tumor | Live | 7 | T3a | intermediate | NO |
| TCGA-TP-A8TT-01A-12R-A41O-07 | tumor | Live | 7 | T3a | intermediate | NO |
| TCGA-TP-A8TV-01A-11R-A41O-07 | tumor | Live | 7 | T2c | intermediate | NO |
| TCGA-V1-A8MF-01A-11R-A36G-07 | tumor | Live | 6 | T3a | intermediate | NO |
| TCGA-V1-A8WL-01A-11R-A37L-07 | tumor | Live | 7 | T3a | intermediate | NO |
| TCGA-V1-A9OQ-01A-11R-A41O-07 | tumor | Live | 6 | T3a | intermediate | NO |
| TCGA-V1-A9OY-01A-11R-A41O-07 | tumor | Live | 7 | T3a | intermediate | NO |
| TCGA-VN-A88N-01A-11R-A36G-07 | tumor | Live | 7 | T2c | intermediate | NO |
| TCGA-VP-A875-01A-31R-A352-07 | tumor | Live | 7 | T3a | intermediate | NO |
| TCGA-VP-A87C-01A-11R-A352-07 | tumor | Live | 7 | T2c | intermediate | NO |
| TCGA-VP-A87J-01A-11R-A352-07 | tumor | Live | 7 | T3a | intermediate | NO |
| TCGA-XA-A8JR-01A-11R-A36G-07 | tumor | Live | 7 | T2c | intermediate | NO |
| TCGA-XJ-A83F-01A-11R-A352-07 | tumor | Live | 7 | T3a | intermediate | NO |
| TCGA-XK-AAJA-01A-11R-A41O-07 | tumor | Live | 7 | T3a | intermediate | NO |
| TCGA-XK-AAJP-01A-11R-A41O-07 | tumor | Live | 7 | T3a | intermediate | NO |
| TCGA-XK-AAJT-01A-11R-A41O-07 | tumor | Live | 7 | T3a | intermediate | NO |
| TCGA-XK-AAJU-01A-11R-A41O-07 | tumor | Live | 7 | T2c | intermediate | NO |
| TCGA-XK-AAK1-01A-11R-A41O-07 | tumor | Live | 7 | T2c | intermediate | NO |
| TCGA-Y6-A8TL-01A-21R-A37L-07 | tumor | Live | 6 | T3a | intermediate | NO |
| TCGA-YL-A8SH-01B-11R-A37L-07 | tumor | Live | 7 | T3a | intermediate | NO |
| TCGA-2A-A8VL-01A-21R-A37L-07 | tumor | Live | 6 | T2b | low | NO |
| TCGA-2A-A8VV-01A-11R-A37L-07 | tumor | Live | 6 | T2b | low | NO |
| TCGA-2A-AAYO-01A-11R-A41O-07 | tumor | Live | 6 | T2c | low | NO |
| TCGA-CH-5738-01A-11R-1580-07 | tumor | Live | 6 | [Discrepancy] | low | NO |
| TCGA-CH-5746-01A-11R-1580-07 | tumor | Live | 7 | T2c | low | NO |
| TCGA-CH-5750-01A-11R-1580-07 | tumor | Live | 7 | T2c | low | NO |
| TCGA-CH-5790-01A-11R-1580-07 | tumor | Live | 7 | T2c | low | NO |
| TCGA-CH-5794-01A-11R-1580-07 | tumor | Live | 7 | T2b | low | NO |
| TCGA-EJ-5496-01A-01R-1580-07 | tumor | Live | 7 | T2c | low | NO |
| TCGA-EJ-5497-01A-02R-1580-07 | tumor | Live | 7 | T2c | low | NO |

| | | | | | | |
|---|---|---|---|---|---|---|
| TCGA-EJ-5509-01A-01R-1580-07 | tumor | Live | 7 | T2c | low | NO |
| **Barcode** | **Tissue_type** | **State** | **gleason_sc** | **TNM** | **Risk group** | **Recurrence** |
| TCGA-EJ-5512-01A-01R-1580-07 | tumor | Live | 7 | T2c | low | NO |
| TCGA-EJ-5517-01A-01R-1580-07 | tumor | Live | 6 | T2c | low | NO |
| TCGA-EJ-5522-01A-01R-1580-07 | tumor | Live | 7 | T2c | low | NO |
| TCGA-EJ-5530-01A-01R-1580-07 | tumor | Live | 7 | T2c | low | NO |
| TCGA-EJ-7123-01A-11R-1965-07 | tumor | Live | 7 | T2c | low | NO |
| TCGA-EJ-7125-01A-11R-1965-07 | tumor | Live | 7 | T2c | low | NO |
| TCGA-EJ-7218-01B-11R-A32O-07 | tumor | Live | 7 | T2c | low | NO |
| TCGA-EJ-7317-01A-31R-2118-07 | tumor | Live | 7 | T2c | low | NO |
| TCGA-EJ-7331-01A-11R-2118-07 | tumor | Live | 7 | T2c | low | NO |
| TCGA-EJ-7781-01A-11R-2118-07 | tumor | Live | 7 | T2c | low | NO |
| TCGA-EJ-7786-01A-11R-2118-07 | tumor | Live | 7 | T2c | low | NO |
| TCGA-EJ-7791-01A-11R-2118-07 | tumor | Live | 7 | T2c | low | NO |
| TCGA-EJ-7792-01A-11R-2118-07 | tumor | Live | 7 | T2c | low | NO |
| TCGA-EJ-7793-01A-31R-2263-07 | tumor | Live | 7 | T2c | low | NO |
| TCGA-EJ-7794-01A-11R-2118-07 | tumor | Live | 7 | T2c | low | NO |
| TCGA-EJ-7797-01A-11R-2263-07 | tumor | Live | 7 | T2c | low | NO |
| TCGA-EJ-A46H-01A-31R-A26U-07 | tumor | Live | 7 | T2c | low | NO |
| TCGA-EJ-A65M-01A-11R-A29R-07 | tumor | Live | 6 | T2c | low | NO |
| TCGA-EJ-A7NF-01A-11R-A33R-07 | tumor | Live | 7 | T2c | low | NO |
| TCGA-EJ-A7NH-01A-12R-A33R-07 | tumor | Live | 7 | T2c | low | NO |
| TCGA-EJ-A8FN-01A-11R-A352-07 | tumor | Live | 7 | T2c | low | NO |
| TCGA-EJ-AB20-01A-12R-A41O-07 | tumor | Live | 6 | T2c | low | NO |
| TCGA-EJ-AB27-01A-11R-A41O-07 | tumor | Live | 6 | T2c | low | NO |
| TCGA-G9-6329-01A-13R-1965-07 | tumor | Live | 7 | T2c | low | NO |
| TCGA-G9-6347-01A-11R-A31N-07 | tumor | Live | 6 | T2c | low | NO |
| TCGA-G9-6348-01A-11R-1789-07 | tumor | Live | 7 | T2c | low | NO |
| TCGA-G9-6351-01A-21R-1965-07 | tumor | Live | 7 | T2c | low | NO |
| TCGA-G9-6354-01A-11R-A311-07 | tumor | Live | 7 | T2b | low | NO |
| TCGA-G9-6371-01A-11R-1789-07 | tumor | Live | 6 | T2c | low | NO |
| TCGA-G9-6377-01A-11R-1965-07 | tumor | Live | 7 | T2c | low | NO |
| TCGA-G9-6378-01A-11R-1789-07 | tumor | Live | 7 | T2c | low | NO |
| TCGA-G9-6385-01A-11R-1789-07 | tumor | Live | 7 | T2c | low | NO |
| TCGA-G9-7509-01A-11R-A41O-07 | tumor | Live | 6 | T2c | low | NO |
| TCGA-G9-7519-01A-11R-2263-07 | tumor | Live | 7 | T2c | low | NO |
| TCGA-G9-7522-01A-11R-2263-07 | tumor | Live | 7 | T2c | low | NO |
| TCGA-H9-7775-01A-11R-2118-07 | tumor | Live | 7 | T2c | low | NO |
| TCGA-H9-A6BX-01A-31R-A311-07 | tumor | Live | 6 | T2c | low | NO |
| TCGA-HC-7075-01A-11R-1965-07 | tumor | Live | 6 | T2a | low | NO |
| TCGA-HC-7209-01A-11R-2118-07 | tumor | Live | 6 | T2c | low | NO |
| TCGA-HC-7210-01A-11R-2118-07 | tumor | Live | 7 | T2a | low | NO |
| TCGA-HC-7211-01A-11R-2118-07 | tumor | Live | 7 | T2c | low | NO |
| TCGA-HC-7230-01A-11R-2118-07 | tumor | Live | 7 | T2c | low | NO |
| TCGA-HC-7231-01A-11R-2118-07 | tumor | Live | 7 | T2c | low | NO |

| | | | | | | |
|---|---|---|---|---|---|---|
| TCGA-HC-7736-01A-11R-2118-07 | tumor | Live | 7 | T2c | low | NO |
| **Barcode** | **Tissue_type** | **State** | **gleason_sc** | **TNM** | **Risk group** | **Recurrence** |
| TCGA-HC-7737-01A-11R-2118-07 | tumor | Live | 7 | T2c | low | NO |
| TCGA-HC-7740-01A-11R-2118-07 | tumor | Live | 7 | T2c | low | NO |
| TCGA-HC-7740-01B-04R-2302-07 | tumor | Live | 7 | T2c | low | NO |
| TCGA-HC-7740-01B-04R-2302-07 | tumor | Live | 7 | T2c | low | NO |
| TCGA-HC-7747-01A-11R-2118-07 | tumor | Live | 7 | T2c | low | NO |
| TCGA-HC-7748-01A-11R-2118-07 | tumor | Live | 6 | T2a | low | NO |
| TCGA-HC-7750-01A-11R-2118-07 | tumor | Live | 7 | T2c | low | NO |
| TCGA-HC-7818-01A-11R-2118-07 | tumor | Live | 7 | T2c | low | NO |
| TCGA-HC-7820-01A-11R-2118-07 | tumor | Live | 7 | T2c | low | NO |
| TCGA-HC-8213-01A-11R-A29R-07 | tumor | Live | 6 | T2c | low | NO |
| TCGA-HC-8256-01A-11R-2263-07 | tumor | Live | 7 | T2c | low | NO |
| TCGA-HC-8258-01A-11R-2263-07 | tumor | Live | 6 | T2c | low | NO |
| TCGA-HC-8258-01B-05R-2302-07 | tumor | Live | 6 | T2c | low | NO |
| TCGA-HC-8258-01B-05R-2302-07 | tumor | Live | 6 | T2c | low | NO |
| TCGA-HC-8259-01A-11R-2263-07 | tumor | Live | 6 | T2a | low | NO |
| TCGA-HC-8260-01A-11R-2263-07 | tumor | Live | 7 | T2c | low | NO |
| TCGA-HC-8261-01A-11R-2263-07 | tumor | Live | 7 | T2c | low | NO |
| TCGA-HC-A6AL-01A-11R-A30B-07 | tumor | Live | 7 | T2c | low | NO |
| TCGA-HC-A6AP-01A-11R-A30B-07 | tumor | Live | 7 | T2c | low | NO |
| TCGA-HC-A6AQ-01A-11R-A30B-07 | tumor | Live | 7 | T2c | low | NO |
| TCGA-HC-A6AS-01A-11R-A30B-07 | tumor | Live | 7 | T2a | low | NO |
| TCGA-HC-A6HY-01A-11R-A31N-07 | tumor | Live | 7 | T2c | low | NO |
| TCGA-HC-A76X-01A-11R-A33R-07 | tumor | Live | 7 | T2c | low | NO |
| TCGA-HC-A8D0-01A-11R-A36G-07 | tumor | Live | 7 | T2a | low | NO |
| TCGA-HC-A8D1-01A-11R-A36G-07 | tumor | Live | 7 | T2c | low | NO |
| TCGA-HI-7170-01A-11R-2118-07 | tumor | Live | 6 | T2c | low | NO |
| TCGA-J4-8200-01A-11R-A29R-07 | tumor | Live | 7 | T2c | low | NO |
| TCGA-J4-A67O-01A-11R-A30B-07 | tumor | Live | 7 | T2c | low | NO |
| TCGA-J4-A67Q-01A-21R-A30B-07 | tumor | Live | 6 | T2c | low | NO |
| TCGA-J4-A67T-01A-11R-A311-07 | tumor | Live | 7 | T2c | low | NO |
| TCGA-J4-A6M7-01A-11R-A31N-07 | tumor | Live | 7 | T2c | low | NO |
| TCGA-J4-A83J-01A-11R-A36G-07 | tumor | Live | 7 | T2c | low | NO |
| TCGA-J4-A83K-01A-11R-A352-07 | tumor | Live | 6 | T2c | low | NO |
| TCGA-J4-A83L-01A-11R-A352-07 | tumor | Live | 7 | T2c | low | NO |
| TCGA-J4-AATV-01A-11R-A41O-07 | tumor | Live | 6 | T2c | low | NO |
| TCGA-J4-AAU2-01A-11R-A41O-07 | tumor | Live | 6 | T2c | low | NO |
| TCGA-KC-A7F3-01A-21R-A33R-07 | tumor | Live | 7 | T2c | low | NO |
| TCGA-KC-A7F6-01A-11R-A33R-07 | tumor | Live | 7 | T2c | low | NO |
| TCGA-KC-A7FE-01A-12R-A33R-07 | tumor | Live | 7 | T2c | low | NO |
| TCGA-KK-A6DY-01A-12R-A311-07 | tumor | Live | 7 | T2c | low | NO |
| TCGA-KK-A6E2-01A-11R-A311-07 | tumor | Live | 7 | T2c | low | NO |
| TCGA-KK-A6E3-01A-21R-A30B-07 | tumor | Live | 7 | T2c | low | NO |
| TCGA-KK-A6E4-01A-11R-A30B-07 | tumor | Live | 7 | T2b | low | NO |

| | | | | | | |
|---|---|---|---|---|---|---|
| TCGA-KK-A7AV-01A-11R-A32O-07 | tumor | Live | 7 | T2c | low | NO |
| **Barcode** | **Tissue_type** | **State** | **gleason_sc** | **TNM** | **Risk group** | **Recurrence** |
| TCGA-KK-A8I5-01A-11R-A36G-07 | tumor | Live | 7 | T2b | low | NO |
| TCGA-M7-A71Y-01A-22R-A32O-07 | tumor | Live | 7 | T2c | low | NO |
| TCGA-M7-A720-01A-12R-A32O-07 | tumor | Live | 6 | T2c | low | NO |
| TCGA-M7-A721-01A-12R-A32O-07 | tumor | Live | 7 | T2c | low | NO |
| TCGA-QU-A6IL-01A-11R-A31N-07 | tumor | Live | 7 | T2c | low | NO |
| TCGA-QU-A6IM-01A-11R-A31N-07 | tumor | Live | 7 | T2a | low | NO |
| TCGA-QU-A6IN-01A-11R-A31N-07 | tumor | Live | 7 | T2c | low | NO |
| TCGA-QU-A6IO-01A-11R-A31N-07 | tumor | Live | 6 | T2c | low | NO |
| TCGA-QU-A6IP-01A-11R-A31N-07 | tumor | Live | 6 | T2a | low | NO |
| TCGA-V1-A8MK-01A-11R-A36G-07 | tumor | Live | 6 | T2c | low | NO |
| TCGA-V1-A8ML-01A-11R-A37L-07 | tumor | Live | 7 | T2c | low | NO |
| TCGA-V1-A8WN-01A-11R-A37L-07 | tumor | Live | 6 | T2c | low | NO |
| TCGA-V1-A8X3-01A-11R-A37L-07 | tumor | Live | 7 | T2c | low | NO |
| TCGA-V1-A9OF-01A-11R-A41O-07 | tumor | Live | 6 | T2c | low | NO |
| TCGA-VN-A88L-01A-11R-A352-07 | tumor | Live | 7 | T2c | low | NO |
| TCGA-VN-A88M-01A-11R-A352-07 | tumor | Live | 7 | T2c | low | NO |
| TCGA-VN-A88O-01A-11R-A352-07 | tumor | Live | 7 | T2c | low | NO |
| TCGA-VN-A88P-01A-11R-A352-07 | tumor | Live | 7 | T2c | low | NO |
| TCGA-VP-A87E-01A-31R-A352-07 | tumor | Live | 6 | T2c | low | NO |
| TCGA-X4-A8KS-01A-12R-A36G-07 | tumor | Live | 7 | T2c | low | NO |
| TCGA-XJ-A83H-01A-11R-A352-07 | tumor | Live | 7 | T2c | low | NO |
| TCGA-XJ-A9DQ-01A-11R-A37L-07 | tumor | Live | 6 | T2a | low | NO |
| TCGA-ZG-A8QX-01A-11R-A37L-07 | tumor | Live | 6 | T2c | low | NO |
| TCGA-EJ-A6RC-01A-11R-A32O-07 | tumor | Live | 7 | [Not Available] | | NO |
| TCGA-G9-6336-01A-11R-1789-07 | tumor | Live | 7 | [Not Available] | | NO |
| TCGA-TK-A8OK-01A-22R-A36G-07 | tumor | Live | 7 | [Not Available] | | NO |
| TCGA-CH-5753-01A-11R-1580-07 | tumor | Live | 9 | T3b | high | NOT AVAILABLE |
| TCGA-CH-5754-01A-11R-1580-07 | tumor | Live | 9 | T3b | high | NOT AVAILABLE |
| TCGA-CH-5761-01A-11R-1580-07 | tumor | Live | 9 | T3b | high | NOT AVAILABLE |
| TCGA-CH-5764-01A-21R-1580-07 | tumor | Live | 7 | T3b | high | NOT AVAILABLE |
| TCGA-CH-5769-01A-11R-1580-07 | tumor | Live | 9 | T3b | high | NOT AVAILABLE |
| TCGA-CH-5788-01A-11R-1580-07 | tumor | Live | 7 | T3b | high | NOT AVAILABLE |
| TCGA-CH-5792-01A-11R-1580-07 | tumor | Live | 9 | T3a | high | NOT AVAILABLE |
| TCGA-G9-6366-01A-11R-2118-07 | tumor | Live | 7 | T3b | high | NOT AVAILABLE |
| TCGA-HC-A8CY-01A-11R-A36G-07 | tumor | Live | 9 | T3a | high | NOT AVAILABLE |
| TCGA-KK-A7AP-01A-12R-A33R-07 | tumor | Live | 9 | T3b | high | NOT AVAILABLE |
| TCGA-KK-A8IB-01A-11R-A36G-07 | tumor | Live | 9 | T3a | high | NOT AVAILABLE |
| TCGA-KK-A8IL-01A-11R-A36G-07 | tumor | Live | 9 | T3b | high | NOT AVAILABLE |
| TCGA-YJ-A8SW-01A-11R-A37L-07 | tumor | Live | 9 | T3b | high | NOT AVAILABLE |
| TCGA-CH-5737-01A-11R-1580-07 | tumor | Live | 7 | T2c | intermediate | NOT AVAILABLE |
| TCGA-CH-5744-01A-11R-1580-07 | tumor | Live | 7 | T2c | intermediate | NOT AVAILABLE |
| TCGA-CH-5745-01A-11R-1580-07 | tumor | Live | 7 | T3a | intermediate | NOT AVAILABLE |
| TCGA-CH-5748-01A-11R-1580-07 | tumor | Live | 7 | T3a | intermediate | NOT AVAILABLE |

| | | | | | | |
|---|---|---|---|---|---|---|
| TCGA-CH-5766-01A-11R-1580-07 | tumor | Live | 7 | T3a | intermediate | NOT AVAILABLE |
| **Barcode** | **Tissue_type** | **State** | **gleason_sc** | **TNM** | **Risk group** | **Recurrence** |
| TCGA-EJ-7330-01A-11R-2118-07 | tumor | Live | 7 | T3a | intermediate | NOT AVAILABLE |
| TCGA-G9-6343-01A-21R-1965-07 | tumor | Live | 7 | T2c | intermediate | NOT AVAILABLE |
| TCGA-KK-A7AW-01A-11R-A32O-07 | tumor | Live | 7 | T3a | intermediate | NOT AVAILABLE |
| TCGA-KK-A7AZ-01A-12R-A32O-07 | tumor | Live | 7 | T3a | intermediate | NOT AVAILABLE |
| TCGA-KK-A7B1-01A-11R-A32O-07 | tumor | Live | 7 | T3a | intermediate | NOT AVAILABLE |
| TCGA-CH-5740-01A-11R-1580-07 | tumor | Live | 7 | T2c | low | NOT AVAILABLE |
| TCGA-HC-7752-01A-11R-2118-07 | tumor | Live | 7 | T2c | low | NOT AVAILABLE |
| TCGA-KC-A4BN-01A-61R-A250-07 | tumor | Live | 7 | T2a | low | NOT AVAILABLE |
| TCGA-2A-A8W3-01A-11R-A37L-07 | tumor | Live | 9 | T3a | high | YES |
| TCGA-CH-5751-01A-11R-1580-07 | tumor | Live | 10 | T4 | high | YES |
| TCGA-EJ-5518-01A-01R-1580-07 | tumor | Live | 9 | T4 | high | YES |
| TCGA-EJ-5524-01A-01R-1580-07 | tumor | Live | 9 | T3a | high | YES |
| TCGA-EJ-5525-01A-01R-1580-07 | tumor | Live | 9 | T2c | high | YES |
| TCGA-EJ-5526-01A-01R-1580-07 | tumor | Live | 8 | T3a | high | YES |
| TCGA-EJ-8472-01A-11R-2403-07 | tumor | Live | 8 | T3a | high | YES |
| TCGA-EJ-A46F-01A-31R-A250-07 | tumor | Live | 8 | T3b | high | YES |
| TCGA-EJ-A65F-01A-21R-A311-07 | tumor | Live | 9 | T3b | high | YES |
| TCGA-EJ-A6RA-01A-11R-A33R-07 | tumor | Live | 8 | T3b | high | YES |
| TCGA-EJ-A7NN-01A-11R-A33R-07 | tumor | Live | 7 | T3b | high | YES |
| TCGA-EJ-A8FP-01A-21R-A36G-07 | tumor | Live | 8 | T2c | high | YES |
| TCGA-G9-6339-01A-12R-A311-07 | tumor | Live | 7 | T3b | high | YES |
| TCGA-G9-A9S0-01A-11R-A41O-07 | tumor | Live | 8 | T3b | high | YES |
| TCGA-HC-7213-01A-11R-2118-07 | tumor | Live | 9 | T3b | high | YES |
| TCGA-HC-7232-01A-11R-2118-07 | tumor | Live | 9 | T3b | high | YES |
| TCGA-HC-7821-01A-12R-2118-07 | tumor | Live | 8 | T3b | high | YES |
| TCGA-HC-A9TE-01A-11R-A41O-07 | tumor | Live | 9 | T3b | high | YES |
| TCGA-HC-A9TH-01A-11R-A41O-07 | tumor | Live | 9 | T3b | high | YES |
| TCGA-HI-7168-01A-11R-2118-07 | tumor | Live | 9 | T3b | high | YES |
| TCGA-HI-7171-01A-12R-2118-07 | tumor | Live | 9 | T3b | high | YES |
| TCGA-J4-A6G3-01A-11R-A311-07 | tumor | Live | 8 | T3b | high | YES |
| TCGA-J4-AATZ-01A-11R-A41O-07 | tumor | Live | 9 | T3a | high | YES |
| TCGA-J9-A52B-01A-11R-A26U-07 | tumor | Live | 9 | T3b | high | YES |
| TCGA-J9-A52E-01A-11R-A26U-07 | tumor | Live | 9 | T3b | high | YES |
| TCGA-J9-A8CL-01A-11R-A352-07 | tumor | Live | 9 | T3b | high | YES |
| TCGA-J9-A8CM-01A-11R-A352-07 | tumor | Live | 9 | T3b | high | YES |
| TCGA-KC-A4BR-01A-32R-A32Y-07 | tumor | Live | 9 | T3b | high | YES |
| TCGA-KC-A4BV-01A-31R-A26U-07 | tumor | Live | 9 | T3a | high | YES |
| TCGA-KK-A59X-01A-11R-A29R-07 | tumor | Live | 9 | T3b | high | YES |
| TCGA-KK-A5A1-01A-11R-A29R-07 | tumor | Live | 9 | T3b | high | YES |
| TCGA-KK-A6E0-01A-11R-A311-07 | tumor | Live | 9 | T3a | high | YES |
| TCGA-KK-A6E7-01A-11R-A31N-07 | tumor | Live | 9 | T3a | high | YES |
| TCGA-KK-A7AU-01A-11R-A32O-07 | tumor | Live | 9 | T3b | high | YES |
| TCGA-KK-A7B0-01A-11R-A32O-07 | tumor | Live | 9 | T3b | high | YES |

| | | | | | | |
|---|---|---|---|---|---|---|
| TCGA-KK-A7B2-01A-12R-A32O-07 | tumor | Live | 9 | T3a | high | YES |
| **Barcode** | **Tissue_type** | **State** | **gleason_sc** | **TNM** | **Risk group** | **Recurrence** |
| TCGA-KK-A7B3-01A-11R-A33R-07 | tumor | Live | 9 | T3b | high | YES |
| TCGA-KK-A7B4-01A-11R-A32O-07 | tumor | Live | 9 | T3b | high | YES |
| TCGA-KK-A8I4-01A-11R-A36G-07 | tumor | Live | 7 | T3b | high | YES |
| TCGA-KK-A8I7-01A-21R-A36G-07 | tumor | Live | 9 | T3b | high | YES |
| TCGA-KK-A8I9-01A-11R-A36G-07 | tumor | Live | 8 | T3a | high | YES |
| TCGA-KK-A8IC-01A-11R-A36G-07 | tumor | Live | 9 | T3b | high | YES |
| TCGA-KK-A8II-01A-11R-A36G-07 | tumor | Live | 9 | T3a | high | YES |
| TCGA-KK-A8IJ-01A-11R-A352-07 | tumor | Live | 7 | T3b | high | YES |
| TCGA-M7-A722-01A-12R-A36G-07 | tumor | Live | 8 | T3a | high | YES |
| TCGA-M7-A724-01A-12R-A32O-07 | tumor | Live | 8 | T3a | high | YES |
| TCGA-V1-A8WS-01A-11R-A37L-07 | tumor | Live | 6 | T3b | high | YES |
| TCGA-V1-A8WW-01A-11R-A37L-07 | tumor | Live | 9 | T3b | high | YES |
| TCGA-V1-A9O5-01A-11R-A41O-07 | tumor | Live | 9 | T4 | high | YES |
| TCGA-V1-A9O7-01A-21R-A41O-07 | tumor | Live | 9 | T3a | high | YES |
| TCGA-V1-A9OL-01A-11R-A41O-07 | tumor | Live | 9 | T3b | high | YES |
| TCGA-V1-A9Z7-01A-11R-A41O-07 | tumor | Live | 9 | T3b | high | YES |
| TCGA-V1-A9ZI-01A-11R-A41O-07 | tumor | Live | 9 | T3b | high | YES |
| TCGA-V1-A9ZR-01A-11R-A41O-07 | tumor | Live | 8 | T4 | high | YES |
| TCGA-VN-A88R-01A-11R-A36G-07 | tumor | Live | 8 | T2c | high | YES |
| TCGA-VP-A878-01A-31R-A352-07 | tumor | Live | 9 | T2c | high | YES |
| TCGA-VP-A87B-01A-11R-A352-07 | tumor | Live | 8 | T2b | high | YES |
| TCGA-VP-A87D-01A-11R-A352-07 | tumor | Live | 9 | T3a | high | YES |
| TCGA-VP-A87K-01A-11R-A352-07 | tumor | Live | 8 | T3b | high | YES |
| TCGA-X4-A8KQ-01A-12R-A36G-07 | tumor | Live | 9 | T3b | high | YES |
| TCGA-XJ-A9DX-01A-11R-A37L-07 | tumor | Live | 9 | T3b | high | YES |
| TCGA-XK-AAIW-01A-11R-A41O-07 | tumor | Live | 9 | T3b | high | YES |
| TCGA-YL-A8HK-01A-11R-A36G-07 | tumor | Live | 9 | T3b | high | YES |
| TCGA-YL-A8HM-01A-11R-A36G-07 | tumor | Live | 9 | T3b | high | YES |
| TCGA-YL-A8S8-01A-11R-A37L-07 | tumor | Live | 9 | T3a | high | YES |
| TCGA-YL-A8S9-01A-11R-A37L-07 | tumor | Live | 9 | T3a | high | YES |
| TCGA-YL-A8SB-01A-31R-A37L-07 | tumor | Live | 9 | T3a | high | YES |
| TCGA-YL-A8SC-01A-11R-A37L-07 | tumor | Live | 9 | T3a | high | YES |
| TCGA-YL-A8SI-01A-11R-A41O-07 | tumor | Live | 9 | T3a | high | YES |
| TCGA-YL-A8SJ-01B-11R-A37L-07 | tumor | Live | 9 | T3b | high | YES |
| TCGA-YL-A8SP-01B-11R-A37L-07 | tumor | Live | 9 | T3b | high | YES |
| TCGA-YL-A8SQ-01B-11R-A37L-07 | tumor | Live | 9 | T3b | high | YES |
| TCGA-YL-A9WJ-01A-11R-A37L-07 | tumor | Live | 8 | T3b | high | YES |
| TCGA-YL-A9WK-01A-11R-A37L-07 | tumor | Live | 9 | T3b | high | YES |
| TCGA-YL-A9WL-01A-11R-A41O-07 | tumor | Live | 9 | T3a | high | YES |
| TCGA-YL-A9WX-01A-21R-A41O-07 | tumor | Live | 9 | T3b | high | YES |
| TCGA-YL-A9WY-01A-11R-A41O-07 | tumor | Live | 9 | T3b | high | YES |
| TCGA-ZG-A9L2-01A-31R-A41O-07 | tumor | Live | 9 | T3b | high | YES |
| TCGA-ZG-A9L6-01A-11R-A41O-07 | tumor | Live | 9 | T3b | high | YES |

| | | | | | | |
|---|---|---|---|---|---|---|
| TCGA-ZG-A9L9-01A-11R-A41O-07 | tumor | Live | 9 | T3b | high | YES |
| **Barcode** | **Tissue_type** | **State** | **gleason_sc** | **TNM** | **Risk group** | **Recurrence** |
| TCGA-ZG-A9LZ-01A-11R-A41O-07 | tumor | Live | 9 | T3b | high | YES |
| TCGA-CH-5791-01A-11R-1580-07 | tumor | Live | 7 | T3a | intermediate | YES |
| TCGA-EJ-5504-01A-01R-1580-07 | tumor | Live | 7 | T3a | intermediate | YES |
| TCGA-EJ-7318-01B-11R-A32O-07 | tumor | Live | 7 | T3a | intermediate | YES |
| TCGA-EJ-A8FS-01A-11R-A352-07 | tumor | Live | 7 | T3a | intermediate | YES |
| TCGA-G9-6332-01A-11R-1789-07 | tumor | Live | 7 | T3a | intermediate | YES |
| TCGA-G9-6362-01A-11R-1789-07 | tumor | Live | 7 | T3a | intermediate | YES |
| TCGA-G9-6498-01A-12R-A311-07 | tumor | Live | 7 | T3a | intermediate | YES |
| TCGA-HC-7079-01A-11R-1965-07 | tumor | Live | 7 | T3a | intermediate | YES |
| TCGA-HC-7742-01A-11R-2118-07 | tumor | Live | 7 | T3a | intermediate | YES |
| TCGA-J4-A67N-01A-11R-A30B-07 | tumor | Live | 7 | T3a | intermediate | YES |
| TCGA-J4-A67S-01A-11R-A30B-07 | tumor | Live | 7 | T3a | intermediate | YES |
| TCGA-J4-A83M-01A-11R-A352-07 | tumor | Live | 7 | T2c | intermediate | YES |
| TCGA-KK-A7AQ-01A-11R-A33R-07 | tumor | Live | 7 | T2c | intermediate | YES |
| TCGA-KK-A7AY-01A-11R-A33R-07 | tumor | Live | 7 | T2c | intermediate | YES |
| TCGA-KK-A8IF-01A-11R-A36G-07 | tumor | Live | 7 | T3a | intermediate | YES |
| TCGA-V1-A8MJ-01A-11R-A36G-07 | tumor | Live | 7 | T3a | intermediate | YES |
| TCGA-V1-A8MM-01A-11R-A37L-07 | tumor | Live | 7 | T3a | intermediate | YES |
| TCGA-V1-A8MU-01A-11R-A37L-07 | tumor | Live | 7 | T3a | intermediate | YES |
| TCGA-XK-AAJR-01A-11R-A41O-07 | tumor | Live | 7 | T3a | intermediate | YES |
| TCGA-YL-A8HO-01A-11R-A36G-07 | tumor | Live | 7 | T3a | intermediate | YES |
| TCGA-CH-5743-01A-21R-1580-07 | tumor | Live | 7 | T2c | low | YES |
| TCGA-HC-7080-01A-11R-1965-07 | tumor | Live | 7 | T2a | low | YES |
| TCGA-HC-7212-01A-11R-2118-07 | tumor | Live | 7 | T2a | low | YES |
| TCGA-HC-7738-01A-11R-2118-07 | tumor | Live | 7 | T2c | low | YES |
| TCGA-J4-A83N-01A-11R-A352-07 | tumor | Live | 7 | T2c | low | YES |
| TCGA-KC-A4BL-01A-31R-A250-07 | tumor | Live | 7 | T2c | low | YES |
| TCGA-V1-A9OT-01A-11R-A41O-07 | tumor | Live | 6 | T2c | low | YES |
| TCGA-HC-8261-01B-05R-2302-07 | tumor | Live | | | | |
| TCGA-HC-8261-01B-05R-2302-07 | tumor | Live | | | | |
