## Supplemental Table S8 for "Blind exploration of the unreferenced transcriptome reveals novel RNAs for prostate cancer diagnosis"

**Table S8. PCA3 and k-mer contigs expression quantification assessed by the poly(A)+ unstranded RNA-seq in 557 prostate specimens from the TCGA-PRAD cohort (Validation Set).**

| Probe_ID | Contig_ID | mean_normal | mean_tumor | log2FC | wilcoxon_pvalue |
|---|---|---|---|---|---|
| P16 | ctg_111348 | 0,033317712 | 0,276915399 | 3,055084053 | 7,74E-19 |
| P1 | ctg_17297 | 0,100628353 | 0,456800544 | 2,182527512 | 7,11E-18 |
| PCA3 | - | 0,035183696 | 0,364625454 | 3,373436337 | 8,68E-15 |
| P19 | ctg_61472 | 0,001842814 | 0,039351168 | 4,416424112 | 1,69E-14 |
| P2 | ctg_28650 | 0,003189305 | 0,032253596 | 3,33814625 | 3,15E-14 |
| P18 | ctg_105149 | 0,006071471 | 0,070266607 | 3,532721288 | 5,64E-14 |
| P7 | ctg_117356 | 0,004177631 | 0,048450817 | 3,535764171 | 7,44E-13 |
| P13 | ctg_37852 | 0,080243954 | 0,324725284 | 2,016755112 | 3,52E-11 |
| P6 | ctg_111158 | 0,001401421 | 0,016630154 | 3,568838674 | 7,98E-10 |
| P10 | ctg_25348 | 0,0088826 | 0,036679258 | 2,045910463 | 1,14E-09 |
| P20 | ctg_44030 | 0,006500788 | 0,036907249 | 2,505217713 | 1,23E-09 |
| P15 | ctg_512 | 0,000227631 | 0,007715559 | 5,082999012 | 1,57E-06 |
| P11 | ctg_104447 | 0,00047629 | 0,009275747 | 4,283552311 | 3,17E-05 |
| P23 | ctg_29077 | 0,000520086 | 0,00800317 | 3,943749061 | 3,36E-04 |
| P21 | ctg_23999 | 0,000241552 | 0,004893751 | 4,340534219 | 4,34E-04 |
| P9 | ctg_9446 | 0,002633435 | 0,008302558 | 1,656610093 | 1,25E-03 |
| P22 | ctg_119680 | 0,002148707 | 0,006930616 | 1,689515145 | 1,50E-03 |
| P14 | ctg_61528 | 0,00032437 | 0,002174545 | 2,745001851 | 1,21E-02 |
| P17 | ctg_36195 | 0 | 0,001401064 | Inf | 3,33E-02 |
| P3 | ctg_57223 | 0,002585139 | 0,003604376 | 0,479507505 | 1,10E-01 |
| P4 | ctg_63866 | 0,000520086 | 0,001390124 | 1,418391574 | 1,19E-01 |
| P12 | ctg_2815 | 0 | 0,000251205 | Inf | 1,81E-01 |
| P5 | ctg_123090 | 0 | 0,000176244 | Inf | 2,37E-01 |
| P8 | ctg_73782 | 0 | 0,000136565 | Inf | 2,37E-01 |
| GAPDH | - | 17,62287189 | 20,93793405 | 0,248670048 | 1,41E-02 |
| RPL11 | - | 27,72614405 | 40,83156891 | 0,558438012 | 8,72E-09 |
| ZNF2 | - | 0,031923613 | 0,024424479 | -0,386296175 | 9,89E-01 |
| GPATCH3 | - | 0,11955761 | 0,140679766 | 0,234708875 | 2,68E-02 |
| NOL7 | - | 0,689786984 | 0,69734313 | 0,015717807 | 2,60E-01 |
| ZNF346 | - | 0,048759098 | 0,042720572 | -0,190740477 | 9,47E-01 |
