## Supplemental Table S9 for "Blind exploration of the unreferenced transcriptome reveals novel RNAs for prostate cancer diagnosis"

**Table S9. Mean and Fold Change expression of PCA3, DE-kupl contigs and housekeeping genes in low-risk (LR) and high-risk (HR) tumors and recurrence negative (NO) and positive (YES) specimens of the PAIR cohort (Selection Set).**

| probe_ID | contig_ID | mean_LR | mean_HR | log2FC | FC, HRvsLR | wilcoxon_pvalue | mean_NO | mean_YES | log2FC | FC, YESvsNO | wilcoxon_pvalue |
|---|---|---|---|---|---|---|---|---|---|---|---|
| P2 | ctg_28650 | 0,73 | 1,31 | 0,84 | 1,78 | 6,08E-05 | 0,88 | 1,28 | 0,55 | 1,46 | 6,00E-04 |
| P12 | ctg_2815 | 1,54 | 1,83 | 0,25 | 1,19 | 1,81E-01 | 1,41 | 1,90 | 0,43 | 1,35 | 1,18E-02 |
| P21 | ctg_23999 | 1,86 | 1,81 | -0,04 | 0,97 | 8,73E-01 | 1,56 | 2,05 | 0,40 | 1,32 | 4,46E-01 |
| P18 | ctg_105149 | 1,41 | 1,68 | 0,25 | 1,19 | 5,61E-01 | 1,36 | 1,74 | 0,36 | 1,28 | 3,11E-01 |
| P6 | ctg_111158 | 1,50 | 2,66 | 0,82 | 1,77 | 2,44E-05 | 1,95 | 2,50 | 0,35 | 1,28 | 1,49E-03 |
| P10 | ctg_25348 | 0,49 | 0,67 | 0,45 | 1,36 | 2,59E-02 | 0,52 | 0,64 | 0,31 | 1,24 | 1,02E-01 |
| P1 | ctg_17297 | 5,47 | 7,90 | 0,53 | 1,44 | 1,20E-01 | 6,60 | 7,84 | 0,25 | 1,19 | 2,41E-01 |
| P5 | ctg_123090 | 0,38 | 0,37 | -0,04 | 0,97 | 6,14E-01 | 0,36 | 0,41 | 0,20 | 1,15 | 1,58E-01 |
| P14 | ctg_61528 | 1,54 | 1,46 | -0,08 | 0,95 | 8,75E-01 | 1,35 | 1,55 | 0,20 | 1,15 | 4,64E-01 |
| P16 | ctg_111348 | 1,65 | 2,53 | 0,61 | 1,53 | 2,33E-02 | 2,05 | 2,30 | 0,16 | 1,12 | 2,41E-01 |
| P17 | ctg_36195 | 0,26 | 0,26 | 0,04 | 1,03 | 1,49E-01 | 0,23 | 0,25 | 0,11 | 1,08 | 1,08E-01 |
| P7 | ctg_117356 | 3,78 | 4,19 | 0,15 | 1,11 | 7,42E-01 | 4,01 | 4,10 | 0,03 | 1,02 | 7,39E-01 |
| P9 | ctg_9446 | 1,93 | 1,78 | -0,12 | 0,92 | 8,17E-01 | 1,92 | 1,92 | 0,00 | 1,00 | 2,98E-01 |
| P19 | ctg_61472 | 2,06 | 2,35 | 0,19 | 1,14 | 6,07E-01 | 2,28 | 2,25 | -0,02 | 0,99 | 7,96E-01 |
| P4 | ctg_63866 | 1,09 | 0,99 | -0,14 | 0,91 | 9,86E-01 | 1,19 | 1,12 | -0,08 | 0,95 | 6,96E-01 |
| P15 | ctg_512 | 1,16 | 1,12 | -0,05 | 0,97 | 5,07E-01 | 1,14 | 1,07 | -0,08 | 0,94 | 7,40E-01 |
| P20 | ctg_44030 | 4,98 | 3,87 | -0,37 | 0,78 | 9,68E-01 | 4,27 | 4,03 | -0,08 | 0,94 | 7,66E-01 |
| P23 | ctg_29077 | 1,73 | 1,62 | -0,10 | 0,93 | 4,82E-01 | 1,67 | 1,49 | -0,16 | 0,89 | 6,67E-01 |
| P22 | ctg_119680 | 0,21 | 0,17 | -0,25 | 0,84 | 8,67E-01 | 0,19 | 0,17 | -0,16 | 0,89 | 8,43E-01 |
| P3 | ctg_57223 | 1,57 | 1,39 | -0,18 | 0,89 | 5,00E-01 | 1,63 | 1,42 | -0,20 | 0,87 | 4,90E-01 |
| P13 | ctg_37852 | 7,08 | 4,90 | -0,53 | 0,69 | 9,46E-01 | 6,31 | 5,30 | -0,25 | 0,84 | 9,74E-01 |
| P11 | ctg_104447 | 1,92 | 0,96 | -1,00 | 0,50 | 9,98E-01 | 1,55 | 1,19 | -0,38 | 0,77 | 9,62E-01 |
| P8 | ctg_73782 | 3,23 | 2,00 | -0,69 | 0,62 | 1,00E+00 | 3,42 | 2,47 | -0,47 | 0,72 | 9,97E-01 |
| PCA3 | - | 18,00 | 14,54 | -0,31 | 0,81 | 9,73E-01 | 17,19 | 13,50 | -0,35 | 0,79 | 9,93E-01 |
| GAPDH | - | 171,61 | 202,58 | 0,24 | 1,18 | 3,48E-03 | 172,42 | 196,58 | 0,19 | 1,14 | 1,02E-02 |
| GPATCH3 | - | 1,19 | 1,22 | 0,03 | 1,02 | 2,24E-01 | 1,23 | 1,20 | -0,03 | 0,98 | 7,37E-01 |
| NOL7 | - | 15,78 | 16,93 | 0,10 | 1,07 | 4,58E-02 | 16,50 | 16,06 | -0,04 | 0,97 | 6,43E-01 |
| RPL11 | - | 274,85 | 272,74 | -0,01 | 0,99 | 6,64E-01 | 273,78 | 254,03 | -0,11 | 0,93 | 7,85E-01 |
| ZNF2 | - | 1,00 | 0,99 | -0,02 | 0,99 | 6,58E-01 | 0,98 | 1,00 | 0,03 | 1,02 | 2,98E-01 |
| ZNF346 | - | 0,80 | 0,79 | -0,03 | 0,98 | 5,82E-01 | 0,79 | 0,80 | 0,02 | 1,01 | 2,43E-01 |
